# Spatially Resolved Microglial State Transitions Govern Strain-Specific Zika Neuropathogenesis

**DOI:** 10.64898/2026.04.30.721456

**Authors:** Md Musaddaqul Hasib, Wen Meng, Gonzalo Mena, Hugh Halloway, Suzane Ramos da Silva, Gary Kohanbash, Yufei Huang, Shou-Jiang Gao

## Abstract

Neurotropic viruses disrupt brain homeostasis through complex interactions among infected cells, resident immune responses, and tissue architecture, yet how these processes unfold across space, time, and cell types, and how viral strain differences shape disease severity, remains poorly understood. Here, we integrate high-resolution spatial transcriptomics with infection-aware cell-type profiling to construct a spatiotemporal atlas of Zika virus (ZIKV) infection in the mouse brain. Comparing Asian and African ZIKV strains across early and late infection stages, we uncover structured, strain-dependent reorganization of immune and structural cell populations that defines discrete infection-associated tissue niches. Microglia undergo region-specific state transitions characterized by both cell-intrinsic antiviral programs and widespread bystander activation, producing tissue-wide immune amplification. In Asian strain infection, we identify disease-associated microglia (DAM) as critical mediators of infection containment: DAM accumulate in regions where viral burden stabilizes, are promoted by Apoe-Trem2 signaling from infected cells and are governed by transcription regulators that restrain inflammatory programs while preserving phagocytic and antiviral functions. In contrast, African strain infection is marked by impaired Apoe-Trem2 signaling, persistent inflammatory microglial activation, and failure of containment. Progressive infection leads to depletion of oligodendrocytes, astrocytes, and neurons, loss of local cellular diversity and disruption of tissue architecture concentrated in somatosensory and motor regions associated with myelination and synaptic programs. These architectural disruptions correlate with severe neurological phenotypes in African strain infection and are preceded by transcriptional dysregulation in infected glial cells, including sustained stress responses, inflammatory signaling, and suppression of myelination and homeostatic pathways. Together, our study establishes a spatially resolved framework linking viral strain-specific microglial states to tissue disorganization and neurological functional impairment, providing mechanistic insight into how neurotropic viruses reshape microenvironments to drive neurological disease.

## INTRODUCTION

Neurotropic viral infections pose a major threat to central nervous system (CNS) integrity, frequently causing long-lasting neurological deficits or death. Viruses such as Zika virus (ZIKV), West Nile virus, and herpesviruses can invade the brain and trigger inflammatory responses that disrupt neuronal circuits, glial support functions, and tissue architecture [1, 2]. Despite extensive study of viral tropism and immune activation, a fundamental gap persists in understanding how infection reorganizes the brain microenvironment across space, time, and cell types, and how this reorganization gives rise to divergent disease outcomes.

ZIKV offers a powerful model to address these questions. Distinct viral strains exhibit strikingly different neuropathological consequences, ranging from mild or transient symptoms to severe neuroinflammation, motor dysfunction, and lethality [3]. While African ZIKV strains are associated with higher virulence and neurotoxicity in experimental models, Asian strains often produce comparatively milder disease, despite sharing core viral features [4]. The cellular and spatial mechanisms underlying these strain-specific outcomes remain poorly understood.

Central to the brain’s response to viral challenge is microglia, the resident immune cells of the CNS. Upon infection, microglia rapidly transition from homeostatic to activated states, producing cytokines, phagocytosing infected cells, and shaping immune cell recruitment [5]. Critically, microglial activation is not monolithic: emerging evidence indicates that microglial responses are highly context-dependent, shaped by local tissue environment, infection burden, and interactions with neighboring cells [6]. Whether microglial responses to viral infection are primarily cell-intrinsic, restricted to infected cells, or propagate broadly to uninfected bystander populations across the brain remains an open question with important implications for understanding neuroinflammatory pathology.

Beyond immune activation, viral infections profoundly affect structural cell types that maintain brain function. Oligodendrocytes, astrocytes, and neurons collectively support myelination, metabolic support, synaptic regulation, and maintenance of extracellular homeostasis [7]. Disruption of these functions can impair neural connectivity and circuit stability, even in the absence of overt neuronal death. Yet infection-induced changes in cell-type composition and spatial organization, factors that may critically influence tissue resilience or vulnerability during infection, have been difficult to quantify in vivo.

Recent advances in spatial transcriptomics enable simultaneous measurement of gene expression, cell-type identity, and spatial context within intact tissue sections [8]. These technologies provide unprecedented opportunities to move beyond bulk or dissociated analyses and interrogate how infection reshapes cellular neighborhoods, tissue niches, and functional states across the brain. However, spatially resolved studies of viral encephalitis remain limited, and few have systematically linked spatial cellular reorganization to strain-specific disease severity.

Here, we leverage high-resolution spatial transcriptomics combined with infection-aware cell-type annotation to construct a comprehensive atlas of ZIKV infection in the mouse brain. By profiling multiple brain regions across early and late infection stages, and directly comparing Asian and African ZIKV strains, we dissect how infection alters cellular composition, spatial organization, and functional programs at a single cell resolution. We identify distinct infection-associated tissue niches defined by coordinated changes in immune and structural cell densities, reveal disease-associated microglia, driven by Apoe-Trem2 signaling and specific transcriptional regulators, are central to infection containment, and uncover progressive loss of cellular diversity in regions critical for motor and sensory function. We further demonstrate that transcriptional dysregulation in infected glial cells, marked by sustained stress and inflammatory signaling and suppression of myelination and homeostatic pathways, precedes and accompanies structural disorganization. Our findings establish a spatial and cellular framework linking viral strain-specific immune responses to tissue-level disruption and neurological dysfunction. By integrating infection status, cell-type-specific transcriptional programs, and spatial microenvironmental context, this study provides mechanistic insight into how neurotropic viruses remodel the brain and highlights cellular pathways that may underlie severe neurological disease.

## RESULTS

### Integrative spatial and single-cell transcriptomics reveal strain- and time-dependent cellular and infection dynamics in ZIKV-infected mouse brains

We inoculated African or Asian strains of ZIKV by direct injection into brains. Mice inoculated with Asian ZIKV strain had minimal pathological effect while those inoculated with African ZIKA strain manifested neurological disorders starting at day 4 and further exacerbated over time. By day 7 after inoculation, all mice inoculated with African ZIKA strain were dead while all mice inoculated with Asian ZIKV strain survived (Fig. 1S1A&B). These results of more virulent nature of the African strain than Asian strain results are consistent with those previously reported [9, 10].

To comprehensively map ZIKV infection dynamics in the developing mouse brain, we generated integrated spatial transcriptomics and single-cell RNA sequencing (scRNA-seq) data from brains infected with African or Asian ZIKV strains at 4- and 6-days post-infection (dpi), alongside mock-infected controls (Fig. 1A). After stringent quality control (see Methods), we obtained approximately 750K spatially resolved single cells with transcriptomic profiles for 218 genes and approximately 400K single cells with whole transcriptomes (Fig. 1S2). The number of spatially resolved cells per sample ranged from 20,066 to 55,387 cells per brain (Fig. 1S2A). For scRNA-seq) (Fig. 1S2B, we captured 25,219 to 73,554 cells per sample with average 515 to 1,098 genes expressed per cell. These differences reflect the trade-off between spatial resolution and transcriptomic depth.

**Figure 1.**
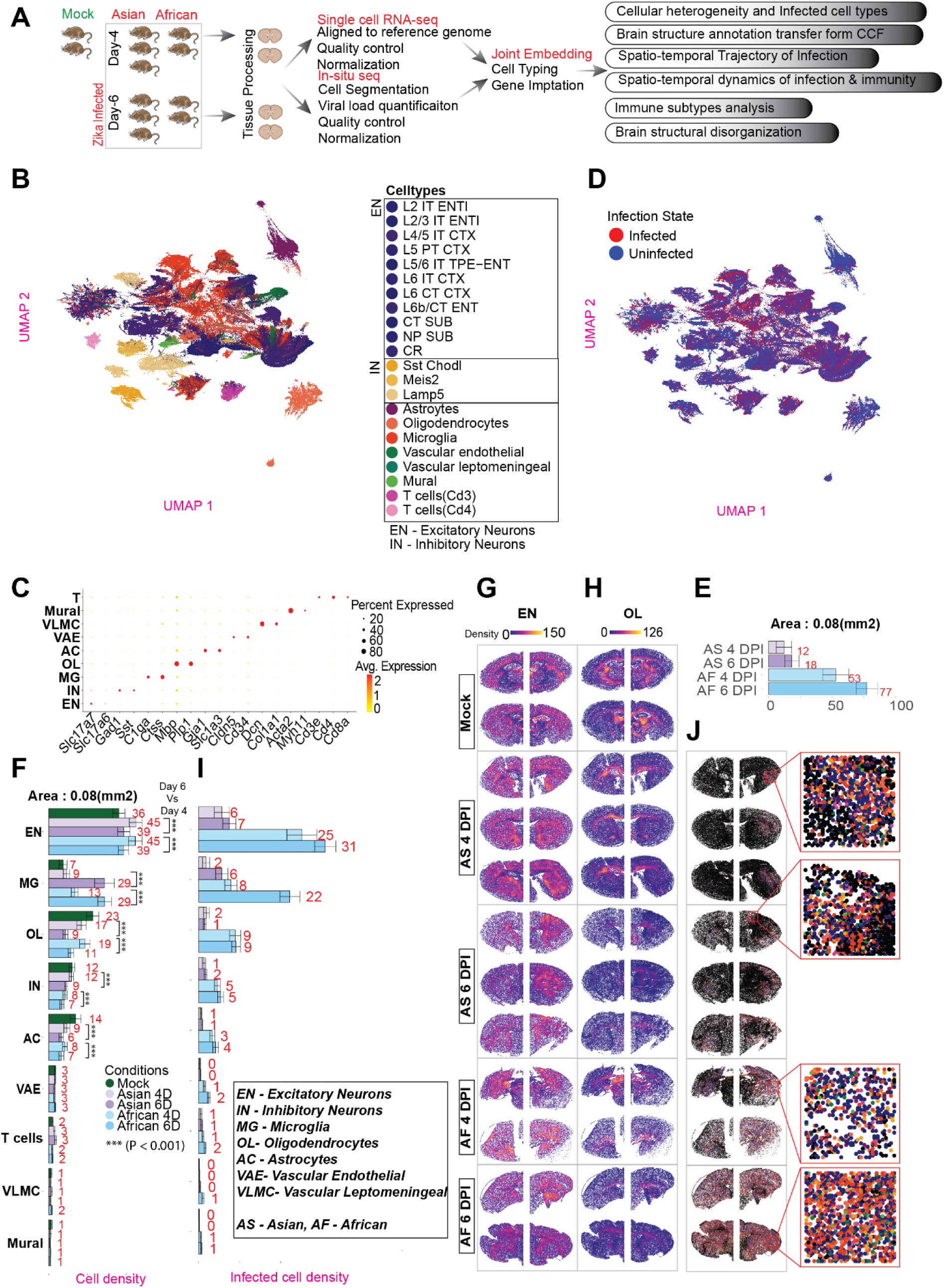
Integrated spatial and single-cell transcriptomic analysis captures cellular composition and infection dynamics in ZIKV-infected mouse brains. **(A)** Schematic overview of experimental design integrating spatially resolved single-cell transcriptomics with whole-transcriptome scRNA-seq to characterize cellular composition and ZIKV infection dynamics across viral strains (Asian and African) and time points (4 and 6 dpi). **(B)** UMAP embedding of spatially resolved cells colored by major cell type, revealing nine principal neuronal, glial, vascular, and immune cell populations. **(C)** Dot plot showing expression of representative marker genes used to annotate major cell types; dot size indicates fraction of cells expressing each gene, and color intensity denotes average expression level. **(D)** UMAP visualization distinguishing infected from uninfected cells within the spatial dataset. **(E)** Quantification of infected cell density across experimental conditions, demonstrating consistently higher infection density in African strain-infected brains at both time points. Local infected cell density was calculated by counting neighboring infected cells within a 0.08 mm² spatial neighborhood centered on each cell. Bars represent mean infected cell density across all individual cells; error bars indicate standard deviation across cells scaled by a factor of 6 (SD/6) to visualize relative dispersion within each condition**. (F)** Densities of major cell types across all conditions, including mock controls, revealing infection-associated expansion of microglia and concomitant reductions in neuronal and glial populations over time. Bars represent mean cell density; error bars indicate scaled standard deviation (SD/6). **(G-H)** Representative spatial maps showing anatomical distribution of excitatory neurons **(G)** and oligodendrocytes **(H)**, illustrating their characteristic localization in cortical/subcortical regions and white matter tracts, respectively. **(I)** Densities of infected cells stratified by cell type and condition, highlighting strain-specific differences in cellular tropism. Bars represent mean cell density across all individual cells; error bars indicate standard deviation across cells scaled by 6 (SD/6). **(J)** Spatial visualization of infected cells colored by infected cell type, revealing anatomical localization and cellular heterogeneity of ZIKV infection within the brain.

To characterize cellular composition in mock and infected brains, we employed a reference-guided, marker-based cell typing approach (see Methods). We first used a Bayesian classifier trained on a reference scRNA-seq atlas to generate high-confidence cell type predictions for the spatial transcriptomics data, leveraging shared gene expression patterns between datasets followed by identification of cell type marker genes through differential expression analysis. To enhance cell type resolution and enable transcriptome-wide analysis, we integrated the spatial transcriptomics data with scRNA-seq data from ZIKV-infected samples (including corresponding mock) using the LIGER framework [11]. This integration leverages the deeper sequencing depth of scRNA-seq data for more stable clustering and enables gene expression imputation across the full transcriptome for spatially resolved cells, which are limited to 218 genes. Joint embedding of both datasets showed no noticeable batch effects, with cells well-mixed across shared clusters regardless of sequencing technology, validating the integration (Fig. 1S3A-B). This analysis identified nine major cell types, including neurons, glia, vascular, and immune cells (Fig. 1B, C). Excitatory neurons (EN) were classified into 11 subtypes, inhibitory neurons (IN) into three, and T cells into two. Neuronal identity was defined by *Slc17a7*, *Slc17a6* (EN), and *Gad1*, *Gad2* (IN). Microglia (*C1qa, Ctss, Csf1r, Spp1*), oligodendrocytes (*Plp1, Mog, Mbp*), and astrocytes (*Slc1a3, Gja1*) were also well-defined, along with vascular (*Cldn5, Cd34*), leptomeningeal (*Dcn, Col1a1*), mural (*Acta2, Cavin1, Myh11*), and T cells (*Cd3e, Cd4, Cd8a*).

ZIKV-infected cells were identified across all infected samples, with infection rates ranging from 9% to 79% in spatial data (Fig. 1S2A) and 1% to 91% in scRNA-seq (Fig. 1S2B). Notably, the African strain exhibited consistently higher infection density (quantified as infected cell number within 0.08 mm² local neighborhoods; Fig. 1S3C, see Methods) than the Asian strain at all time points (Asian 4D = 12, Asian 6D = 18, African 4D = 53, African 6D = 77) (Fig. 1D, E). This widespread and severe infection in African strain samples aligns with prior reports of increased replication and neurovirulence associated with African ZIKV [9, 10] and motivated our investigation into the cellular and spatial mechanisms underlying these differences.

To assess CNS viral burden, we performed RT-qPCR on brain tissue at 4 and 6 dpi. African-lineage-infected brains contained significantly higher Zika RNA than Asian-lineage brains at both timepoints, with levels rising further by 6 dpi (Fig. 1S1C). Asian-lineage infection produced low, relatively stable viral RNA over the same interval. ZIKV protein NS1 immunofluorescence staining confirmed these spatial differences. Mock samples showed no detectable signal, while Asian-lineage infection resulted in sparse NS1-positive cells. In comparison, African-lineage infection produced widespread and intense NS1 staining throughout the parenchyma, particularly at 6 dpi (Fig. 1S1D). NS1 signal was cytoplasmic and clustered, indicative of focal viral expansion. Together, these findings establish that African-lineage ZIKV replicates more efficiently in the CNS, in line with the greater neuropathology observed *in vivo*.

In mock brains, neurons exhibited the highest density, with excitatory neurons dominating (density = 36), followed by inhibitory neurons (density = 12) (Fig. 1F, 1S4). Among glial cells, oligodendrocytes were most abundant (density = 23), followed by astrocytes (density = 14) and microglia (density = 7). Spatial mapping confirmed the expected anatomical organization, with excitatory neurons concentrated in cortical and subcortical regions and oligodendrocytes aligning with white matter tracts (Fig. 1G, H), consistent with known brain architecture.

Upon infection, significant alterations in cell densities emerged. Microglia expanded over time, peaking at 6 dpi, while excitatory neurons, inhibitory neurons, oligodendrocytes, and astrocytes declined (Fig. 1F). The increase in microglia suggests an active immune response, potentially resulting in phagocytosis of infected or damaged neurons [12, 13]. The loss of these neurons and some glial cells suggest a combination of direct viral cytopathology and secondary damage mediated by neuroinflammation. This cellular depletion may disrupt local neural circuits and impair normal brain development, mirroring neuropathological features seen in congenital Zika syndrome. Such findings are consistent with prior studies reporting microglia-driven neurotoxicity and glial dysregulation following ZIKV infection [14].

Strain-specific infection dynamics revealed higher infected cell densities in African strain-infected samples. Infected cell densities were consistently elevated in African strain-infected brains across multiple cell types, including neurons, glia, and vascular cells (Fig. 1I, J). This widespread infection suggests that the African strain’s increased neurovirulence may stem from enhanced viral entry, replication, or immune evasion strategies, as previously suggested by its greater NS1 protein stability and interferon antagonism [15, 16]. The broad infection of glial and vascular cells in African strain-infected brains further implies potential disruption of neurovascular integrity, which could exacerbate neuroinflammation and neuronal dysfunction. In fact, H&E staining revealed that brains infected by African strain manifested damages by day 4 and progressed to severe damages by day 6 while those infected Asian strain had mild or no damages at all the timepoints (Fig. 1S1E&F). Together, these findings highlight the distinct cellular and spatial impact of ZIKV infection in the brain, with African strains exhibiting more widespread and severe infection. The differential susceptibility of neural and immune cell populations provides insights into potential mechanisms driving ZIKV pathogenesis and underscores the importance of strain-specific differences in viral-host interactions.

### Spatiotemporal Trajectories of ZIKV Infection in the Mouse Brain

Having established that African and Asian ZIKV strains differ markedly in infection burden and cellular impact, we next sought to understand how infection propagates through brain tissue over time. Specifically, we asked whether infection spreads uniformly or follows preferential anatomical routes. To capture the continuum of infection progression, we ordered infected brain samples by the percentage of infected cells, generating a natural pseudo-temporal trajectory that represents increasing disease severity (Fig. 2A). The resulting ordering stratified samples from least to most infected: Asian strain 4 day post infection (AS4DPI), Asian strain 6 day post infection (AS6DPI), African strain 4 day post infection (AF4DPI), and African strain 6 day post infection (AF6DPI). This pseudo-temporal progression captures increasing infection burden over time and across strains, distinguishing the gradual expansion of infection in the Asian strain from the inherently more aggressive African strain. Importantly, this ordering is based on infection extent rather than actual experimental time points, providing a continuous view of disease progression.

**Figure 2.**
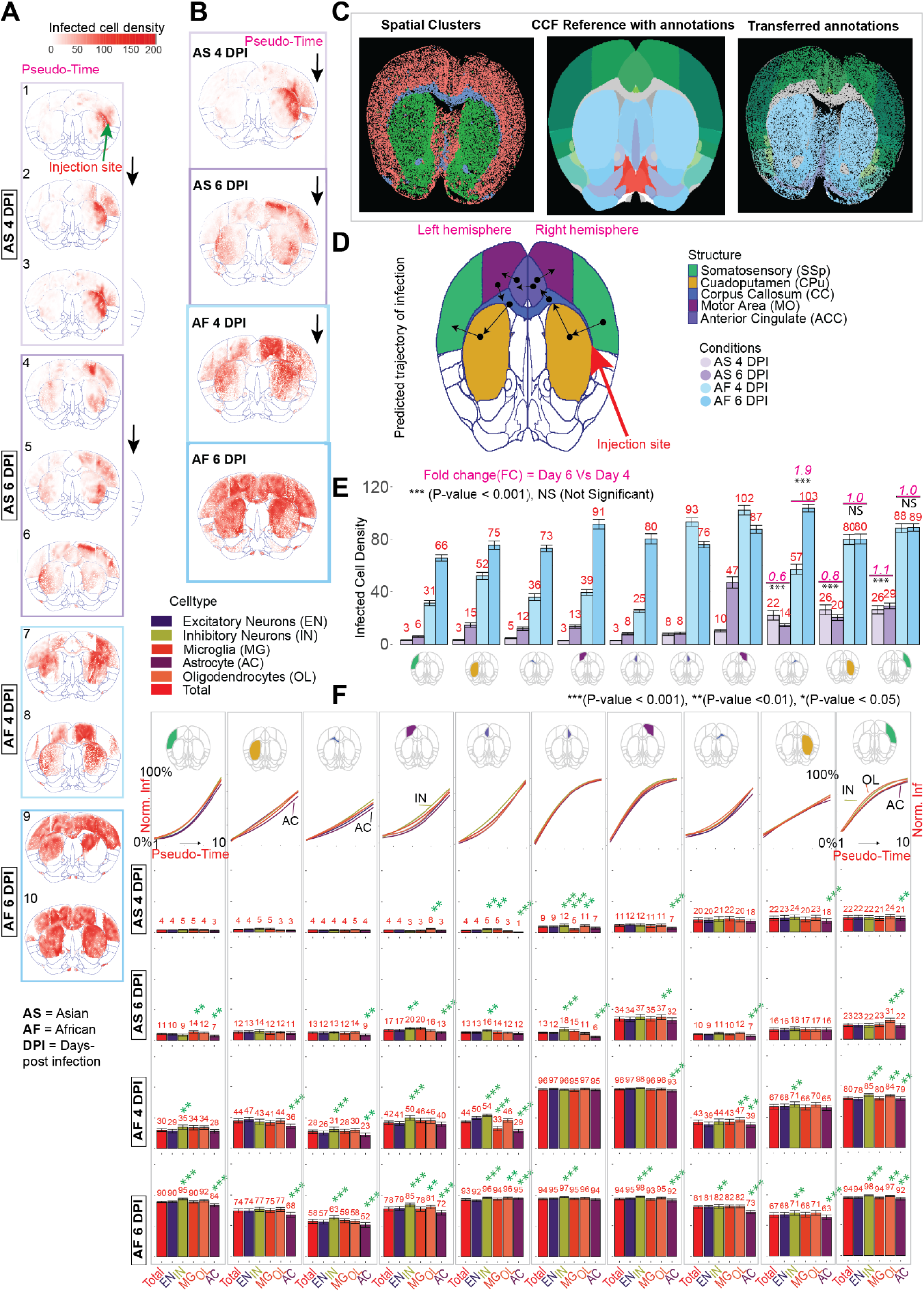
Spatiotemporal progression of ZIKV infection across brain structures and cell types. **(A)** Spatial tissue maps of all cells colored by local infected-cell density, calculated as the number of infected neighboring cells within a 0.08 mm² spatial neighborhood. Samples are ordered along a pseudo-temporal axis (1–10) defined by sample infection rate (percentage of infected cells per sample), capturing increasing infection burden across strains and time points. All cell coordinates are aligned to a common spatial reference frame (see Methods). **(B)** Condition-aggregated spatial tissue maps. For each condition (Asian 4 dpi, Asian 6 dpi, African 4 dpi, African 6 dpi), cells from all biological replicates were projected onto a shared reference space, revealing progressive spatial expansion of infection from the injection site in the right hemisphere to distal brain regions and ultimately the contralateral hemisphere. **(C)** Workflow for anatomical structure annotation and registration. Spatial transcriptomic data were aligned to the Allen Common Coordinate Framework (CCFv3) using spatially variable gene expression patterns, enabling transfer of anatomical structure labels to individual cells. **(D)** Inferred anatomical trajectory of ZIKV dissemination across brain structures. Arrows indicate the most likely direction of viral spread over pseudo-time based on spatial proximity and infection density dynamics. **(E)** Infection densities across major brain structures for each condition, showing time- and strain-dependent changes in regional infection burden. Bars represent mean infected-cell density across individual cells within each structure; error bars indicate standard deviation across cells scaled by 6 (SD/6) to visualize relative dispersion. **(F)** Cell-type-specific infection dynamics across brain structures and pseudo-time. Top: trends of normalized infection levels for major cell types across pseudo-time (1-10) within each brain structure. Bottom: bar plots showing normalized infection levels for excitatory neurons (EN), inhibitory neurons (IN), astrocytes (AC), microglia (MG), oligodendrocytes (OL), and total infection across structures and conditions. Bars represent mean normalized infection levels; error bars indicate standard deviation scaled by 6 (SD/6). Statistical significance markers indicate whether specific cell types are disproportionately infected relative to overall infection burden within each structure.

Spatial mapping of infected cells across this pseudo-temporal trajectory revealed non-random dissemination patterns (Fig. 2B; right side of each brain image corresponds to the mouse right hemisphere). At early pseudo-time points (least infected AS4DPI samples), ZIKV infection was primarily localized near the injection site in the right hemisphere. As infection progressed through the pseudo-temporal trajectory-spanning AS6DPI, AF4DPI, and ultimately AF6DPI samples, viral spread became increasingly widespread, extending to the left hemisphere and, in later stages, encompassing nearly the entire brain. Notably, the spatial trajectory of viral dissemination followed a distinct pattern rather than simple radial diffusion from the injection site, suggesting preferential routes of spread guided by neuroanatomical connectivity rather than uniform diffusion.

To elucidate how infection propagates across discrete brain structures, we employed reference-based anatomical segmentation. Accurately segmenting brain structures in infected samples is challenging due to infection-induced tissue disruption. To address this, we employed a reference-based approach, aligning spatial transcriptomic data to the CCFv3 reference atlas using QuickNII [17]. By leveraging spatially variable gene expression, we refined anatomical registration and transferred structural annotations to our dataset, enabling high-resolution mapping of infection spread (Fig. 2C, 2S1, 2S2, 2S3; see Methods). This approach enabled high-resolution tracking of viral dissemination from the injection site to distant structures, revealing potential differences in cellular susceptibility, immune activation, and tissue remodeling across brain regions.

Neurotropic viruses can disseminate through multiple pathways, including direct invasion of adjacent structures, axonal transport via peripheral nerves, and vascular dissemination [18, 19]. To systematically trace the spatial trajectory of infection, we developed a trajectory inference algorithm that integrates infection density with the spatial proximity of brain structures (see Methods). The algorithm models infection propagation based on both the spatial distance between brain regions and the temporal progression of infection burden. The inferred trajectory revealed preferential spread to the contralateral hemisphere through the anterior cingulate cortex (ACC), a region proximal to major white matter tracts including the corpus callosum (CC) (Fig. 2B, 2D). This pattern implicates neuroanatomical connectivity, particularly myelinated fiber tracts may facilitate axonal transport or provide permissive cellular environments, in guiding infection spread, consistent with established tissue mechanisms of neurotropic viral dissemination including direct invasion, axonal transport, and vascular routes.

Quantification of infection density within individual brain structures revealed a spatially heterogeneous pattern: while global infection burden increased over pseudo-time, early-infected regions near the injection site exhibited stable or declining infection densities from day 4 to day 6, suggesting active regional immune responses during this critical transition period (Fig. 2E). In the right somatosensory cortex (SSp-R), right caudate-putamen (CPu-R), and right corpus callosum (CC-R), structures heavily infected at 4 dpi, infection density plateaued or significantly decreased by 6 dpi in Asian strain samples (SSp-R: FC = 1.11, P < 0.05; CPu-R: FC = 0.77, P < 0.05; CC-R: FC = 0.64, P < 0.05). African strain samples showed similar stabilization in some regions (SSp-R: FC = 1.01, P > 0.05; CPu-R: FC = 1.00, P > 0.05), though CC-R infection continued to increase (FC = 1.91, P < 0.05). The stabilization or decline of infection at these early-infected sites, concurrent with rising infection in distant regions, strongly suggests that localized immune activation restricts viral replication near the injection site during the 4 to 6 dpi transition.

Our spatial mapping revealed that ZIKV infects all major brain cell types, including neuronal and glia cells across all major brain structures, a finding that extends beyond previous reports that have focused primarily on neuronal or progenitor cell infection[20]. To determine whether ZIKV exhibits preferential infection of specific cell types despite this broad cellular tropism, we calculated normalized infection densities accounting for both infection burden and baseline cell-type abundance (Fig. 2F, 2S5-2S6). At early pseudo-time points, ZIKV infected all major cell types, though astrocytes were already under-infected in several structures near the injection site. As infection burden increased across pseudo-time, inhibitory neurons showed disproportionately higher infection levels while astrocytes exhibited persistent under-infection across an expanding set of brain regions (Fig. 2F). Notably, anterior cingulate (ACC) regions were the earliest structures to show clear tropism patterns, with inhibitory neurons significantly over-infected relative to total infection beginning at AS4DPI. The preferential infection of inhibitory neurons suggests these cells may be particularly susceptible to sustained ZIKV replication, or that other cell types engage more effective antiviral responses. Given the essential role of inhibitory neurons in maintaining excitatory-inhibitory balance and regulating synaptic transmission, their selective infection may directly contribute to the motor dysfunction and behavioral deficits observed in ZIKV-infected animals[21]. Collectively, our findings delineate a structured progression of ZIKV dissemination in the mouse brain, influenced by anatomical connectivity and cell-type susceptibility. Notably, the African strain demonstrates heightened neuroinvasiveness. The preferential infection of inhibitory neurons may underlie observed neurological dysfunctions.

### Spatial Dynamics of Microglia Response to Zika Infection

Having observed that infection stabilizes in injection-proximal regions (Somatosensory: SSp-R, Cuadoputamen: CPu-R and Corpus Callosum: CC-R) while spreading to distant structures, we hypothesized that differential microglial recruitment underlies this spatial heterogeneity in infection control. To test this, we examined microglial spatial distribution and recruitment dynamics in different brain regions.

At early time points (4 dpi) microglia were accumulated near the injection site, with substantially higher densities in injection-proximal structures (SSp-R: 14; CPu-R: 16; CC-R: 14) compared to distant contralateral structures (SSp-L: 5; CPu-L: 6; CC-L: 6), suggesting an immediate and spatially localized immune response (Fig. 3A, B). African 4D samples showed a similar spatial pattern in injection-proximal (SSp-R: 22; CPu-R: 19; CC-R: 20) and distant regions (SSp-L: 9; CPu-L: 10; CC-L: 15). Microglial density increased over time (6 dpi) in injection-proximal structures, with Asian strain showing substantially greater fold changes than African strain (SSp-R: Asian FC 6 dpi vs 4 dpi = 3.14 vs African FC = 1.36; CPu-R: Asian FC = 3.25 vs African FC = 1.63; CC-R: Asian FC = 2.25 vs African FC = 1.70; MO-R: Asian FC = 5.50 vs African FC = 1.58) (Fig. 3A, B, 1F), highlighting faster microglial recruitment in Asian strain samples. However, given the significantly higher infection burden in African strain samples, understanding how microglia contribute to infection containment is critical.

**Figure 3.**
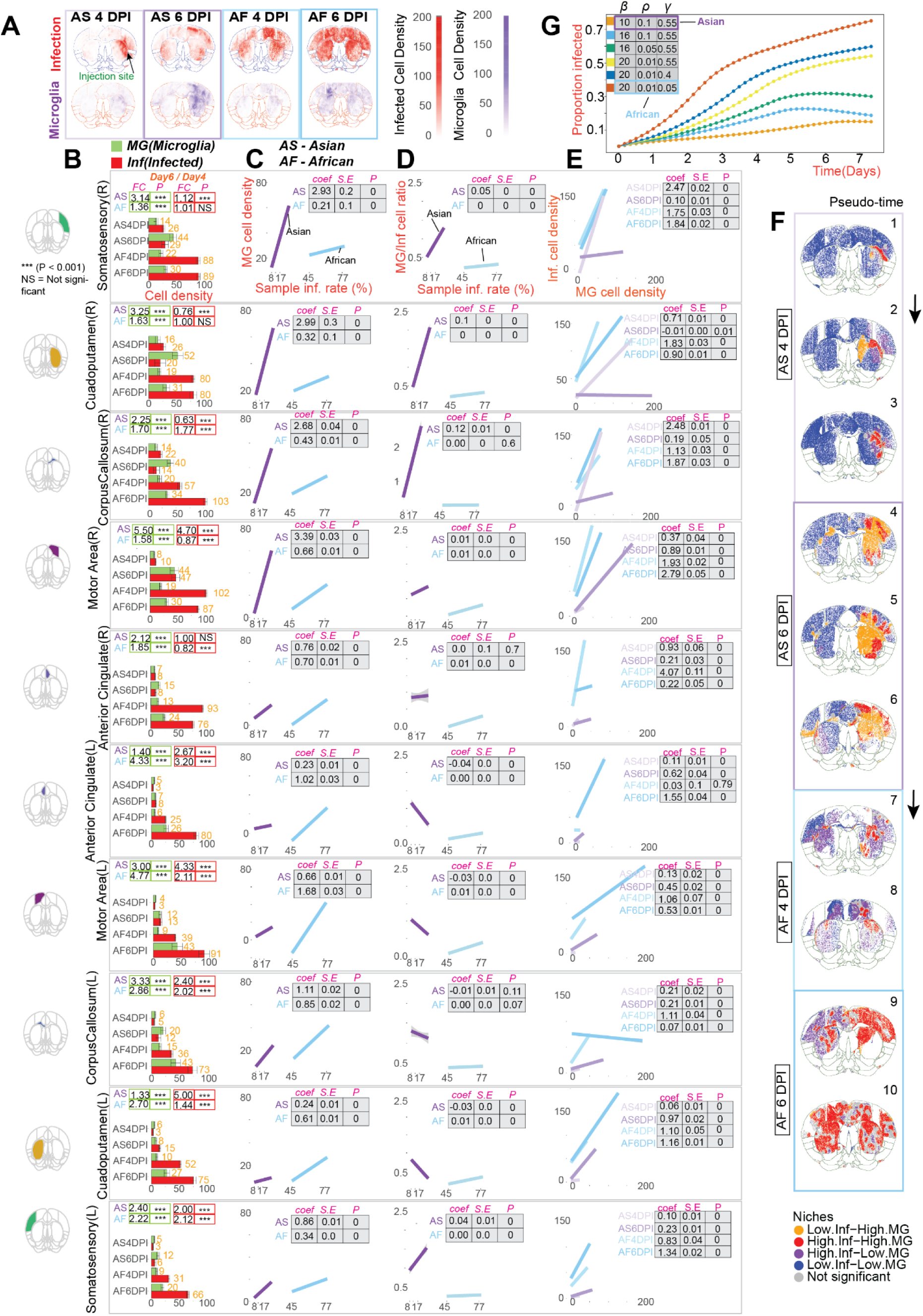
Spatial dynamics and effectiveness of microglial responses to ZIKV infection. **(A)**Spatial tissue maps showing local densities of infected cells (top) and microglia (bottom) across major brain structures, including somatosensory cortex (SSp), caudate-putamen (CPu), motor area (MO), corpus callosum (CC), and anterior cingulate area (ACA), under four infection conditions: Asian 4 dpi (AS4DPI), Asian 6 dpi (AS6DPI), African 4 dpi (AF4DPI), and African 6 dpi (AF6DPI). Local infection and microglial densities were calculated by counting neighboring infected cells and microglia, respectively, within a 0.08 mm² tissue microenvironment centered on each cell. **(B)** Bar plots summarizing mean local densities of infected cells and microglia within each brain structure across infection conditions. Fold changes (FC) comparing 6 dpi to 4 dpi are shown for each strain. Bars represent mean local density across individual cells; error bars indicate standard deviation across cells scaled by 6 (SD/6) to visualize relative dispersion. Green outlined boxes denote microglial density comparisons; red outlined boxes denote infected cell density comparisons. **(C)** Microglial recruitment kinetics along the infection trajectory. Pseudo-time, defined by sample infection rate (percentage of infected cells per sample), was used as a proxy for infection progression. Local microglial density was regressed against sample infection rate using linear models fitted separately for Asian and African strain-infected samples within each brain structure. Regression coefficients (coef), standard errors, and P-values are shown; steeper slopes indicate faster microglial accumulation as infection burden increases. **(D)** Immune efficiency as infection burden increases, quantified by the microglia-to-infected cell ratio. The ratio of local microglial density to infected cell density was regressed against sample infection rate. Higher ratios indicate greater immune presence relative to infection burden. Differences in regression slopes reveal strain-specific changes in immune efficiency over the course of infection. **(E)** Relationship between local microglial density and infection burden within tissue microenvironments. Infected cell density was modeled as a function of local microglial density within 0.08 mm² neighborhoods using linear regression, with separate models fitted for each condition. Regression coefficients are shown for injection-proximal structures (SSp-R, CPu-R, CC-R). Flatter or negative slopes indicate reduced sensitivity of infection to increasing microglial presence, consistent with effective containment, whereas steeper positive slopes indicate continued infection expansion despite microglial accumulation. **(F)** Spatial co-distribution of infection and microglial responses assessed by bivariate Moran’s I analysis. Cells were classified into four significant spatial niches (P < 0.01): High infection-High microglia (High.Inf-High.MG), Low infection-High microglia (Low.Inf-High.MG), High infection-Low microglia (High.Inf-Low.MG), and Low infection-Low microglia (Low.Inf-Low.MG). Percentages indicate the proportion of each niche across conditions. Gray regions denote locations without significant spatial association. **(G)** Simulation of infection dynamics under varying immune and virulence parameter regimes. The proportion of infected cells over time is shown for scenarios generated by systematically varying infection expansion potential (β), microglial effectiveness in dampening spread (γ), and infected cell clearance probability (ρ). Parameter regimes with higher microglial effectiveness (γ ≥ 0.55) exhibit stabilization of infection over time, whereas regimes with reduced immune effectiveness (γ ≤ 0.4), particularly when combined with high virulence (β = 20), show sustained infection expansion without stabilization.

Critically, by AS6DPI, injection-proximal regions exhibited stable or significantly reduced infection densities (SSp-R: FC 6 dpi vs 4 dpi = 1.12, P<0.001; CPu-R: FC = 0.76, P<0.001; CC-R: FC = 0.63, P<0.001), indicating potential infection containment, possibly mediated by early microglial activation (Fig. 3B). In contrast, AF4DPI samples already displayed higher infection densities (SSp-R: 88; CPu-R: 80; CC-R: 57) but without correspondingly robust microglial responses relative to infection burden (Fig. 3B).

To quantify the kinetics of microglial recruitment, we performed linear modeling using microglial cell density as the response variable and pseudotime (infection percentage in sample) as the predictor, yielding regression coefficients that reflect microglial recruitment rate. The Asian strain exhibited significantly steeper recruitment slopes than the African strain in regions proximal to the injection site (SSp-R: Asian 2.93 vs African 0.20, P<0.001; CPu-R: Asian 2.99 vs African 0.32, P<0.001; CC-R: Asian 2.68 vs African 0.43, P<0.001) (Fig. 3C), indicating faster and more coordinated microglial mobilization. None of the remaining structures showed such dramatic increases in microglia. These findings suggest that early and efficient microglial recruitment in the Asian strain may contribute to more effective containment of infection.

The effectiveness of the microglial response can be assessed through the microglia-to-infected cell ratio in local microenvironments (0.08 mm²), a metric of immune efficiency. This ratio was consistently higher in structures near the injection site in Asian strain-infected samples compared to African strain samples (SSp-R: Asian 0.05 vs African 0.00, P<0.001; CPu-R: Asian 0.1 vs African 0.00, P<0.001; CC-R: Asian 0.12 vs African 0.00, P<0.001) (Fig. 3D). A higher microglia-to-infected cell ratio in the Asian strain suggests enhanced immune surveillance, faster viral clearance, and reduced bystander damage to surrounding neural tissue. In contrast, African strain-infected samples may suffer from an insufficient microglial response, allowing greater viral spread and increased disease severity.

To directly evaluate how infection responds to increasing microglial presence, we modeled infected cell density as a function of microglial cell density within local tissue microenvironments (0.08 mm²) using linear regression (Fig. 3E). This approach allowed us to estimate the degree to which infection expands or contracts as microglia accumulate. In AS4DPI samples (pseudo-time 1-3), infected cell density increased with microglial density in structures near the injection site (SSp-R: 2.47, P<0.001; CPu-R: 0.71, P<0.001; CC-R: 2.48, P<0.001), suggesting that microglial recruitment was still occurring in response to ongoing viral spread. By AS6DPI (pseudo-time 4-6), however, this relationship flattened (SSp-R: 0.10, P<0.001; CPu-R: -0.01, P<0.001; CC-R: 0.19, P<0.001), indicating that infection progression had stabilized despite sustained microglial presence, evidence of successful containment. In contrast, both AF4DPI (SSp-R: 1.75, P<0.001; CPu-R: 1.83, P<0.001; CC-R: 1.13, P<0.001) and AF6DPI (SSp-R: 1.84, P<0.001; CPu-R: 0.90, P<0.001; CC-R: 1.87, P<0.001) samples (pseudo-time 7-10) maintained strong positive associations between microglial and infected cell densities throughout, indicating that infection persisted or expanded despite microglial accumulation.These findings underscore a key distinction: while microglial recruitment is essential, it is the timing and effectiveness of that recruitment that determines outcome. In the Asian strain, microglia appear to respond early and then suppress infection, whereas in the African strain, microglia may be recruited too late or fail to control the infection effectively.

The spatial relationship between microglia and infected cells provides mechanistic insight into differential infection outcomes in AS6DPI samples. We applied bivariate Moran’s I analysis to classify microglia-infection co-distribution into four spatial niches (P <0.01): High infection-High microglia, High infection-Low microglia, Low infection-High microglia, and Low infection-Low microglia (Fig. 3F; see Methods). Notably, the Low infection-High microglia niche was more prevalent in AS6DPI samples (35.1%) (Fig. 3F, 3S1A) suggesting that robust microglial activation contributed to viral containment. Conversely, the AF4DPI samples had the highest proportion of High infection-Low microglia regions (39.7%), reflecting an insufficient early immune response. By AF6DPI, microglia recruitment intensified, shifting to a higher proportion of High infection-High microglia regions (67.5%), indicating a delayed but strong immune activation. We quantified the proportions of these spatial niches in different brain structures and revealed that injection-proximal regions had the highest prevalence (SSp-R = 56.8%, CPu-R = 65.5%, CC-R = 76.7%) of Low infection-High microglia regions (Fig. 3S1A-B), reinforcing the importance of early microglial recruitment for effective viral containment.

Beyond their relationship to infected cells, microglia exhibited preferential spatial associations with specific cell types. By analyzing microglia density relative to different types of neighboring cells (see Methods). We revealed that microglia predominantly clustered around other microglia, a pattern observed in both viral strains (Fig. 3S1C). This suggests that microglia engage in cooperative active immune responses, potentially amplifying neuroprotection, phagocytosis and immune signaling. Beyond self-clustering, microglia were frequently positioned near excitatory neurons, suggesting a targeted immune response essential for preserving neural homeostasis during infection through interactions that modulate synaptic activity and maintain neural circuit integrity [22, 23].

To gain a deeper understanding of the broader dynamic interplay between infection spread and microglia recruitement beyond the experimentally measured time points, we modeled infection dynamics using stocastic, a density based framework inspired by Reed-Frost chain-binomial model [24] (see Methods), where local infection dynamics are governed by three key parameters: β (virulence) represents infection expansion potential, γ (immune effectiveness) quantifies effectiveness of microglia in dampening infection spread and ρ denotes the probability that an infected cell is cleared (see Methods). We fitted seprate parameter regimes for Asian and African strains to the observed data, enabling us to simulate the full temporal interplay between infection expansion, microglial recruitment, and clearance beyond the experimentally measured time points (Video S1). For the Asian strain, the inferred parameter regime (β = 10, γ = 0.55, and ρ = 0.1; Fig. 3S2) indicates controlled infection expansion coupled with effective microglial containment, consistent with the stabilization seen in the spatial data. In contrast, the African strain’s parameters (β = 20, γ = 0.05, and ρ = 0.01, Fig. 3S2) reflect rapid viral amplification and weak microglial restraint, explaining the continued infection growth and limited immune control. To further dissect the condition under which microglia fail to effectively contain infection, we performed systematic parameter-tuning that interpolates between the Asian and African strain regimes. Rather than treating these strains as discrete categories, this analysis explores a continuum of infection dynamics by gradually varying the virulence (β), immune dampening (γ), and recovery (ρ) parameters while monitoring the resulting proportion of infected cells over time (Fig. 3G; see Methods). Despite differences in intrinsic virulence (β = 10-16) and recovery rate (ρ = 0.05-0.1), these scenarios share a relatively strong microglia effectiveness (γ = 0.55). Under these conditions, the proportion of infected cells initially increases but progressively flattens with increasing time, indicating that microglial-mediated immune pressure is sufficient to counterbalance ongoing infection expansion. This stabilization behavior suggests that once microglial effectiveness exceeds a critical threshold, increases in virulence can be tolerated without leading to uncontrolled infection. In contrast, scenarios with high virulence (β = 20) combined with reduced microglia effectiveness (γ ≤ 0.4) and low recovery (ρ = 0.01), the proportion of infected cells continues to increase monotonically over time without evidence of stabilization. This sustained growth indicates a failure of microglia to effectively suppress infection spread. These findings underscore the role of timely and robust microglial activation in limiting ZIKV spread and suggest that modulating microglial responses may be a potential therapeutic strategy to improve infection outcomes.

### Microglia Subtype Contributions to Infection Containment

Having established that microglial recruitment timing and effectiveness determine infection outcomes, we asked whether specific microglial subtypes contribute disproportionately to viral containment. Microglia, display diverse functional states, enabling them to respond adaptively to physiological and pathological cues and influence neurodevelopment, synaptic function, and disease progression [25, 26]. Based on established subtype markers [27], we identified five distinct microglial states (Fig. 4A; see Methods). They include Inflammatory (*Cd72, Ccl12, Ctss, C1qa*), representing classical pro-inflammatory microglia involved in immune activation and pathogen response; Inflammatory_1 (*Gzma, Gzmb, Nkg7*), characterized by granzyme-expressing microglia, representing an enhanced cytotoxic immune function; Disease-associated or DAM (*Spp1, Ifitm1, Hba.a1, Hbb.bs*), previously linked to neurodegenerative conditions, indicating a role in infection-related tissue remodeling and immune modulation; Stressed (*Mt1, Vim, Dusp1*), marked by stress-response genes reflecting activation under adverse conditions; and Homeostatic (*P2ry12, Hexb, Trem2*), maintaining normal surveillance and neuroprotective functions (Fig. 4B).

**Figure 4.**
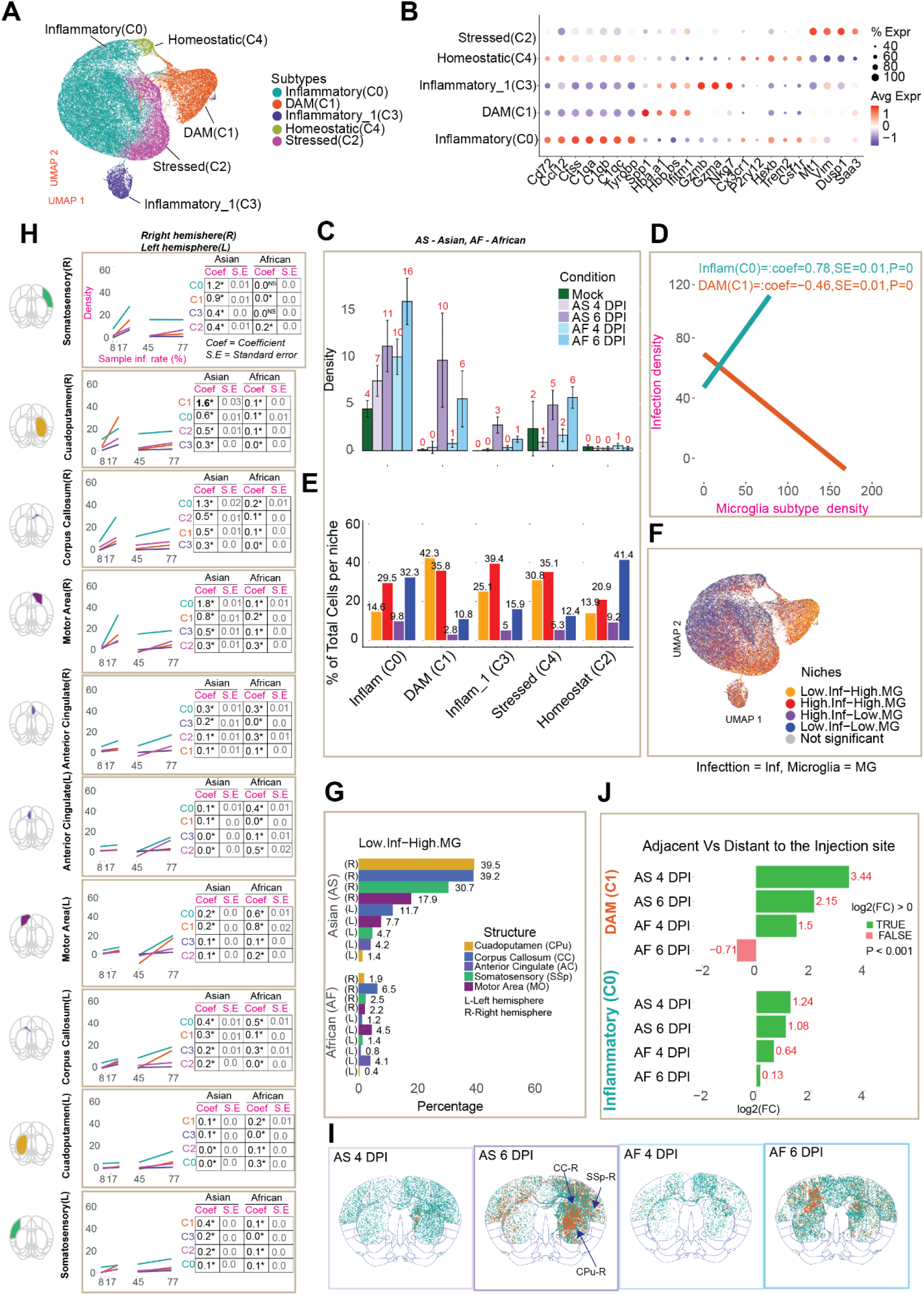
Microglial heterogeneity and subtype-specific contributions to ZIKV infection containment. **(A)**UMAP visualization of microglial cells colored by inferred microglial state, revealing five distinct subtypes: Inflammatory, Inflammatory_1, Disease-associated microglia (DAM), Stressed, and Homeostatic. **(B)** Dot plot showing representative marker gene expression profiles for each microglial subtype. Dot color indicates average expression level; dot size denotes the fraction of cells expressing each gene. **(C)** Bar plot showing local microglial subtype densities across infection conditions. Local subtype densities were computed by counting neighboring microglia of the same subtype within 0.08 mm² spatial neighborhoods centered on each cell. Bars represent mean subtype density across individual cells; error bars indicate standard deviation across cells scaled by 6 (SD/6). **(D)** Fitted relationships between local microglial subtype density and local infected-cell density within tissue microenvironments. Local densities of DAM and Inflammatory microglia were regressed against infected-cell density within 0.08 mm² neighborhoods. Regression coefficients and standard errors quantify the strength and direction of association, revealing opposing subtype-specific relationships with infection burden (DAM: negative association; Inflammatory: positive association). **(E)** Bar plot showing the distribution of infection-microglia spatial niches within each microglial subtype, expressed as the proportion of cells in each niche category. Bars represent mean proportions across cells of each subtype. **(F)** UMAP visualization of microglia colored by infection-microglia spatial niche classification (High infection-High microglia, Low infection-High microglia, High infection-Low microglia, Low infection-Low microglia). **(G)** Bar plot showing the proportion of infection-contained regions (Low infection-High microglia niche) within each major brain structure across conditions. **(H)** Microglial subtype recruitment kinetics across major brain structures. Local subtype densities were computed within 0.08 mm² neighborhoods and regressed against sample infection rate (percentage of infected cells per sample) to estimate structure- and subtype-specific accumulation dynamics. Regression coefficients are shown for both DAM and Inflammatory subtypes in Asian and African strain samples. Strain labels (Asian/African) are indicated in panel headers. **(I)** Spatial tissue maps showing microglial cells projected onto a common reference brain and colored by DAM and Inflammatory microglia subtype identity, illustrating subtype-specific anatomical organization across infected brains. **(J)** Bar plot showing enrichment of microglial subtypes in injection-proximal regions (SSp-R, CPu-R, CC-R) relative to remaining brain structures. Bars represent log₂ fold change comparing subtype density in injection-proximal regions versus all other structures. Positive values indicate enrichment near the injection site.

At early days (Asian & African 4 dpi), microglia were predominantly composed of a single state, Inflammatory (Density: AS4DPI = 7; AF4DPI = 10 and Mock = 4), suggesting a rapid immune activation in response to viral presence (Fig. 4C). This early dominance likely reflects an immediate pro-inflammatory reaction aimed at containing infection. However, by 6 dpi, microglial state composition became more diverse in addition to increased density (AS6DPI: Inflammatory = 11, DAM = 10, Stressed = 5, Inflammatory_1 = 3, Homeostatic = 0; AF6DPI: Inflammatory = 16, DAM = 6, Stressed = 6, Inflammatory_1 = 1, Homeostatic = 0), with notable increases in DAM and other states. Given that Inflammatory and DAM were the most prevalent subtypes, we examined whether their spatial distribution differed between strains and related to infection outcomes.

To test whether specific subtypes contribute directly to infection resolution, we examined the relationship between subtype abundance and infected cell density within local microenvironments (0.08 mm²). Local environments with higher DAM density exhibited significantly decreased infection burden (coefficient: -0.46, S.E.: 0.01, P < 0.001; Fig. 4D), indicating a critical role in restricting viral spread. In contrast, Inflammatory microglia density increased proportionally with infection density (coefficient: 0.78, S.E.: 0.01, P < 0.001; Fig. 4D), suggesting that while these cells are recruited in response to infection, their presence alone does not ensure containment. Inflammatory microglia are typically associated with prolonged neuroinflammation, cytokine production, and tissue damage rather than effective viral restriction. This aligns with prior studies showing that DAM-like microglia contribute to immune resolution and tissue repair, whereas persistent inflammatory activation may exacerbate neurotoxicity [28].

Spatial niche analysis using bivariate Moran’s I further distinguished the roles of these subtypes. Within the DAM population, the highest proportion of cells (42.3%) were in Low infection-High microglia niches regions where infection is successfully contained (Fig. 4E, F). In contrast, Inflammatory microglia were distributed across Low infection-Low microglia (32.3%) and High infection-High microglia (29.5%) niches, consistent with their recruitment to sites of ongoing infection without necessarily achieving containment. These findings indicate that DAM accumulation is strongly associated with successful infection control, while Inflammatory microglia proliferate in response to infection but does not drive resolution.

To understand the spatial organization underlying these differential outcomes, we quantified the prevalence of infection-contained regions (Low infection-High microglia) across brain structures (Fig. 4G). In Asian strain samples, the highest proportions of contained regions were found in injection-proximal structures: Cuadoputamen (CPu-R) (39.5%), Corpus Callosum (CC-R) (39.2%), Somatosensory (SSp-R) (30.7%), and MO-R (17.9%). Notably, none of the brain structures in African strain samples reached double-digit percentages of contained regions, underscoring the failure of effective immune responses in African strain infection.

We next examined whether the kinetics of subtype-specific recruitment differed between strains. Using linear modeling with subtype density as the response variable and pseudo-time as the predictor, we found that both DAM and Inflammatory subtypes increased most rapidly with infection in Asian strain injection-proximal regions (Fig. 4H, I) (DAM: CPu-R coefficient = 1.6, P < 0.001; SSp-R coefficient = 0.9, P < 0.001; MO-R coefficient = 0.8, P < 0.001; CC-R coefficient = 0.5, P < 0.001). In contrast, none of the corresponding regions in African strain samples showed comparable subtype recruitment rates. Instead, in African samples, modest increases in DAM and Inflammatory subtypes occurred in distant regions (DAM: MO-L coefficient = 0.8, P < 0.001; CC-L coefficient = 0.3, P < 0.001; Inflammatory: MO-L coefficient = 0.6, P < 0.001; CC-L coefficient = 0.5, P < 0.001), where they failed to achieve infection containment. Furthermore, DAM and Inflammatory subtypes were significantly enriched near the injection site in Asian samples compared to African samples (AS4DPI: log2FC = 3.44, P < 0.001; AS6DPI: log2FC = 2.15, P < 0.001) in SSp-R, CPu-R, and CC-R (Fig. 4J). The early and robust recruitment of DAM to injection-proximal regions in Asian strain samples precisely where infection stabilizes, reinforces the conclusion that DAM plays a key role in sustaining immune resolution and preventing viral dissemination.

Collectively, these findings demonstrate that not all microglia are equally effective at infection control. The accumulation of DAM in early infected brain regions where infection is better controlled suggests a key role in sustaining immune resolution and preventing viral dissemination. DAM microglia have been implicated in neurodegenerative diseases, where they aid in clearing debris and restricting tissue damage [29]. Their accumulation near infection sites suggests they may contribute to viral clearance by enhancing phagocytosis, promoting tissue repair, or limiting viral replication through immune signaling.

### Microglia functional dynamics and cell-cell interactions in response to infection

Having established that DAM microglia drive infection containment while Inflammatory microglia does not, we sought to understand the molecular basis underlying these functional differences. We first examined transcriptional responses of microglia to ZIKV infection to identify core activation programs and strain-dependent differences.

Differential expression analysis revealed robust activation of conserved immune programs in infected microglia across both viral strains and time points (Fig. 5A). Both Asian and African strain-infected microglia upregulated canonical antiviral and interferon-stimulated genes (Oasl1, Oas2, Oas3, Oas1a, Gbp6, Slfn4), inflammatory and chemotactic regulators (Cxcl9, Ccl8), acute phase response genes (Lcn2, Saa2), and innate immune effectors (Plac8, Chil3), consistent with a shared type I interferon-driven response. Functional enrichment analysis confirmed that infected microglia from both strains consistently exhibited enrichment of innate immune response, inflammatory response, TNF production, cell killing, and phagocytosis pathways at both 4 dpi and 6 dpi (Fig. 5B). These results define a core microglial activation program induced by ZIKV infection, characterized by interferon-mediated antiviral defense and inflammatory effector functions shared across strains.

**Figure 5.**
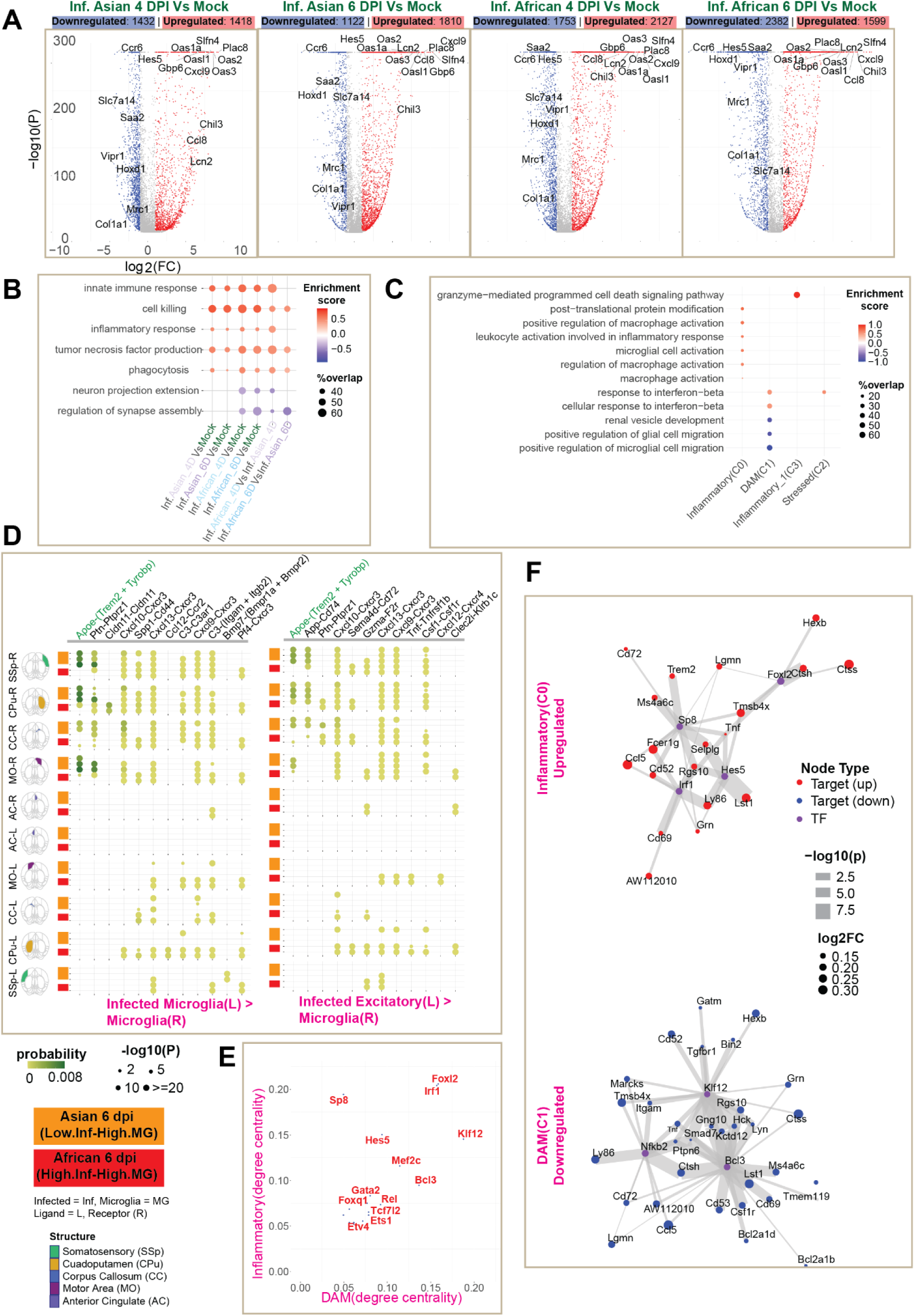
Infection-induced cellular disruptions and functional impairment of brain structural cell types. **(A)** Volcano plots showing differential gene expression in infected microglia compared to mock controls. Each panel compares infected microglia versus mock for: Asian 4 dpi, Asian 6 dpi, African 4 dpi, and African 6 dpi. X-axis: log₂ fold change; y-axis: –log₁₀ adjusted P-value. Genes with |log₂ fold change| ≥ 1.0 and adjusted P-value < 0.001 are considered significant (upregulated: red; downregulated: blue; non-significant: gray). Total numbers of significantly up- and downregulated genes are indicated for each comparison. **(B)** Bubble plots summarizing significantly enriched Gene Ontology biological process (GO:BP) terms identified by gene set enrichment analysis (GSEA; adjusted P-value < 0.05). Comparisons include infected microglia versus mock controls for Asian and African strains at 4 dpi and 6 dpi, as well as direct strain-to-strain comparisons at matched time points. Bubble color indicates enrichment score (red: positive enrichment; blue: negative enrichment); bubble size reflects the proportion of genes within each gene set that are significantly differentially expressed (|log₂ fold change| ≥ 0.30 and adjusted P-value < 0.05). **(C)** Functional pathway enrichment for microglial subtypes. Gene Ontology biological process terms enriched in each microglial subtype compared to the Homeostatic subtype baseline. Bubble color indicates enrichment score; bubble size reflects proportion of significant genes. **(D)** Cell-cell communication networks between infected cells and microglia in spatially distinct niches. CellChat-inferred ligand-receptor interactions between infected sender cell types (microglia, excitatory neurons, inhibitory neurons, oligodendrocytes) and microglia as receivers. Analysis restricted to infection-contained regions (Low infection-High microglia) in Asian 6 dpi samples (top rows, yellow) versus highly infected regions (High infection-High microglia) in African 6 dpi samples (bottom rows, red). Bubble color represents communication probability; bubble size denotes interaction significance (–log₁₀ P-value). **(E)** Scatter plot showing transcription factors (TFs) identified by CellOracle as significant regulators (P < 0.05) of DAM and Inflammatory microglial subtypes. Each point represents a TF, positioned by its degree centrality within the inferred gene regulatory networks of DAM (x-axis) and Inflammatory microglia (y-axis). **(F)** Gene regulatory networks linking top transcription factors (TFs; purple nodes) to their significant target genes (adjusted P-value < 0.05) that are also differentially expressed compared to Homeostatic microglia baseline. Target gene nodes are colored red (upregulated) or blue (downregulated) relative to Homeostatic state; node size represents absolute log₂ fold change. Edge width reflects strength of TF-gene regulatory prediction from CellOracle (–log₁₀ P-value). Top panel: Inflammatory microglia showing TF-driven upregulation of pro-inflammatory targets. Bottom panel: DAM microglia showing TF-driven downregulation of inflammatory targets.

Despite this shared baseline, critical strain-dependent differences emerged in the magnitude and functional balance of microglial responses. African strain infection triggered broader transcriptional perturbation than Asian strain at both time points (African 4 dpi: 2127 up/1753 down; African 6 dpi: 1599 up/2382 down; versus Asian 4 dpi: 1418 up/1432 down; Asian 6 dpi: 1810 up/1122 down) (Fig. 5A). More importantly, the functional character of these responses differed markedly between strains. At 4 dpi, African strain-infected microglia exhibited significantly stronger enrichment of innate immune, inflammatory, and effector-associated programs (TNF production, phagocytosis, cell killing) compared with Asian strain-infected microglia, consistent with early and amplified activation. By 6 dpi, African strain-infected microglia remained strongly enriched for effector functions while also showing prominent suppression of neuron-support pathways (neuron projection extension, regulation of synapse assembly) (Fig. 5B). These strain-dependent transcriptional patterns align with our microglia state-resolved results from Section 4. The Inflammatory microglial state is enriched for macrophage/microglia activation and activation involved in inflammatory response (Fig. 5C) and colocalizes with infection and increases with infection burden in African samples. The DAM state shows selective enrichment of interferon-β response terms, while showing downregulation of glial cell migration. DAM expands from 4 to 6 dpi in Asian infection-clearing regions (Low infection – High microglia) and inversely correlates with infection density. The downregulation of migration pathways in DAM suggests these cells adopt a stationary, resident phenotype focused on local debris clearance and interferon-mediated antiviral defense, contrasting with the more mobile, infiltrating character of Inflammatory microglia. The concurrent enrichment of inflammatory and cytotoxic programs alongside reduced expression of neuron-associated functions in African strain infection reflects a functional trade-off, with microglial functions shifting toward immune effector programs at the expense of neuro-supportive roles. Regional analysis of functional enrichment patterns revealed that these strain-dependent differences in innate immune and inflammatory pathway activation were consistent across brain structures (Fig. 5S1-8). Critically, by 6 dpi, African strain-infected microglia showed prominent downregulation of neuron projection extension and synapse assembly pathways in injection-proximal regions (SSp-R, CPu-R, CC-R, MO-R)—the same regions that showed the highest infection containment (Low infection-High microglia niches) in Asian strain samples (Fig. 4), whereas Asian strain-infected microglia in these regions maintained neuron-support functions. Specifically, key neuron-supportive genes including Ntrk2, Nrxn1, Caskin1, Ppg1r9a, Ptprd, Mef2c, Cxcl12, Csf1r, Bcan, and Slc39a12 were more strongly downregulated in African 6 dpi compared to Asian 6 dpi (relative to mock), while effector-associated genes including Thbs1, Msr1, Il2rb, Nos2, Cxcl9, Klrk1, and Lgals3 were more strongly upregulated in African samples (Fig. 5S1-8). This loss of homeostatic microglial functions in African strain infection at sites where containment is most successful in Asian strain may explain the failure of effective infection control by impairing the balance between antiviral defense and tissue preservation. These divergent functional outcomes in comparable brain regions, successful containment with preserved homeostatic functions versus failed containment with suppressed neuron-support programs, motivated us to investigate what upstream signals from infected cells might drive these distinct microglial states in infection-contained (Low infection - High microglia in Asian 6 dpi) versus uncontained (High-infection- High microglia) spatial niches. Importantly, similar functional changes occur in uninfected microglia within infected brains (Fig. 5S1-8), suggesting that viral infection induces tissue-wide immune state changes even in bystander cells.

To identify signals that might promote distinct microglial states, we analyzed cell-cell communication networks between infected cells (as ligand sources) and microglia (as receptor targets) in spatially distinct niches. We restricted our analysis to infection-contained regions (Low infection-High microglia) in Asian 6 dpi samples versus highly infected regions (High infection-High microglia) in African 6 dpi samples, allowing us to identify signaling patterns associated with successful versus failed infection control. Our analysis revealed distinct signaling patterns between these spatial niches (Fig. 5D). In infection-contained regions of Asian 6 dpi samples, infected microglia and infected excitatory neurons predominantly communicated with surrounding microglia through canonical damage-sensing and coordination pathways, most notably Apoe-(Trem2+Tyrobp) signaling. This interaction pattern is consistent with regulated microglial responses that support debris sensing, phagocytic clearance, and coordinated microglial activity during infection resolution. Notably, these infection-contained regions are precisely where DAM microglia are enriched (as shown in Section 4), suggesting that Apoe signaling from infected cells may promote DAM differentiation and function, enabling effective phagocytosis and immune resolution.

Importantly, Apoe-Trem2 signaling was disrupted in highly infected regions of AF6D samples. In these high infection-high microglia regions at 6 dpi, both Apoe ligands and the Trem2 receptor were downregulated, with substantially larger fold changes in African strain samples compared to mock controls (Fig. 5S10). This coordinated transcriptional suppression likely contributes to impaired microglial damage sensing, lipid metabolism, and phagocytic activation - processes crucial for clearing cellular debris and mounting efficient immune responses. Instead, infected inhibitory neurons and oligodendrocytes in African samples engaged microglia through diverse inflammatory, cytotoxic, and adhesion-associated signals, including Sema4d-Cd72, Tnf-Tnfr, and Gzma-F2r (Fig. 5S9). The convergence of these signals from multiple infected cell types suggests a shift toward effector-dominant microglial activity without the coordinated damage-sensing and phagocytic programs that characterize infection-contained regions.

Having identified distinct signaling environments that associate different microglial states and infection outcomes, we sought to understand the transcriptional mechanisms by which these signals establish DAM versus Inflammatory programs. We identified transcriptional regulators governing each subtype using CellOracle[30]. Inflammatory microglia are primarily regulated by Irf1, Foxl2, and Sp8, whereas Bcl3, Nfkb2, and Klf12 dominate the regulatory landscape of DAM (Fig. 5E). In Inflammatory microglia, Irf1 drives expression of pro-inflammatory genes such as Ccl5 and Ly86 (Fig. 5F), while repressing DAM-associated genes like Spp1 (Fig. 5S9B). Foxl2 promotes Ctss, a lysosomal protease essential for antigen processing, and Sp8 induces Cd72, an immune co-receptor. Together, these TFs establish a potent inflammatory state, potentially effective early in infection but prone to dysregulation if sustained, leading to prolonged neuroinflammation and tissue damage. By contrast, Bcl3 and Nfkb2 act as transcriptional repressors in the DAM subtype, downregulating inflammatory targets including Ccl5, Ctss, Cd72, and Ly86 (Fig. 5S9D), thereby counteracting Irf1-driven activation. This shift supports a more balanced state, preserving phagocytic and neuroprotective functions while restraining excessive inflammation-aligning with DAM’s role in infection resolution.

Together, these findings indicate that microglia in African strain-infected brains mount a stronger but less effective immune response, marked by prolonged inflammation and declining phagocytic function, whereas Asian strain-infected microglia exhibit a more controlled immune activation, enriched for DAM and interferon-beta signaling, potentially leading to more effective viral clearance.

### Infection-induced cellular imbalance and functional impairment of brain cell types

Having defined the microglial molecular programs underlying infection outcomes, we next examined the downstream consequences of effective versus failed immune responses on brain cellular architecture and function. Infection with the African strain produced markedly more severe neurological phenotypes (motor imbalance, weight loss, early mortality) than the Asian strain, motivating us to test whether these outcome differences track with infection-induced disruption of brain cellular architecture. We therefore quantified local changes in the densities of major structural populations across conditions and time.

Across both strains, infection progression from 4 dpi to 6 dpi was accompanied by a significant decline in the densities of oligodendrocytes (Asian: FC = 0.53, P <0.001, African: FC = 0.58, P<0.001), astrocytes (Asian: FC = 0.67, P<0.001, African: FC: 0.88, P<0.001), inhibitory neurons (Asian: FC = 0.75, P<0.001, African: FC = 0.88, P <0.001), excitatory neurons(Asian: FC = 0.87, P <0.001, African: FC: 0.87, P <0.001) (Fig. 6A-D, 1F, 2S4-5). To quantify how these coordinated density shifts alter local tissue composition, we calculated Shannon entropy of the relative cell densities of oligodendrocytes, astrocytes, inhibitory neurons, and excitatory neurons within local neighborhoods (0.08 mm²). In this context, higher entropy indicates a more even mixture of these structural cell populations, whereas lower entropy reflects compositional skew consistent with preferential depletion of one or more structural cell types. Because entropy varies across structures with different baseline cell-type compositions, we analyzed changes within each structure over time (Fig. 6E, F).

**Figure 6.**
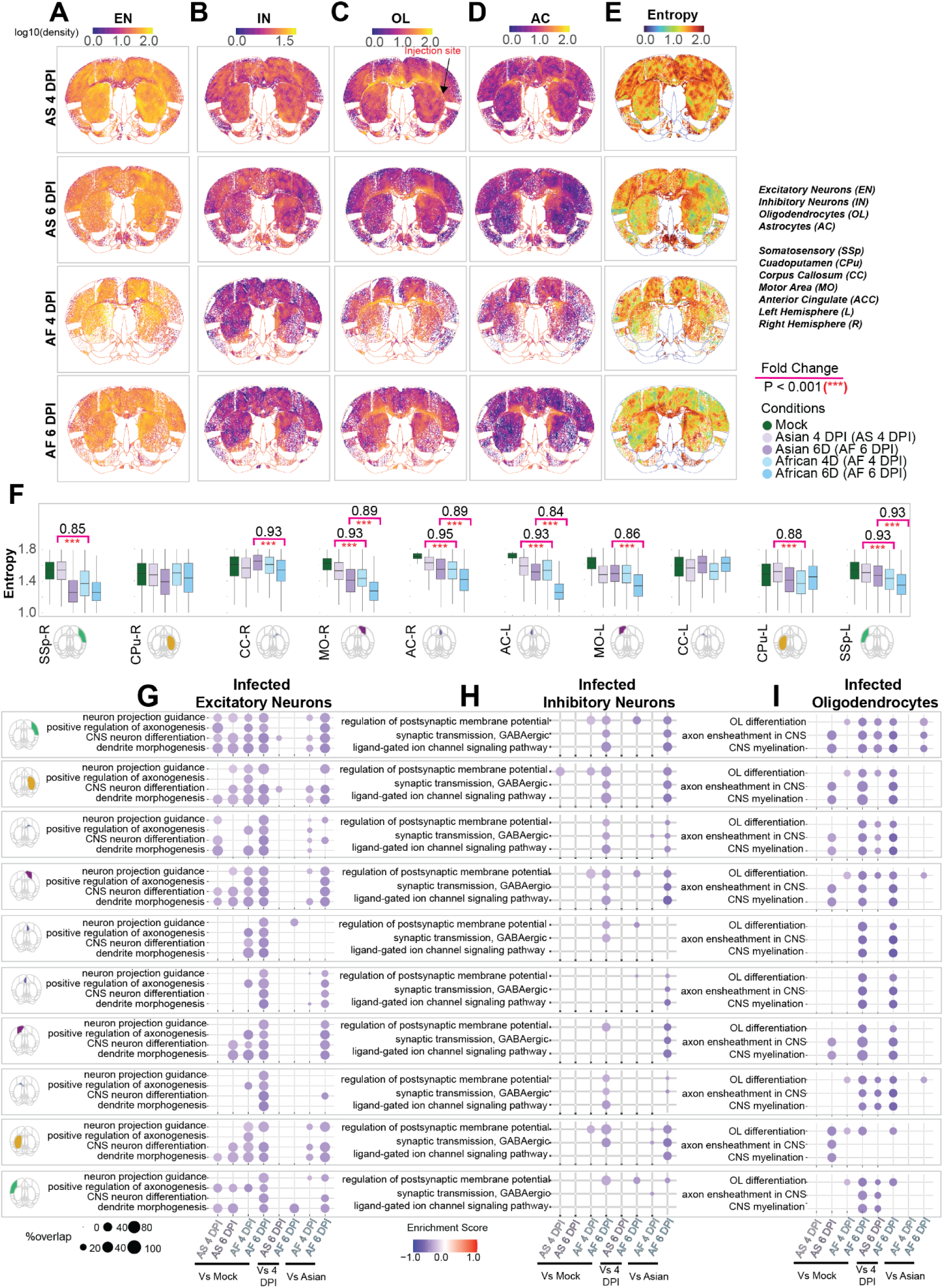
Infection-induced cellular imbalance and functional impairment of brain structural cell types. (A-D) Spatial maps of local densities for excitatory neurons **(A)**, inhibitory neurons **(B)**, oligodendrocytes **(C)**, and astrocytes **(D)** across brain sections and conditions. Local densities were calculated by counting neighboring cells of each type within 0.08 mm² neighborhoods centered on each cell, then log₁₀-transformed for visualization. **(E)** Spatial map of Shannon entropy computed from the local densities of the four major structural cell types (excitatory neurons, inhibitory neurons, oligodendrocytes, astrocytes), reflecting local cellular diversity and compositional balance. Higher entropy indicates more even cell-type distribution; lower entropy indicates compositional skew. **(F)** Distribution of local entropy values across major brain structures for mock and infected samples (Asian 4 dpi, Asian 6 dpi, African 4 dpi, African 6 dpi). Fold changes (FC) comparing African to Asian strains at matched time points are shown. Statistical significance was assessed using two-sided t-tests (***P < 0.001). **(G-I)** Bubble plots showing significantly enriched Gene Ontology Biological Process (GO:BP) terms in infected excitatory neurons **(G)**, inhibitory neurons **(H)**, and oligodendrocytes **(I)** across brain structures. Comparisons include infected versus mock, temporal comparisons (6 dpi vs 4 dpi), and strain-specific contrasts (African vs Asian). Bubble color represents enrichment direction (red: upregulated; blue: downregulated); bubble size reflects the proportion of genes within each gene set that are significantly differentially expressed (adjusted P-value < 0.05, |log₂ fold change| ≥ 0.30).

Entropy maps and region-level comparisons revealed pronounced infection-induced disruptions in local structural diversity across the brain. African samples exhibited significantly lower entropy than Asian samples at matched time points across multiple brain regions (P < 0.001, Fig. 6F), indicating widespread cellular imbalance (CC-R : FC = 0.93 at 6 dpi, MO-R: FC = 0.93 at 4 dpi; FC = 0.89 at 6 dpi, ACC-R : FC = 0.95 at 4 dpi; FC = 0.89 at 6 dpi, ACC-L : FC = 0.93 at 4 dpi; FC = 0.84 at 6 dpi, MO-L: FC = 0.89 at 6 dpi, SSp-R: FC = 0.93 at 4 dpi, 0.93 at 6 dpi). The prevalence of entropy reductions across diverse brain structures spanning both hemispheres and including cortical, subcortical, and white matter regions, indicates that African strain infection induces widespread disruption of cellular architecture throughout the brain. These reflect what has been observed in H&E staining. ZIKV infection induces progressive neuropathology in the mouse brain, with both lineages showing time-dependent neuronal injury and perivascular inflammation. The African lineage exhibits pronounced pyroptotic pathology by 6 dpi (Fig. 1S1E) and more extensive leukocyte accumulation around CNS vessels (Fig. 1S1F), whereas the Asian lineage displays comparatively milder changes, highlighting lineage-dependent differences in neurovirulence and host inflammatory response. Together, these results indicate that African strain infection is associated with a broader and deeper collapse of local structural-cell-type balance, particularly along dorsal cortical and callosal-adjacent regions, providing a tissue-architecture correlate for the more severe motor deficits and early mortality observed in African strain-infected animals.

Beyond cell loss, infection suppressed core functional programs in surviving excitatory neurons, inhibitory neurons, and oligodendrocytes, with the strongest deficits in African 6 dpi and concentrated in dorsal cortical and callosal-adjacent regions (Fig. 6G-I). In excitatory neurons, programs for neuron projection guidance, axonogenesis, CNS neuron differentiation, and dendrite morphogenesis were reduced across multiple regions, with particularly consistent suppression in MO-R, CO-R, ACC-(R/L), and SSp-(R/L) (Fig. 6G). In inhibitory neurons, synaptic transmission (GABAergic), ligand-gated ion channel signaling, and regulation of postsynaptic membrane potential were downregulated in the same dorsal regions, again most pronounced in MO-R, ACC-(R/L), and SSp-(R/L) (Fig. 6H). In oligodendrocytes, OL differentiation, axon ensheathment in CNS, and CNS myelination were broadly suppressed, with strong effects in Corpus Callosum(R/L) and adjacent motor/cingulate regions (MO-(R/L), ACC-(R/L)) (Fig. 6I). Together, the convergence of impaired neuronal connectivity, loss of inhibitory control, and myelin dysfunction provides a mechanistic framework linking infection-induced cellular failure to the severe motor imbalance, coordination deficits, and early mortality observed in African strain-infected mice.

Strain-specific transcriptional responses further highlighted greater immune dysregulation in African strain-infected brains. At AF4DPI, infected structural cells showed robust activation of innate immune, inflammatory, and apoptotic programs, with higher normalized gene ratios than Asian counterparts (Fig. 6S1-2). By AF6DPI, these pathways remained strongly elevated, particularly in inhibitory neurons and astrocytes, indicating failure to resolve early stress responses. Sustained immune activation coincided with progressive loss and functional decline of structural cell types, linking transcriptional dysregulation to impaired neuronal connectivity, disrupted glial support, and compromised myelination. Together, these observations provide a mechanistic framework connecting unresolved infection-induced stress responses to the severe motor deficits, coordination impairments, and early mortality observed in African strain-infected mice, whereas more controlled immune responses in Asian strain-infected brains preserve structural and functional integrity.

## DISCUSSION

Our study provides a spatially resolved, infection-aware framework for understanding how ZIKV strains differentially remodel the brain microenvironment. By integrating spatial transcriptomics, infection mapping, subtype-resolved microglial analysis, and structural entropy measurements, we demonstrate that viral strain-specific neuropathogenesis is not merely a function of replication rate but reflects a coordinated interplay between spatial dissemination, microglial state transitions, damage-sensing pathways, and progressive collapse of neural architecture. The divergent neurological outcomes observed between African and Asian strains arise from fundamentally different immune-structural equilibria established within defined anatomical niches. African strain infection caused widespread viral dissemination, persistent immune activation, structural cell diversity loss, and profound functional impairment, whereas Asian strain infection was relatively contained, eliciting controlled immune responses and preserving tissue integrity.

Trajectory analysis revealed that ZIKV preferentially spreads along neuroanatomical pathways, initially near the injection site and later through connected structures such as the anterior cingulate near corpus callosum regions. African strain infection propagated more rapidly and extensively than the Asian strain, consistent with its higher neuropathogenicity. Early infection broadly targeted all major cell types, but sustained infection preferentially affected inhibitory neurons, potentially disrupting excitatory-inhibitory balance and contributing to motor deficits.

We found that microglial responses are spatially compartmentalized and functionally heterogeneous. Microglia do not expand uniformly across infected tissue; instead, they form regionally restricted niches whose composition predicts infection trajectory. In Asian strain infection, early microglial recruitment in injection-proximal regions leads to the emergence of spatial domains characterized by high microglial density and declining infection burden. These regions represent effective containment zones. In contrast, African strain infection produces extensive regions in which both infection and microglial density remain high, indicating recruitment without resolution. The critical distinction is therefore not the magnitude of microglial accumulation, but the timing and qualitative state of microglial activation. This uncontrolled infection and the accompanied immune response likely amplify tissue stress and contributes to collateral damage.

Subtype analysis reveals that disease-associated microglia (DAM) play a disproportionate role in infection containment. DAM density inversely correlates with infected cell burden at the level of local microenvironments, and DAM are enriched precisely within spatial niches where infection stabilizes in Asian strain samples. These cells exhibit interferon-associated antiviral signatures while downregulating migration pathways, consistent with a locally resident, phagocytic phenotype optimized for debris clearance and immune coordination rather than inflammatory amplification. In contrast, inflammatory microglia expand proportionally with infection density but remain strongly associated with high-infection niches, particularly in African strain infection.

Persistent inflammatory activation, dominated by IRF1-driven programs, appears insufficient to restrain viral propagation and instead coincides with suppression of neuron-supportive pathways.

The spatial organization of these microglial states suggests that containment is a structured microenvironment-associated process rather than a simple antiviral response. In Asian infection, DAM accumulate early in injection-proximal somatosensory cortex, caudate-putamen, corpus callosum, and motor regions, where infection density subsequently plateaus or declines. In African infection, similar regions exhibit continued infection expansion despite microglial presence, accompanied by widespread inflammatory transcriptional activation and progressive loss of structural cell diversity. These findings indicate that DAM are not merely reactive but are mechanistically involved in reshaping the local tissue environment to favor viral restrictions.

A possible key mechanistic driver of this divergence appears to be differential engagement of the ApoE-Trem2-Tyrobp signaling axis. In infection-contained niches of Asian strain samples, infected neurons and microglia express ApoE, while microglia maintain Trem2 and Tyrobp expression, enabling canonical lipid-sensing and phagocytic signaling. ApoE-Trem2 signaling is known to regulate microglial differentiation into DAM-like states, enhance lipid metabolism, promote debris clearance, and coordinate microglial clustering [29]. Its preservation in Asian infection provides a plausible mechanism for efficient microglial recruitment and maturation into states that limit ZKV replication [31]. In contrast, African strain infection is characterized by coordinated downregulation of both ApoE ligands and Trem2 receptor expression within highly infected regions. Disruption of this axis likely impairs damage sensing and lipid-associated phagocytic programs, preventing microglia from transitioning into effective DAM states despite their numerical expansion.

This signaling divergence may be directly influenced by viral structural differences. ZIKV envelope (E) protein interacts with ApoE, and African and Asian E proteins differ in sequence and potentially in lipid-binding properties [32], which deserve further confirmation. Strain-specific modulation of ApoE availability or receptor engagement could therefore alter microglial differentiation trajectories. Such a mechanism would link viral structural distintion to spatial immune response, providing a coherent explanation for strain-dependent microglial state programming.

Microglial recruitment kinetics further distinguish infection outcomes. In Asian strain infection, microglial recruitment slopes in injection-proximal regions are steep, producing high microglia-to-infected cell ratios early in disease progression. This ratio correlates with stabilization of infection density between four and six days post-infection. In African infection, recruitment slopes are shallow relative to infection burden, resulting in persistently low microglia-to-infected ratios and continued viral expansion. These findings indicate that early, spatially concentrated recruitment is essential for containment.

The consequences of failed containment extend beyond viral burden to structural and functional collapse of neural circuits. African strain infection is associated with marked reductions in oligodendrocytes, astrocytes, and both excitatory and inhibitory neurons, accompanied by significant decreases in local entropy across multiple brain regions. Entropy reduction reflects compositional skew and preferential depletion of specific structural populations, particularly in dorsal cortical and callosal-adjacent regions. This collapse of cellular diversity is spatially aligned with regions exhibiting persistent infection and inflammatory microglial dominance. Surviving neurons show transcriptional suppression of projection guidance, synapse assembly, and postsynaptic signaling pathways, while oligodendrocytes display downregulation of myelination and axon ensheathment programs. Together, these alterations provide a mechanistic explanation for severe motor imbalance and early mortality observed in African strain-infected mice.

The divergence between strains thus reflects a shift in microglial functional equilibrium. In Asian infection, antiviral and phagocytic programs are engaged while neuron-supportive functions are relatively preserved. In African infection, inflammatory effector programs are amplified at the expense of neuro-supportive pathways, producing a trade-off that favors immune activation but compromises tissue integrity. Microglia therefore function as a critical checkpoint controlling viral containment and tissue destruction, and the balance of their transcriptional programs determines regional fate.

These findings also have broader implications for neurotropic viral disease and neurodegeneration. The resemblance of infection-associated DAM to disease-associated microglia observed in Alzheimer’s disease and other degenerative contexts suggests that common lipid-sensing and phagocytic pathways regulate tissue remodeling across diverse pathologies. Viral encephalitis may therefore exploit or disrupt conserved microglial differentiation programs, with ApoE signaling serving as a central regulatory node.

In summary, our data support a unified strain-divergent model of ZIKV neuropathogenesis in which viral structural differences influence ApoE-mediated microglial signaling, recruitment kinetics determine the timing of containment, and spatially restricted microglial state transitions govern whether infection stabilizes or progresses to structural collapse. By integrating infection mapping, spatial niche analysis, and transcriptional regulation, this study establishes a mechanistic framework linking viral strain identity to immune response and neural circuit disruption. These insights suggest that therapeutic strategies aimed at enhancing ApoE-Trem2 signaling, promoting DAM signaling, or restraining persistent inflammatory activation may shift infection trajectories toward containment and preserve neural integrity.

## METHODS

### Mice and viral infection

Four-week-old female CD-1/ICR mice (Charles River Laboratories) were intracranially (i.c.) injected with 2 μL of DMEM (mock) or DMEM containing 10L PFU of Zika virus strains MR766 (African lineage) or PRVABC59 (Asian lineage). Mice were euthanized at 4 or 6 days post-infection. Following transcranial perfusion, brains were harvested and embedded in OCT for downstream analyses.

### Intracranial injection

Mice were anesthetized with isoflurane and secured in a stereotactic frame. After confirming loss of pedal reflex, the scalp was sterilized (betadine/ethanol) and locally anesthetized with lidocaine. A ∼5 mm midline incision was made, and bregma was identified as the anatomical reference point. A burr hole was generated using a sterile 21-gauge needle at the injection site (coordinates relative to bregma: AP 1.5 mm, ML -2.0 mm). A 10 μL Hamilton syringe was inserted to a depth of 3 mm from the skull surface, and 2 μL of inoculum was delivered at a controlled rate (300 nL/min). The needle was withdrawn gradually (1μM/3 min) to minimize reflux. The incision was closed with wound clips, and mice received meloxicam (1–2 mg/kg, subcutaneous). Animals were monitored daily until endpoint.

### Study approval

All animal procedures were performed in accordance with guidelines from the National Institutes of Health and were approved by the University of Pittsburgh IACUC (Protocol #25087246).

### In-situ sequencing (ISS)

ISS libraries were prepared using the HS Library Preparation Kit (CARTANA AB, part of 10x Genomics) following the manufacturer’s protocol with minor modifications[33]. Fresh-frozen brain sections (10 μm) were fixed in formaldehyde, permeabilized with 0.1 M HCl, and dehydrated through graded ethanol. Sections were incubated with chimeric padlock probes (one custom and four predesigned panels; Supplementary Table 1) overnight at 37°C, followed by ligation at 30°C.

Rolling circle amplification products were quality-controlled using Cy5-labeled anchor probes. Samples passing QC were processed for in situ barcode sequencing and imaging. Fluorescent signals were sequentially hybridized and imaged across five channels (DAPI, Alexa Fluor 488, Cy3, Cy5, Alexa Fluor 750) using a Nikon Ti2 microscope (20× objective; Zyla 4.2 camera). Z-stacks were projected by maximum intensity projection. Six sequencing cycles enabled full barcode decoding.

### Immunofluorescence (IF)

Fresh-frozen (FF) brain tissues were cryosectioned at 10 μm thickness, mounted on Superfrost slides, and stored at −80°C until use. Sections were air-dried and fixed in 4% paraformaldehyde for 15 min at room temperature, followed by permeabilization with 0.1% Triton X-100 in PBS. After blocking with 5% bovine serum albumin (BSA) in TBST for 1 h at room temperature. Zika NS1 antibody (GeneTex, Cat. No. GTX133307) were incubated at 25 μg/ml overnight at 4°C. Following TBST washes, sections were incubated with Goat anti-Rabbit IgG (H+L) Highly Cross-Adsorbed Secondary Antibody, Alexa Fluor™ Plus 647 (Invitrogen, Cat. No. A32733) for 1 h at room temperature in the dark. Nuclei were counterstained with DAPI. Slides were mounted using antifade mounting medium and imaged using Olympus IX83 Microscope with cellSens Software.

### Quantitative real-time PCR (RT–qPCR)

Total RNA was extracted from mosue brain TRI reagent (Sigma, Cat. No. T9424) according to the manufacturer’s instructions. RNA (1 μg) was reverse transcribed into cDNA using Maxima H Minus First Strand cDNA Synthesis Kit (Thermo Fisher, cat. no: K1652). Quantitative PCR (qPCR) was performed using SsoAdvanced™ Universal SYBR® Green Supermix Kit (Bio-Rad, cat. no: 172-5272). Reactions were carried out in technical triplicates using Zika NS1 primers (Forward: 5’-CCATACGGCCAACAAAGAGT-3’, reverse: 5’-CAGCTCCTTCCATAGCCAAG-3’) under standard cycling conditions. Gene expression levels were normalized to mATCB (Forward: 5’-CCCTGAAGTACCCCATTGAA-3’, Reversed: 5’-GGGGTGTTGAAGGTCTCAAA- 3’), and relative expression was calculated using the ΔΔCt method.

### Cell segmentation

Cell segmentation was performed by running Baysor on the spatially resolved reads [1]. A nuclei mask prior was also generated for Baysor by segmenting DAPI images that correspond to each sample. The nuclei in the DAPI images were segmented with a fine-tuned Cellpose 2.0 model[34]. Instances where the model performed poorly were detected by performing a global Otsu thresholding on the DAPI signal that remained after removing the existing nuclei masks[35]. The centroids remaining Otsu masks were clustered with HDBSCAN to identify poor performing regions and Cellpose was rerun within the problematic regions to bolster the segmentation accuracy[36].

The Baysor cell segmentation was executed as a two-step process. First, Baysor was run on all reads, and the nuclei mask image was used as a prior. Baysor heavily favors reads that align with the nuclei mask when assigning cells, so many valid cells were erroneously classified as noise by Baysor after the first run. Thus, a second run was executed, wherein all reads classified as noise were input into Baysor along with the nuclei mask image. The cells identified by both runs were collated into a spatial single cell table.

### Quantification of infection intensity

Zika infection signals were obtained from infection-stained images. First the foreground is segmented with Otsu thresholding, then a second round of thresholding is performed within the foreground to identify the zika signal. The zika intensity can vary across the whole-slide images, which can result in incomplete segmentation. To combat this issue, the centroids of all the segmentation masks identified via the thresholding are taken and clustered with HSBSCAN. Then Otsu thresholding is performed within each group of masks identified by the clustering. This results in a more specific mask for the zika intensity signals.

A statistical test is leveraged to identify cells that are positive for infection signals. Within each sample, all cells with 2% or less overlap between the cell mask and the zika infection mask are grouped together. From this group 60% of the cells are sampled to create an infection negative distribution. A one sample t-test is performed to evaluate whether the mean zika intensity for each cell in a sample is greater than the mean intensities from the infection negative distribution. The p-values for all cells are corrected with Benjamini-Hochberg correction and all cells with a p-value less than 0.001 are considered to be infected.

### Quality control

To ensure high-quality data for downstream analysis, we applied filtering criteria to both spatial transcriptomics and single-cell RNA-seq datasets. For the spatial transcriptomics data, we exclude cells with fewer than 5 total transcript counts or with fewer than 4 detected genes, as such cells are likely to represent low-quality captures or ambient signals. For the single-cell RNA-seq dataset, we filtered out cells with fewer than 100 total counts, fewer than 75 expressed genes, or with a mitochondrial gene expression percentage greater than 20%. These thresholds were selected to remove low-quality cells, which typically exhibit reduced transcriptomic complexity and elevated mitochondrial content, indicative of compromised cell integrity. This quality control step ensured that downstream analyses were based on biologically meaningful and technically robust cellular profiles.

### Cell typing and Data integration

To obtain a refined and data-centric list of cell type marker genes, we employed a two-step approach.

First, we used a likelihood-based Bayesian classifier to predict cell types in the spatial transcriptomics dataset (query) with high confidence (posterior probability ≥ 0.90). For this, we applied the supervised classification functionality of the *Insitutype* package[37], training the model on a reference single-cell RNA-seq dataset [38]. Second, we identified significant marker genes for each predicted cell type by differential expression analysis using the FindAllMarkers function from the *Seurat* package, selecting genes with adjusted p-values < 0.05 and absolute log₂ fold-change ≥ 0.5. These marker genes were then utilized in a subsequent cluster-based cell typing approach.

For cluster-level cell type assignment, we employed an AUC-based method described below. Separately, to integrate the single-cell RNA-seq data with the spatial transcriptomics data, we used the *LIGER* framework[11]. Integration served two purposes: (1) to leverage the higher sequencing depth of the single-cell data to potentially improve clustering in the spatial data, which inherently suffers from high sparsity[39]; and (2) to enable transcriptome-wide gene imputation (described later). We optimized the number of non-negative matrix factorization (NMF) factors (K) used in *LIGER* by maximizing the number of unique high-confidence cell types (AUC score > 0.60), empirically testing K values in the range of 10 to 45.

The AUC-based cluster cell typing process was conducted as follows. Let ***Q*** ∈ ℝ*^nxp^* denote the ‘SCTtransform’ normalized spatial gene expression matrix, where *n* is the number of cells and *p* is the number of spatially resolved genes. For each predefined cell type *c*, associated with a marker gene set *M*_(*C*)_ ⊆ = {1,…,.p} we computed a per-cell score by summing the expression values across genes in *M*_(*C*)_. This resulted in a score matrix ***S*** ∈ ℝ*^nxm^* where *m* is the number of cell types. For each cluster *k* ∈ {1,…,.*K*} and each cell type *c*, we evaluated the ability of *S·,c* to distinguish cells belonging to cluster *k* from other cells using the Area Under the Receiver Operating Characteristic Curve (AUC), generating an AUC matrix ***A*** ∈ ℝ*^kxm^* Finally, we assigned each cluster the cell type with the highest AUC score, meaning the one that best matched the expression profile of that cluster.

### Local cell density and microenvironment definition

To quantify local cell-type-specific or infected cell density in spatial transcriptomics data, we calculated the number of cells of interest within a fixed-radius neighborhood around each spatial cell. Specifically, we defined a circular area of 0.08 mm² centered on each cell’s spatial coordinates and counted the number of neighboring cells that matched the target identity (e.g., a specific cell type or infection status). This yielded a per-cell local density score.

Because absolute cell density can vary across brain samples due to technical variability (e.g., differences in tissue section thickness, capture efficiency, or imaging resolution) rather than biological differences, we normalized the density scores within each brain sample to facilitate cross-sample comparison. For each sample, the raw density scores were divided by the sample’s mean cell density and scaled by a factor of 100, resulting in normalized density scores expressed as relative percentages. Infected cell density was calculated using the same procedure, treating infection-positive cells as the target population. This approach minimizes the influence of sample-specific artifacts and allows for more accurate biological interpretation of spatial cell distribution and microenvironment structure.

#### Statistical analysis of distributional differences

To assess statistical differences in spatial features (e.g., infected cell density or cell type density) between experimental groups, we performed a resampling-based *t*-test approach. For each comparison, such as for example, infection density between Asian 6DPI and 4DPI samples in the somatosensory cortex-we randomly sampled 100 values from each distribution and performed an unpaired two-sided *t*-test. This process was repeated 10,000 times to robustly estimate variability in the test statistics. Resampling was necessary for several reasons. First, conventional *t*-tests can yield artificially low *p*-values in large datasets due to their high sensitivity, especially when sample sizes are large, as even minor differences can become statistically significant regardless of biological relevance. Second, spatial transcriptomics data often violates the assumptions of the *t*-test, such as normality and equal variances. Resampling avoids relying on these assumptions and yields more robust and interpretable results. Third, this approach accounts for local variability and heterogeneity in the data, reducing the influence of outliers or sampling bias. Lastly, it enables estimation of a distribution of effect sizes, rather than relying on a single test outcome, allowing for a more reliable assessment of consistent biological effects.

To determine statistical significance, we computed empirical *p*-values[40] by comparing the observed *t*-statistic (from a single *t*-test on the full sampled distributions) to the distribution of test statistics generated from the resampling procedure. Specifically, we calculated the proportion of resampled *t*-statistics whose absolute value was greater than or equal to that of the observed test statistic. This empirical approach controls potential violations of *t*-test assumptions (e.g., non-normality, unequal variances) and accounts for variability due to subsampling.

### Brain atlas registration and anatomical structure transfer

To assign anatomically accurate brain region labels to spatial transcriptomics sections, we used a two-step image registration workflow involving *QuickNII* followed by *VisuAlign[17]*, allowing both affine and nonlinear adjustments relative to the Allen Mouse Brain Common Coordinate Framework (CCF v3)[41].

#### Generation of Customized Spatial Transcriptomics Tissue Images

Prior to registration, we generated customized spatial transcriptomics tissue images to enhance anatomical landmark visibility and improve atlas alignment accuracy. For each brain sample, we identified globally significant spatially variable genes using the moran.test function from the *rgeoda* package. This function computes the global Moran’s I statistic for each gene to assess spatial autocorrelation. Genes with *p*-value < 0.001 and mean expression > 0.01 were selected as spatially informative. The expression matrix of these selected genes was then used for spatial clustering. We applied the *Seurat* pipeline using RunPCA (npcs = 5), followed by FindNeighbors (dims = 1:5) and FindClusters (resolution = 0.08). These cluster labels and gene expression features were used to generate customized images that captured prominent spatial transcriptional patterns, improving the interpretability of anatomical boundaries during registration.

#### Affine and Nonlinear Atlas Alignment

In the first registration step, high-resolution customized spatial transcriptomics images were loaded into *QuickNII*, and the Allen Mouse Brain CCF v3 was selected as the reference atlas. Each tissue section was manually aligned to the atlas in 2D space by adjusting translation, rotation, and scaling to best match anatomical landmarks (e.g., corpus callosum, hippocampus). This affine alignment allowed us to approximately position the section within the coordinate system of the atlas.

In the second step, we used *VisuAlign* for nonlinear registration refinement. The affine transformation from *QuickNII* was imported into *VisuAlign*, and a grid of control points was manually placed on anatomical features visible in the tissue section. These control points were used to locally deform the atlas overlay using elastic warping, allowing finer alignment of atlas borders to curved or distorted anatomical structures that could not be fully aligned by affine transformation alone.

The final warped atlas annotations were exported and used to assign anatomical labels to each spatially barcoded region.

### Mapping brain samples to a common coordinate system

(Used in visualization and simulation purposes)

To enable anatomically consistent and spatially aligned comparisons across brain samples, we mapped each registered brain section to a common coordinate system. While tools such as *QuickNII* and *VisuAlign* were used to achieve high-precision registration of individual brain samples to the Allen Mouse Brain Common Coordinate Framework (CCF v3), the resulting aligned sections can still vary across samples due to differences in slicing angle, and tissue deformation. These variations affect the spatial orientation of structures and can hinder direct comparison across brain samples. To address this, we implemented a standardization strategy that maps cells from each brain onto a unified spatial grid based on a shared reference atlas section. This approach ensures consistent structure annotation and facilitates accurate, comparative spatial analyses across brains, even when the underlying reference slice differs slightly due to individual registration outcomes.

We began by selecting a representative reference brain section from the Allen CCF that closely matched the morphology of all 10 ZIKV infected experimental samples. For each anatomical structure within this reference brain, we subdivided the region into a grid system composed of equally sized spatial units. Each grid was indexed along two spatial dimensions: the x-axis (left to right) and the y-axis (bottom to top), creating a two-dimensional matrix of spatial identifiers.

Each corresponding anatomical structure in the experimental (query) brain samples was divided into the same number of spatial grids as in the reference structure, using an identical x–y indexing scheme. This ensured that cells within each grid of a query brain could be accurately mapped to the corresponding spatial grid in the reference brain, enabling consistent spatial alignment across samples.

To project cells from each query brain into the reference space, we first normalized the location of each cell within its local grid by rescaling its position to a 0-to-1 range along both spatial axes. These normalized coordinates captured the relative position of a cell within its own grid cell, independent of absolute dimensions. To map these cells into the reference brain, we then linearly scaled the normalized coordinates to the actual spatial extent of the corresponding reference grid cell. Specifically, each normalized x and y coordinate was transformed using the minimum and maximum boundaries of the reference grid cell along each axis, effectively stretching or shrinking the relative position to match the physical dimensions of the reference grid. This linear scaling process preserves the internal spatial organization of each grid while ensuring that cells from all query brains are accurately repositioned within a shared anatomical framework.

This grid-based projection strategy enabled robust, anatomically informed spatial registration across samples, ensuring that each cell was placed in a structurally and spatially comparable context. This facilitated more reliable analyses of cell-type distributions, density estimates, and region-specific spatial patterns across the dataset.

### Trajectory analysis of infection dissemination in the brain

To analyze how infection disseminates through brain structures over time, we designed a trajectory inference method that combines spatial and density-based metrics across multiple samples. This approach was inspired by previous work[42], and subsequently adapted and customized to fit the specific requirements of our brain infection model.

#### Root Node Selection

We began by identifying the root node, defined as the initial site of infection dissemination. For each brain structure, we computed the average infection density across all samples during early pseudo-time points (1 to 3). The structure with the highest average infection density was selected as the root node (R). The selected root node is also spatially close to the injection site.

#### Pairwise Distance Computation

For each brain sample and each pair of brain structures *(a, b)*, we computed a total distance between the two structures, defined as:

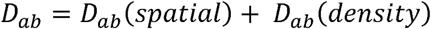

where:

- *D_ab_* (*spatial*) is the spatial distance between the infection epicenters in a and b
- *D_ab_* (*density*) is the infection density difference between a and b

#### Spatial Distance

To identify infection epicenters, we used DBSCAN clustering (with parameters: ε *= 0.5*, *min_samples = 10*) on the infection intensity values of individual cells within each structure.

Let H_a_ = {h_a1_, h_a2_, …, h_am’_} and H_b_ = {h_b1_, h_b2_, …, h_bn’_} represent subclusters (infection epicenters) of structures a and b, respectively. We then computed the average Euclidean distance between all pairs of subcluster centroids across H**_a_** and H_b_ as *D_ab_* (*spatial*).

### Infection Density Difference

We computed the infection density difference as the Euclidean distance between the average infection densities of the subclusters:

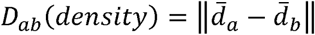

where *d̄*_*a*_ and *d̄*_*b*_ are vectors of mean infection densities for subclusters within **a** and **b**, respectively.

#### Total Distance Across Samples

We performed the above calculations for all 10 brain samples, obtaining a set of distances {D_ab_(o)}, where *o* indexes the sample. The final pairwise distance was computed as:

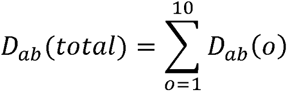

#### Graph Construction and Trajectory Inference

Using all structures with non-zero infection density and the matrix of distances *D_ab_* (*total*), we constructed a directed graph G = (V, E), where: V is the set of brain structures, E is the set of directed edges with weights *D_ab_* (*total*).

To identify the most likely dissemination paths, we computed the minimum spanning arborescence of this graph, rooted at the previously identified node R. This yielded an optimal tree structure representing the most likely directional flow of infection through the brain.

### Whole-transcriptome gene imputation in spatial cells

To impute full transcriptome gene expression for spatial transcriptomics cells, we first integrated the spatial transcriptomics and single-cell RNA-seq datasets using the *LIGER* package (as described earlier). We then computed the 2D UMAP coordinates on ‘integrated dataset’ using the *Seurat* pipeline. Let X^st^ ∈ ℝ^m×2^ represent the UMAP coordinates of *n* spatial transcriptomics cells, and X^wt^ ∈ ℝ^r×2^ those of *r* single-cell RNA-seq cells.

For each target spatial cell *i*, we identified its q nearest neighbors in X^wt^ based on Euclidean distance in UMAP space, with *q* set to 100. To ensure high-quality imputation, we applied a stringent filtering strategy to retain only biologically relevant neighbors. Specifically, candidate neighbors were required to: (a) belong to the same cluster as the target cell, (b) originate from the same experimental condition (e.g., tissue source or cohort), and (c) exhibit a Pearson correlation coefficient ≥ 0.50 with the target cell, calculated over the expression values of the 218 shared genes used in the LIGER embedding.

Let **N_i_** denote the set of single-cell neighbors that satisfied all filtering criteria for spatial cell *i*. Using ‘SCTransform’ normalized gene expression of single-cell RNA-seq dataset is *Z* ∈ ℝ*^r×p^’*, where *p’* is the number of genes in *Z*. The imputed value, *z̑*_*ig*_ for gene g in spatial cell i is:

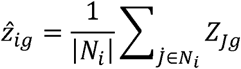

This procedure was repeated for all spatial cells to generate a complete imputed transcriptome matrix for the spatial dataset.

#### Bivariate spatial autocorrelation analysis (Moran’s I)

To assess the spatial relationship between infected cells and microglia density, we performed a Bivariate Moran’s I analysis using the local_bimoran function from the *rgeoda* R package. This method quantifies the local spatial correlation between two spatial variables-in our case, the density of infected cells and the density of microglia-by evaluating how the value of one variable at a location relates to the spatial lag (average in the neighborhood) of the second variable. Spatial coordinates for each cell were provided to the model to preserve spatial context.

The local_bimoran function returns a classification of each spatial location into one of several categories: “High-High” (both variables high, spatially clustered), “Low-Low” (both variables low, clustered), “Low-High” or “High-Low” (spatial outliers), “Not significant” (no meaningful local spatial autocorrelation), “Isolated” (cells with no spatial neighbors), and “Undefined” (ambiguous or missing data). It also provides corresponding *p*-values to assess statistical significance for each region. In this study we used p<0.01 as significant region.

This approach allows for localized, fine-grained analysis of co-enrichment patterns, enabling us to identify potential spatial interactions or exclusion zones between infected cells and microglia. Such analysis is critical for understanding spatial microenvironmental dynamics in tissue contexts like the brain, where spatial organization of immune and infected cells may influence disease progression.

### Stochastic simulation of infection dynamics

To explore infection-microglia interactions beyond the experimentally measured time points, we modeled infection dynamics using stocastic, a density based framework inspired by Reed-Frost chain-binomial model [24]. The framework extends the Reed–Frost chain-binomial approach to a 2D spatial lattice in which cell-state and microglial transitions are sampled from binomial distributions at each discrete time step.

The model tracks four spatially indexed variables - susceptible (S), infected (I), recovered (R), and microglia (G) - across D spatial units and T time steps. The cell balance S(t,d) + I(t,d) + R(t,d) = N_c is preserved at every site, and microglia are bounded by n_g ≤ G(t,d) ≤ N_g (N_c = 200, N_g = 120, n_g = 20). Brain coordinates were mapped to a 168 × 104 grid (∼0.005 mm² / grid), with adjacency defined as A_0(i, j) = 1 when Euclidean distance ≤ 9 (Euclidean radius 3 grid units); the row-normalized matrix A was used in all subsequent updates.

#### SIR dynamics

New infections were drawn as I_new (t+1, d) ∼ Binomial(S(t,d), p_i), with p_i = 1 − exp(−β · max{0, A·I(t)/N_c − γ · A·G(t)/N_g}). The β term controls virulence, γ controls microglial dampening, and the maximum imposes a deterministic threshold below which local microglial recruitment fully suppresses spread. Recoveries were drawn as R_new(t+1, d) ∼ Binomial(I(t,d), ρ · G(t)/N_g), with ρ the recovery rate. States were updated as I(t+1) = I(t) + I_new − R_new, R(t+1) = R(t) + R_new, and S(t+1) = S(t) − I_new.

#### Microglia dynamics

Microglia were updated through three sequential mechanisms. Proliferation was triggered when a lagged local infection signal exceeded a threshold Δ_prol = 0.2: G_prol ∼ Binomial(min{G, N_g − G}, κ · H(Ī − Δ_prol)), where H is the Heaviside step function, κ = 0.6, and Ī is the infection signal averaged over a lag window (δ₁ = 2, δ₂ = 4). Chemotaxis migrated microglia from the intermediate state G¹ = G + G_prol to neighbors via a multinomial draw weighted by the smoothed infection profile A·I(t); migrated cells were treated as duplicates of their source sites to prevent local depletion. Deactivation removed microglia where lagged infection had cleared: G_deact ∼ Binomial(G², μ · max{0, 1 − Ī}) with μ = 0.1. The final state was clamped to [n_g, N_g].

#### Initialization and parameter regimes

Simulations were seeded at the CPu-R, SSp-R intersection (grid coordinate (122, 51)) within an elliptical region (rotated by θ = −120°; eccentricities a₁ = 1/2, a₂ = 10), with initial infection intensities drawn independently from 0.8 · Beta(0.1, 0.3) · N_c. Shared parameters (N_c, N_g, n_g, δ₁, δ₂, Δ_prol, κ, μ) were fixed across strains. Strain-specific parameters (β, γ, ρ) were tuned by comparing simulated outputs to the observed infection density maps and microglia density maps at 4 and 6 dpi for Asian and African strains, adjusting parameters until simulated maps qualitatively matched experimental data. Final parameter values: Asian - β = 10, γ = 0.55, ρ = 0.10; African - β = 20, γ = 0.05, ρ = 0.01 (Figure 3S2). For sensitivity analyses (Figure 3G), β, γ, and ρ were systematically varied over β ∈ {1-20}, γ ∈ {0.05, 0.1, 0.4, 0.55}, and ρ ∈ {0.001, 0.01, 0.05, 0.1}; T = 50 simulation cycles were mapped to the 0-6 dpi experimental scale.

### Microglia subtype identification

To identify transcriptionally distinct subtypes of microglia, we analyzed a total of 72,251 spatially resolved microglia cells collected from two mock controls and ten ZIKV-infected brain samples. Clustering was performed using dimensionality reduction and neighborhood-based graph clustering on the expression profiles of 218 spatially resolved genes. Principal component analysis (PCA) was first applied using all selected genes to capture the major axes of transcriptional variability. The top five principal components were then used to construct a k-nearest neighbor (KNN) graph with FindNeighbors(dims = 1:5, reduction = “pca”), representing local transcriptional similarity among cells.

Clustering was performed using FindClusters(resolution = 0.06), a graph-based community detection method that identifies discrete cell populations. The resolution parameter was set to 0.06 to balance cluster granularity and biological interpretability. A lower resolution was selected to avoid over-fragmentation of closely related microglial states, ensuring stable subtype definitions. Uniform Manifold Approximation and Projection (UMAP) was used for visualization of the identified clusters using the same five PCA dimensions (RunUMAP(dims = 1:5)).

To validate and biologically characterize the resulting clusters, we leveraged imputed whole-transcriptome gene expression (described earlier) to examine known and candidate marker genes[27]. This enabled robust functional interpretation of microglial subtypes, including infection-induced activation states, that might not be fully distinguishable using only the spatially resolved gene set.

### Differential expression analysis and pathway enrichment

Differential expression analysis was performed on imputed transcriptome data using Seurat’s FindMarkers function with the MAST hurdle model via test.use = “MAST”; all other parameters were left at default values. Gene Set Enrichment Analysis (GSEA) was performed using the gseGO function from the clusterProfiler R package[43] against the Gene Ontology Biological Process (GO:BP) collection, using the org.Mm.eg.db annotation database for mouse gene identifiers. Genes were ranked by their log₂ fold change values from the MAST differential expression analysis (highest to lowest). Analyses were run with the following parameters: ont = “BP”, minGSSize = 20, maxGSSize = 500, and pvalueCutoff = 0.05; all other parameters were left at default values.

### Spatial cell-cell communication analysis

To investigate spatially resolved intercellular communication, we performed *CellChat* analysis[44] on the spatial transcriptomics data for each brain sample. Analyses were conducted not only at the whole-sample level but also within specific anatomical structures and infection–microglia microenvironments. These spatial niches-such as “low infection–high microglia” and “high infection–high microglia” regions-were identified using Bivariate Moran’s I spatial autocorrelation analysis, enabling biologically focused interrogation of local cell-cell signaling dynamics.

We used the *spatial* module of *CellChat*, which incorporates the physical location of cells to infer spatially constrained communication networks by identifying neighboring cells based on distance. We configured the spatial.factors with a conversion.factor of 0.16 and a spot.size of 10. The conversion.factor adjusts the coordinate system of the spatial dataset to match micrometer-scale distances, ensuring accurate modeling of spatial proximity, while spot.size estimates the physical diameter of a cell or capture spot, influencing neighborhood detection.

Cell-cell communication probabilities were computed using the computeCommunProb function with the following parameters: type = “truncatedMean” and trim = 0.05 to robustly average expression values by removing the top and bottom 5% of outliers, reducing the influence of extreme values. The option distance.use = TRUE enabled spatial distance-based weighting, and interaction.range = 250 restricted inferred interactions to a maximum radius of 250 microns. The parameter scale.distance = 1 preserved original spatial distances without rescaling. To prioritize spatially localized and potentially contact-mediated interactions, we enabled contact.dependent = TRUE with contact.range = 10, limiting physical contact-based interactions to cells within 10 microns.

Finally, we applied filterCommunication(cellchat, min.cells = 10) to remove inferred interactions involving fewer than 10 cells per interacting group, ensuring the reliability of the inferred signaling pathways. For visualization, we selected ligand–receptor pairs with a communication probability > 0.0005 and a p-value < 0.05. To highlight condition-specific signaling patterns, we further restricted visualization to L–R pairs that were consistently identified across all samples within a given condition (e.g., Asian 6DPI or African 6DPI).

### Gene regulatory network inference

To reconstruct gene regulatory networks (GRNs) in spatial microglia, we applied *CellOracle* analysis[30] independently to each microglial subtype. This allowed us to capture subtype-specific regulatory programs with spatial resolution. As input, we used the full transcriptome expression matrix derived from our previously described imputation strategy, which enabled comprehensive modeling of transcriptional regulation beyond the limited number of directly measured genes in spatial transcriptomics.

Since we did not generate single-cell ATAC-seq data for these samples, we initialized *CellOracle* using a predefined base GRN derived from a mouse scATAC-seq atlas, loaded via co.data.load_mouse_scATAC_atlas_base_GRN(). This base GRN provides a curated prior regulatory structure, allowing CellOracle to perform regression-based GRN inference grounded in biologically plausible transcription factor–target relationships.

We selected highly variable genes using the sc.pp.filter_genes_dispersion function from the *Scanpy* package with parameters flavor = ‘cell_ranger’, n_top_genes = 3000, and log = False. This method mimics the gene selection strategy used in 10x Genomics pipelines, identifying genes with high expression variability, which improves the accuracy and interpretability of GRN modeling.

GRNs were inferred using oracle.get_links(cluster_name_for_GRN_unit = “micro_subtype”, alpha = 10, verbose_level = 10), ensuring that each microglial subtype was analyzed independently. The alpha parameter controls model sparsity, with higher values promoting simpler, more stable networks. verbose_level = 10 enabled detailed logging for monitoring model convergence and output.

To retain only high-confidence interactions, we applied links.filter_links(p = 0.001, weight = “coef_abs”, threshold_number = 2000). This step filters edges based on statistical significance (*p* < 0.001), ranks them by the absolute value of regression coefficients (indicating regulatory influence), and retains the top 2000 interactions per subtype.

### Lead contact

Further information and requests for resources and reagents should be directed to and will be fulfilled by the lead contact, Shou-Jiang Gao.

### Data and code availability

- Source code is available at https://github.com/qsl734/zika_spatial_analysis. and processed data archived at Zenodo: https://doi.org/10.5281/zenodo.19685781
- Any additional information required to reanalyze the data reported in this paper is available from the lead contact upon request.

## ACKNOWLEDGMENTS

We thank members of Drs. Shou-Jiang Gao and Yufei Huang laboratories for technical assistance and discussions. This study was supported by grants from the National Institutes of Health (CA096512, CA284554, CA278812, CA291244 and CA124332 to S.-J. Gao; U01CA279618 and R21GM155774 to Y. Huang), UPMC Hillman Cancer Center Startup Funds to S.-J. Gao and Y. Huang, and in part by award P30CA047904. This research was also supported in part by the University of Pittsburgh Center for Research Computing, RRID:SCR_022735. Specifically, this work used the HTC cluster, which is supported by S10OD028483.

## AUTHOR CONTRIBUTIONS

Conceptualization, S.-J.G.; methodology, M.H., W.M, G.M, H.H, Y.H.; Investigation, M.H., W.M, Y.H., S.-J.G.; writing-original draft: M.H., W.M, Y.H., S.-J.G., writing-review & editing, M.H., W.M, H.H., Y.H., S.-J.G.; funding acquisition, Y.H., S.-J.G.; resources, Y.H., S.-J.G.; supervision, Y.H., S.-J.G.

## SUPPLEMENTAL INFORMATION

**Figure 1S1.**
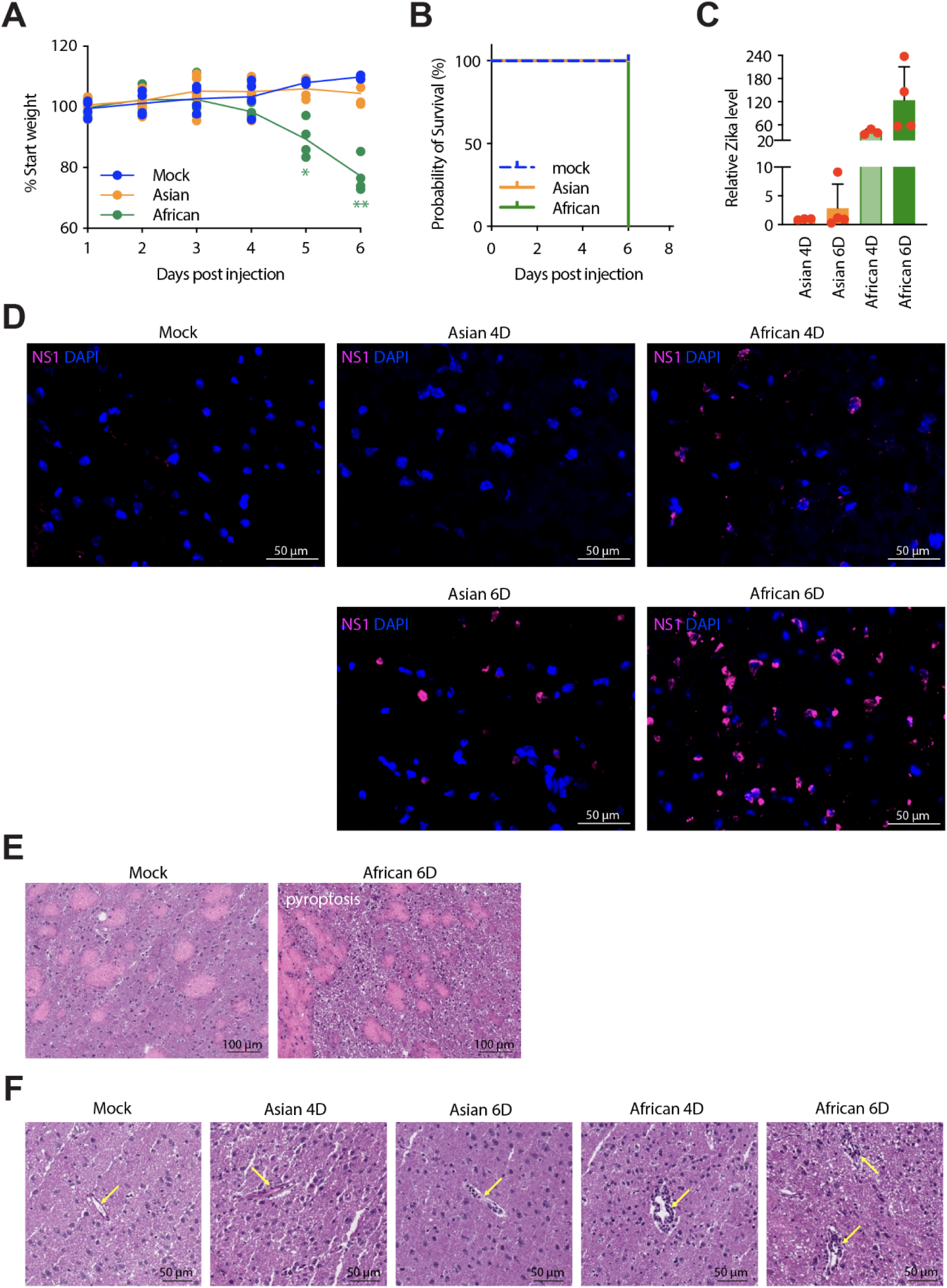
African-lineage Zika virus exhibits enhanced pathogenicity compared to Asian-lineage virus *in vivo.* (A) Percent starting body weight following intracranial infection with mock, Asian-lineage (PRVABC59), or African-lineage (MR766) Zika virus. (B) Kaplan–Meier survival analysis. (C) Relative viral RNA levels at 4 and 6 dpi. (D) Representative H&E-stained brain sections from mock and African-lineage–infected mice at 6 dpi, showing extensive tissue damage. (E) Representative H&E images across indicated conditions; arrows pointing at blood vessels in the CNS parenchyma, extensive infiltration of leukocytes seen in ZIKV infected mice, especially at 6 dpi. (F) Immunofluorescence staining for NS1 (magenta) and DAPI (blue), showing increased viral antigen in African-lineage infection.

**Figure 1S2.**
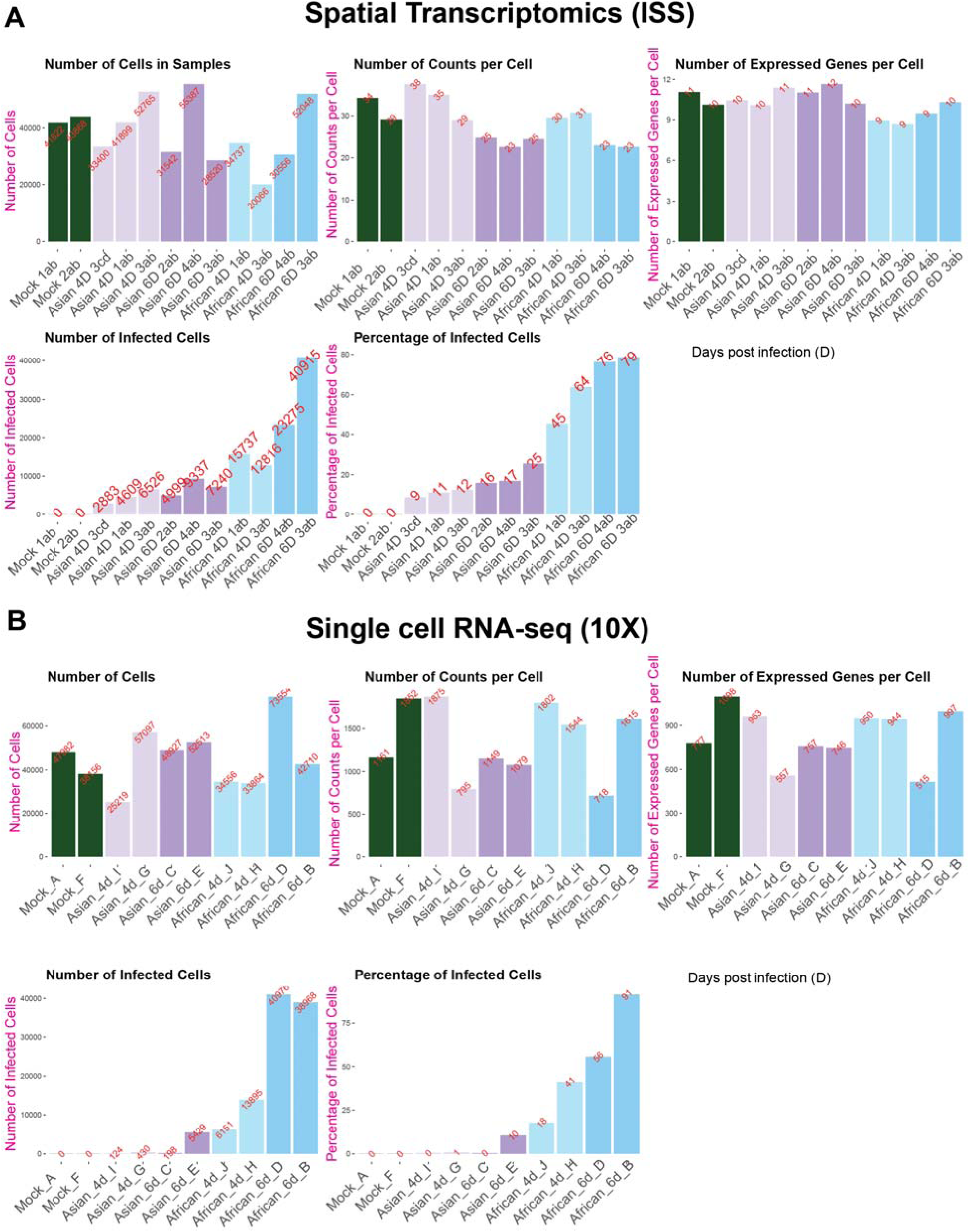
Data quality and infection characteristics of spatial transcriptomics and scRNA-seq datasets. **(A)** Per-sample quality metrics for spatial transcriptomics data across all experimental conditions (mock, Asian 4 dpi, Asian 6 dpi, African 4 dpi, African 6 dpi). Metrics include: total number of spatially resolved cells per sample (range: 20,066-55,387; median: 38,279.5), average transcript counts per cell, average number of detected genes per cell (from 218-gene panel), number of infected cells, and percentage of infected cells (infection rate: 9-79%). **(B)** Corresponding per-sample quality metrics for scRNA-seq data. Metrics include: total number of captured cells per sample (range: 25,219-73,554), average transcript counts per cell, average number of detected genes per cell (range: 515-1,098 genes), number of infected cells, and percentage of infected cells (infection rate: 1-91%). Together, these metrics demonstrate robust data quality across both platforms and highlight their complementary strengths: spatial transcriptomics provides spatial resolution with moderate transcriptomic depth (218 genes), while scRNA-seq provides deep transcriptomic profiling (whole transcriptome) without spatial information, justifying their integration for comprehensive analysis.

**Figure 1S3.**
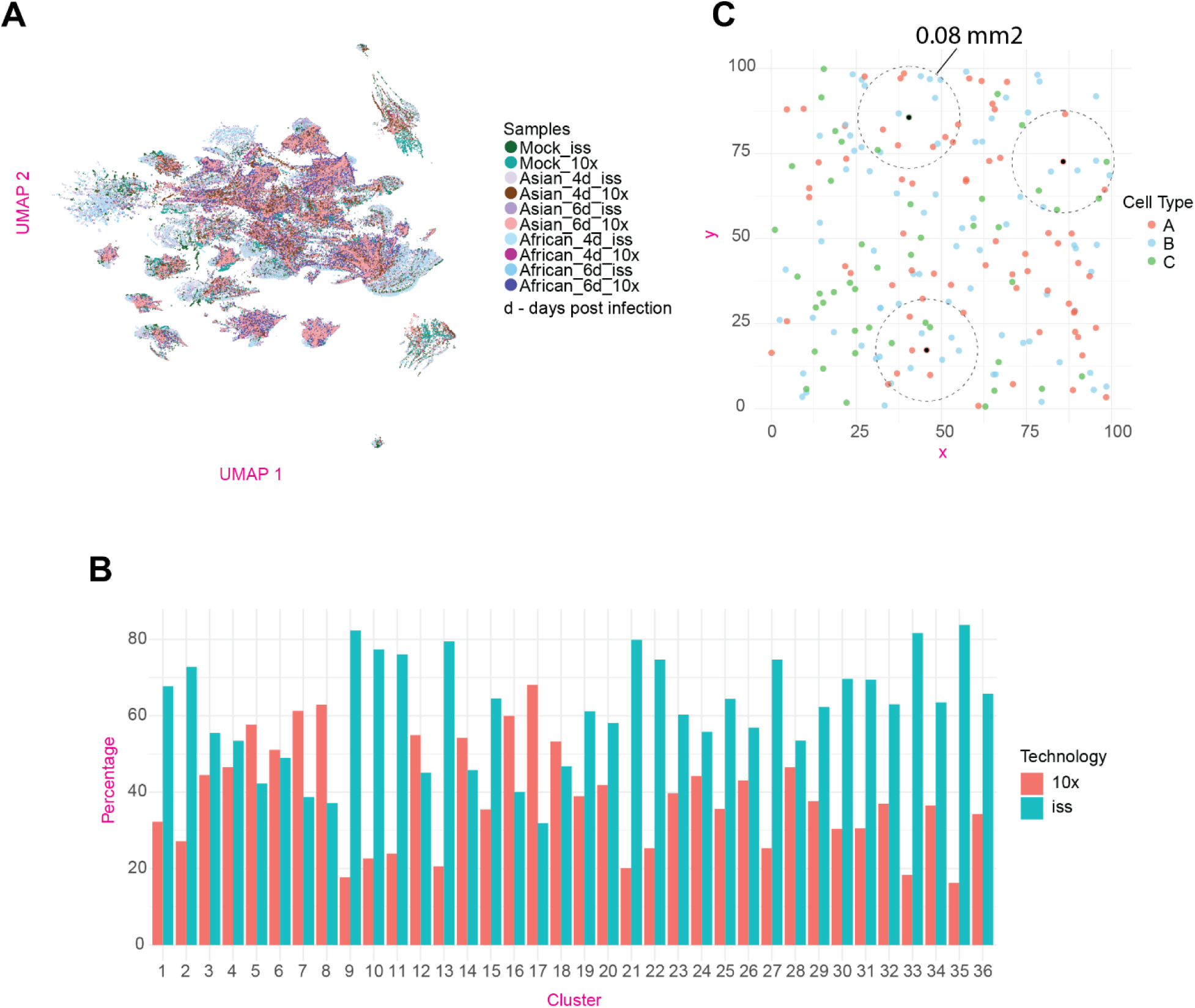
Cross-modality integration of spatial transcriptomics and scRNA-seq data and computation of local cell density. **(A)** Joint UMAP embedding of spatial transcriptomics (ISS) and scRNA-seq (10x) cells colored by sample identity, demonstrating effective integration across sequencing technologies and experimental conditions (mock, Asian 4d, Asian 6d African 4d, African 6d). Cells from both platforms are well-mixed within shared cell-type clusters, indicating successful integration without technology-driven batch effects. **(B)** Proportion of cells derived from spatial transcriptomics versus scRNA-seq within each identified cluster. Balanced contributions from both modalities across clusters confirm minimal technology-driven batch effects and validate the integration approach. **(C)** Schematic illustration of local cell density computation methodology used throughout the study. For each cell serving as an anchor point, all neighboring cells within a 0.08 mm² circular spatial neighborhood are counted to calculate local cell density. This metric quantifies the local abundance of specific cell types (e.g., microglia, infected cells) within tissue microenvironments. Cell types shown are for illustration purposes only.

**Figure 1S4.**
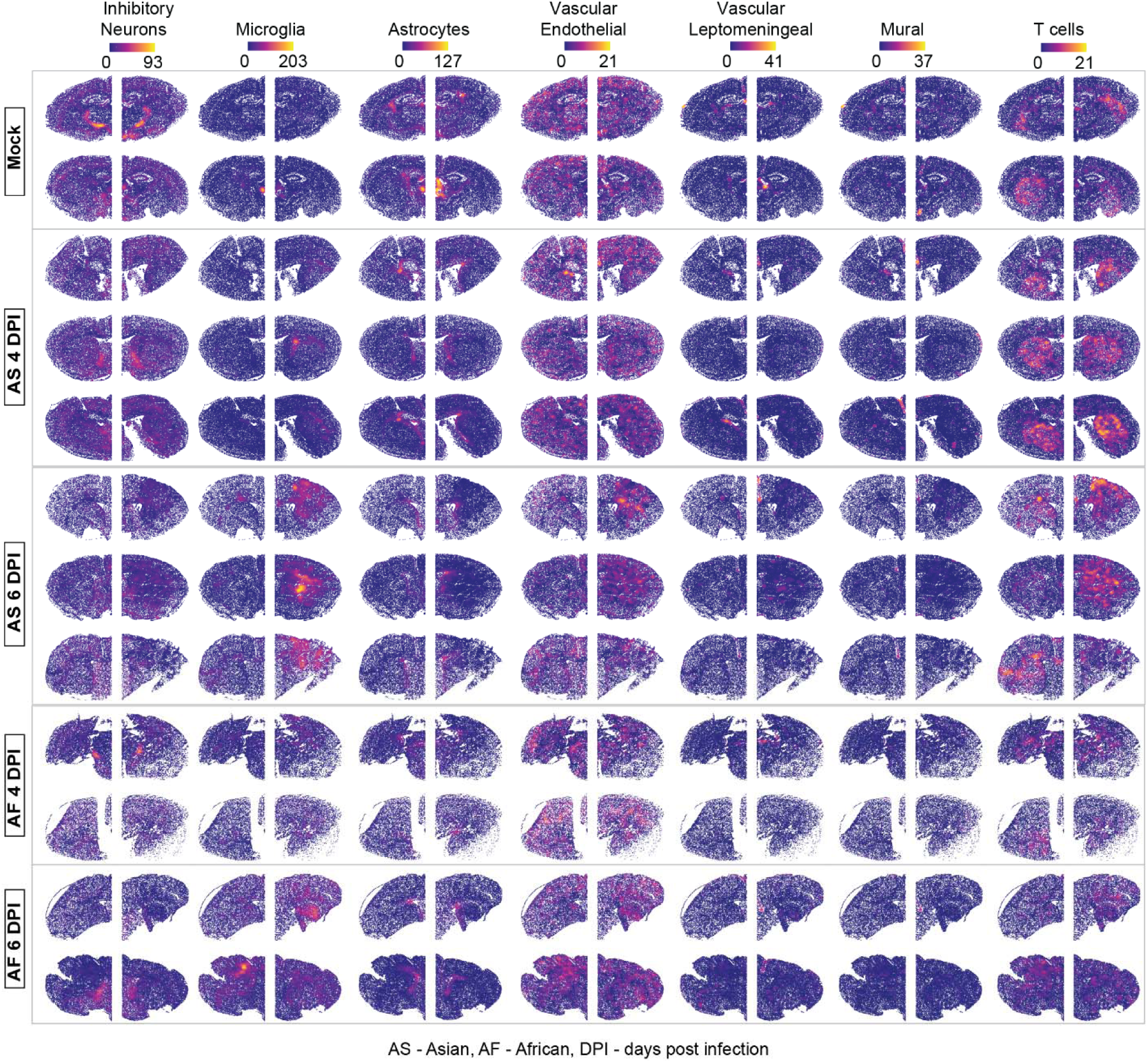
Spatial organization of major cell types across infection strains and time points. Spatial tissue maps showing the spatial distribution and local density of major cell types across all experimental conditions. Columns correspond to individual cell types (inhibitory neurons, microglia, astrocytes, vascular endothelial cells, vascular leptomeningeal cells, mural cells, and T cells); rows represent conditions (mock, Asian 4 dpi, Asian 6 dpi, African 4 dpi, African 6 dpi). Cells are colored by local cell-type density, calculated by counting neighboring cells of the same type within a 0.08 mm² spatial neighborhood centered on each cell, revealing strain- and time-dependent remodeling of cellular organization during ZIKV infection. Note: Excitatory neurons and oligodendrocytes are shown separately in main Figure 1G-H.

**Figure 2S1.**
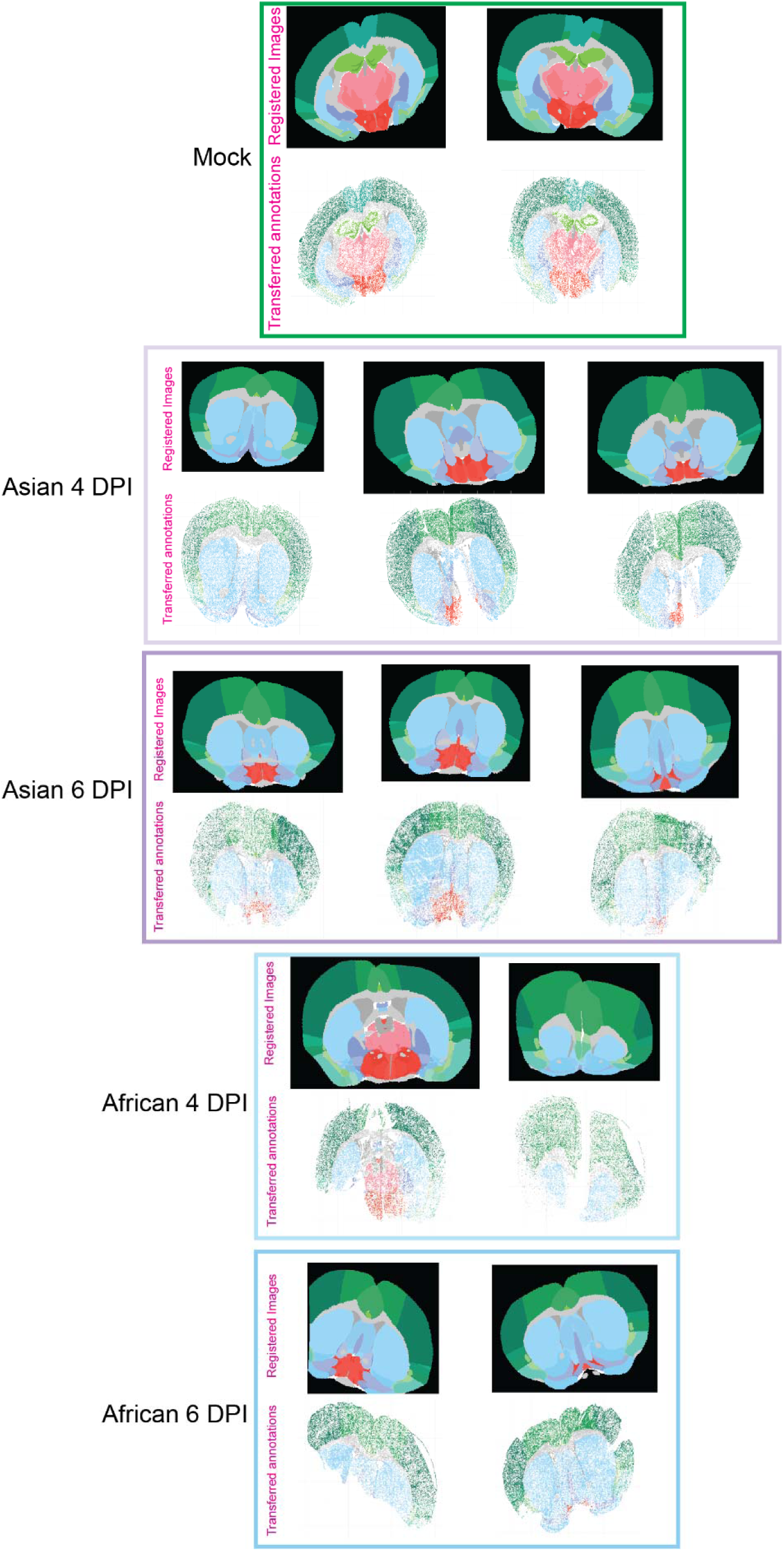
Registration of spatial transcriptomic data to the Allen Common Coordinate Framework and transfer of anatomical annotations. Spatial tissue maps showing all brain samples (mock, Asian 4 dpi, Asian 6 dpi, African 4 dpi, African 6 dpi) aligned to the Allen Common Coordinate Framework version 3 (CCFv3) reference atlas using QuickNII. Each panel shows a representative brain section with cells positioned according to their registered coordinates.

**Figure 2S2.**
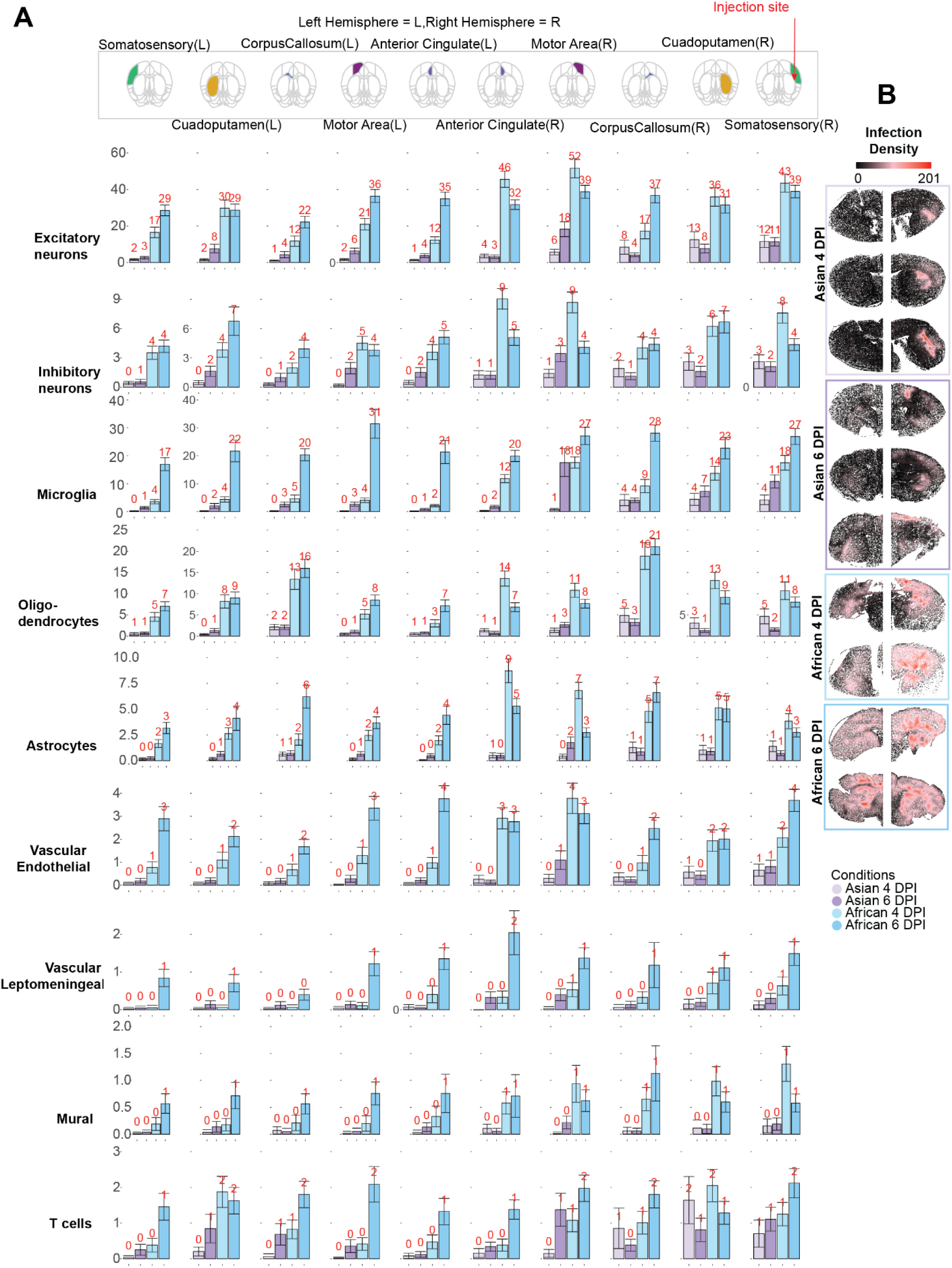
Cell-type-specific infection densities across brain structures. (A) Bar plots showing densities of infected cells stratified by cell type across major brain structures under four experimental conditions (Asian 4 dpi, Asian 6 dpi, African 4 dpi, African 6 dpi). Local infected cell-type densities were calculated by counting neighboring infected cells of the same type within a 0.08 mm² spatial neighborhood centered on each cell. Bars represent mean infected cell-type density across individual cells within each structure; error bars indicate standard deviation across cells scaled by 6 (SD/6) to visualize relative dispersion. Brain structures are organized by hemisphere (R: right; L: left). (B) Spatial tissue maps showing the spatial distribution of infected cells across all conditions in brains. Cells are positioned according to their spatial coordinates and colored by local infection density (number of infected neighboring cells within a 0.08 mm² neighborhood). These maps complement the quantitative bar plots in panel A by visualizing the spatial organization of infection across brain regions.

**Figure 2S3.**
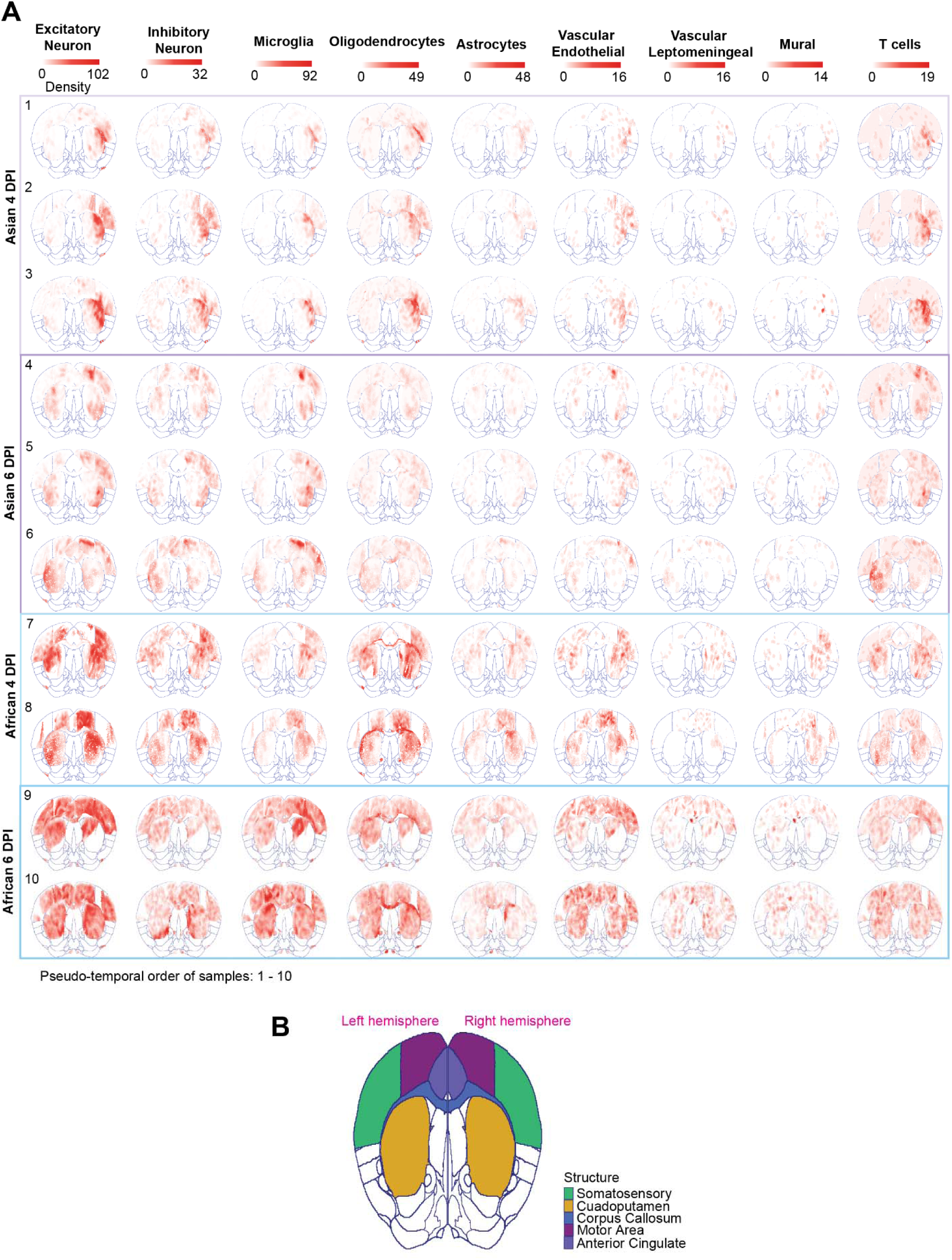
Spatial distribution of infected cell types ordered by pseudo-temporal progression. **(A)** Spatial tissue maps showing cells from major brain structures projected onto a shared reference space and ordered from top to bottom (rows 1-10) according to pseudo-temporal rank, defined by sample infection rate (percentage of infected cells per sample). Each column represents a different infected cell type (excitatory neurons, inhibitory neurons, microglia, astrocytes, oligodendrocytes). Cells are colored by local infected cell-type density, calculated by counting neighboring infected cells of the same type within a 0.08 mm² neighborhood centered on each cell. This visualization complements main Figure 2A by revealing cell-type-specific spatial patterns of infection progression across the pseudo-temporal trajectory. **(B)** Allen Common Coordinate Framework (CCFv3) reference map colored by corresponding major brain structures, providing anatomical context for interpreting the spatial maps in panel A. Structures are labeled and color-coded to facilitate identification of regions showing cell-type-specific infection dynamics.

**Figure 2S4.**
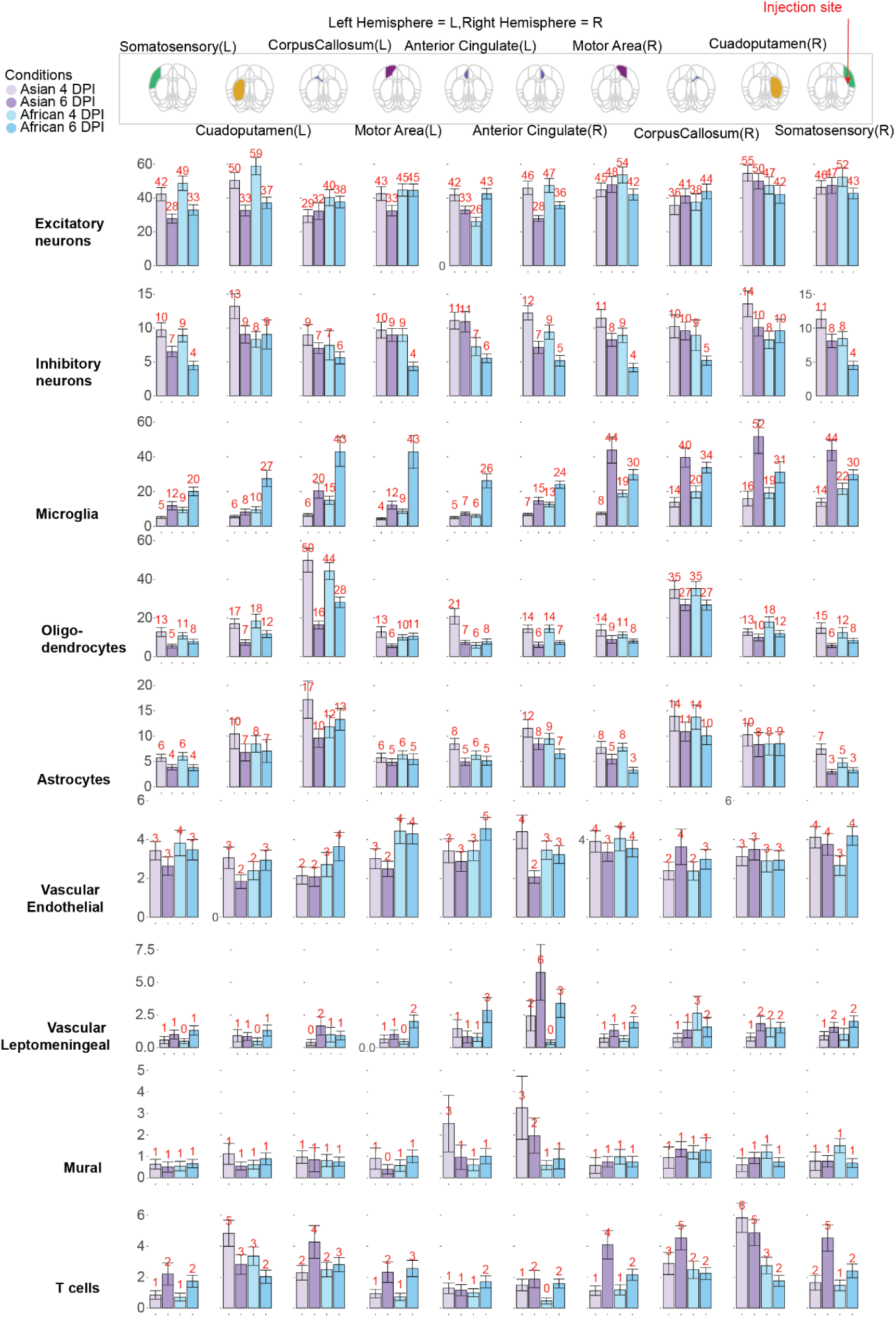
Cell-type density profiles across major brain structures. Bar plots showing local densities of major cell types (excitatory neurons, inhibitory neurons, microglia, astrocytes, oligodendrocytes, vascular cells) within each brain structure across experimental conditions (Asian 4 dpi, Asian 6 dpi, African 4 dpi, African 6 dpi). Cell-type densities were calculated by counting neighboring cells of the same type within a 0.08 mm² spatial neighborhood centered on each cell. Bars represent mean cell-type density across individual cells within each structure; error bars indicate standard deviation across cells scaled by 6 (SD/6) to visualize relative dispersion.

**Figure 2S5.**
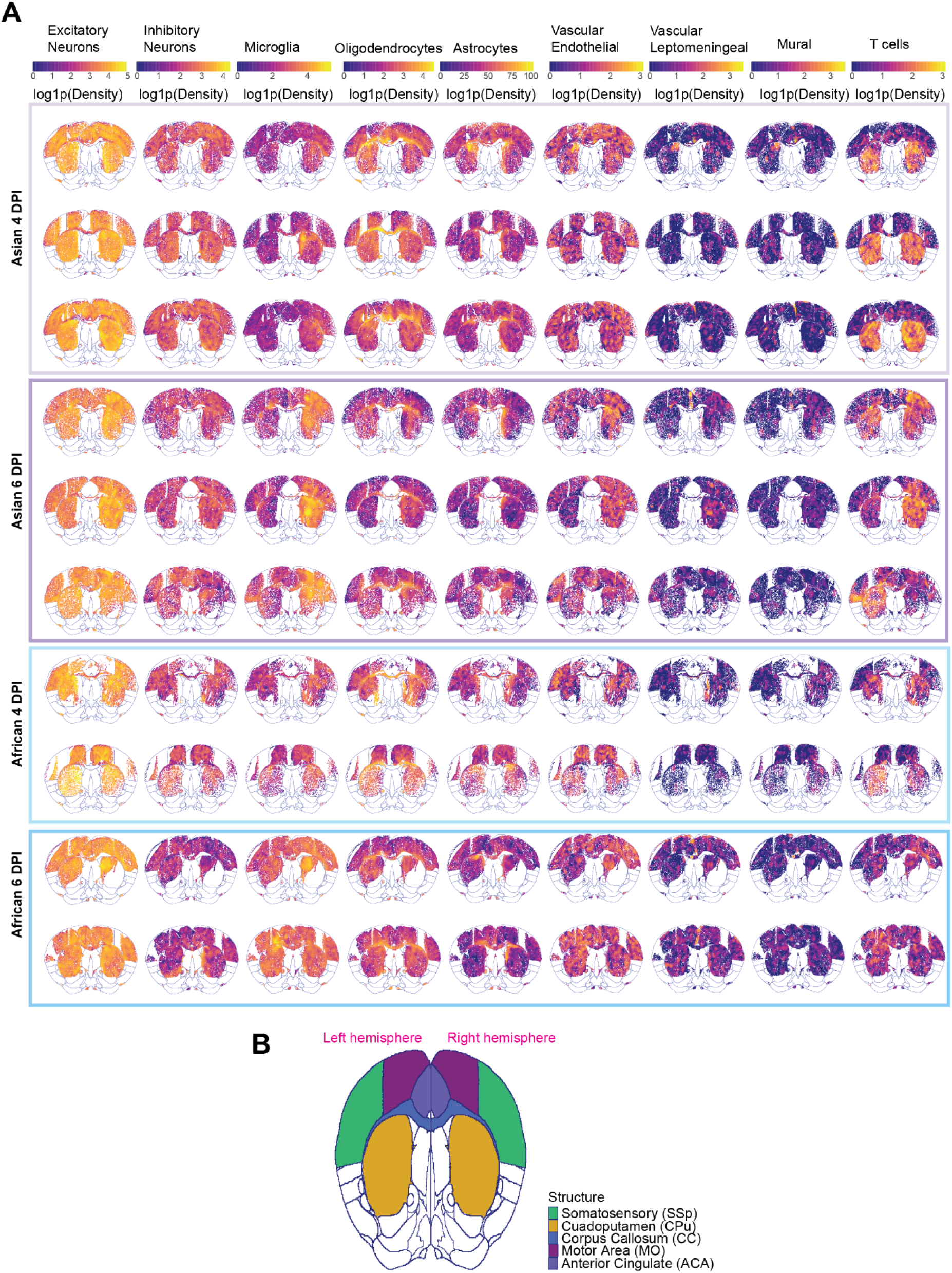
Spatial organization of cell-type densities within major brain structures across infection conditions. (A) Spatial tissue maps showing local cell-type densities within major brain structures—including somatosensory cortex (SSp), caudate-putamen (CPu), motor area (MO), corpus callosum (CC), and anterior cingulate area (ACA) across infected conditions (Asian 4 dpi, Asian 6 dpi, African 4 dpi, African 6 dpi). Rows represent different cell types (excitatory neurons, inhibitory neurons, microglia, astrocytes, oligodendrocytes); columns represent different brain structures. Local cell-type densities were calculated by counting neighboring cells of the same type within a 0.08 mm² neighborhood centered on each cell, then natural log-transformed as log1p(density) for visualization to accommodate the range of density values. (B) Allen Common Coordinate Framework (CCFv3) reference map colored by corresponding brain structures, providing anatomical context for the spatial tissue maps in panel A. Structure abbreviations and color coding match panel A to facilitate spatial localization.

**Figure 2S6.**
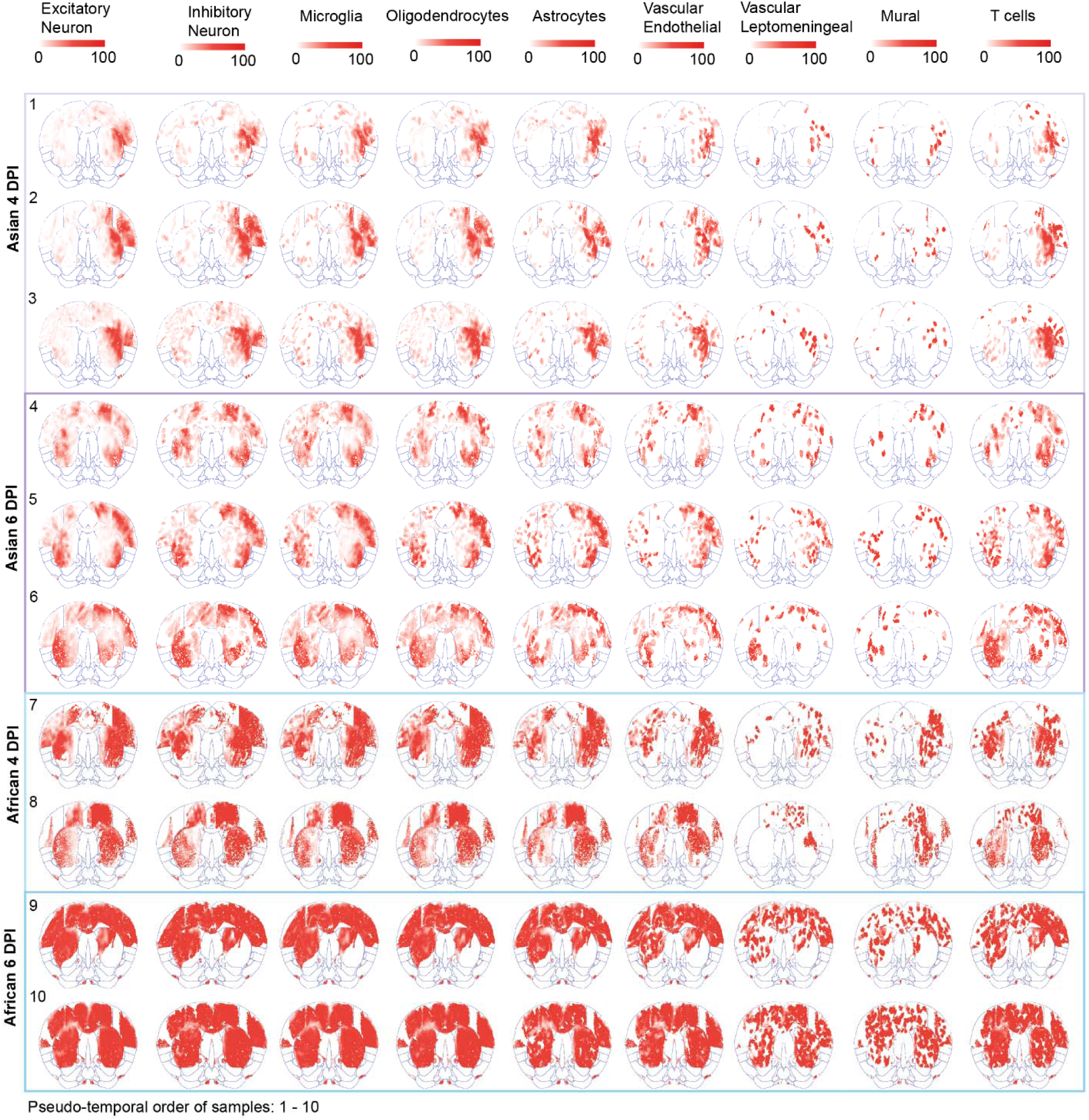
Spatial organization of normalized infected cell-type densities within major brain structures. Spatial tissue maps showing normalized infected cell-type densities within major brain structures, across infected conditions (Asian 4 dpi, Asian 6 dpi, African 4 dpi, African 6 dpi). Cells are colored by normalized infected cell-type density, calculated by accounting for both local infection burden (total infected cells) and local cell-type abundance (total cells of that type) within 0.08 mm² neighborhoods. This figure spatially resolves the cell-type-specific infection preferences quantified in Figure 2F, revealing how preferentially targeted versus avoided cell types are distributed across anatomical regions.

**Figure 3S1.**
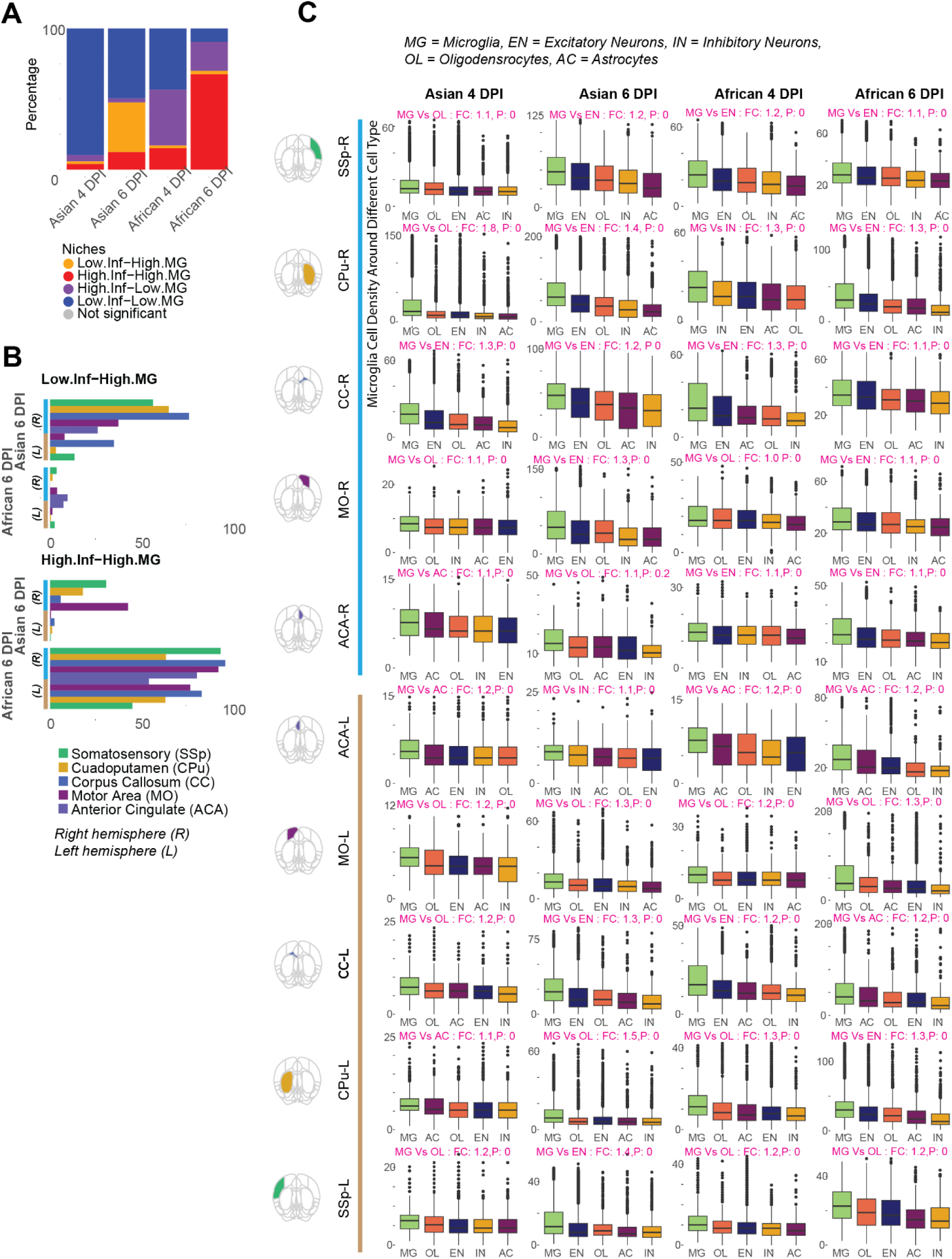
Spatial organization and cellular context of microglial responses across infection conditions. **(A)** Stacked bar plots showing the proportional distribution of bivariate Moran’s I-defined spatial niches across infection conditions (Asian 4 dpi, Asian 6 dpi, African 4 dpi, African 6 dpi) and brain structures. Four spatial niche categories are shown: High infection-High microglia (red), Low infection-High microglia (yellow; infection-contained), High infection-Low microglia (purple), and Low infection-Low microglia (gray). Each bar represents the proportion of cells within each structure classified into each niche category. This reveals how the spatial relationship between infection and microglial responses varies across conditions and anatomical regions. **(B)** Distribution of immune-enriched spatial niches across brain structures at 6 dpi. Bar plots show the percentage of Low infection-High microglia (infection-contained) and High infection-High microglia niches within individual brain structures for Asian and African strain-infected samples, highlighting strain-specific differences in spatial infection control. Structures are organized by hemisphere (R: right; L: left). **(C)** Local microglial density in the vicinity of different cell types. Box plots show microglial density within 0.08 mm² tissue microenvironments centered on cells of different types (microglia, excitatory neurons, inhibitory neurons, astrocytes, oligodendrocytes), pooled across all infected conditions. Cell types are ordered by descending median microglial density, reflecting preferential microglial spatial associations. Boxes indicate interquartile range (25th-75th percentile); whiskers extend to 1.5× IQR. Fold change (FC) comparing microglial density around the top two cell types where microglia most aggregated and P-value from two-sided t-test are shown.

**Figure 3S2.**
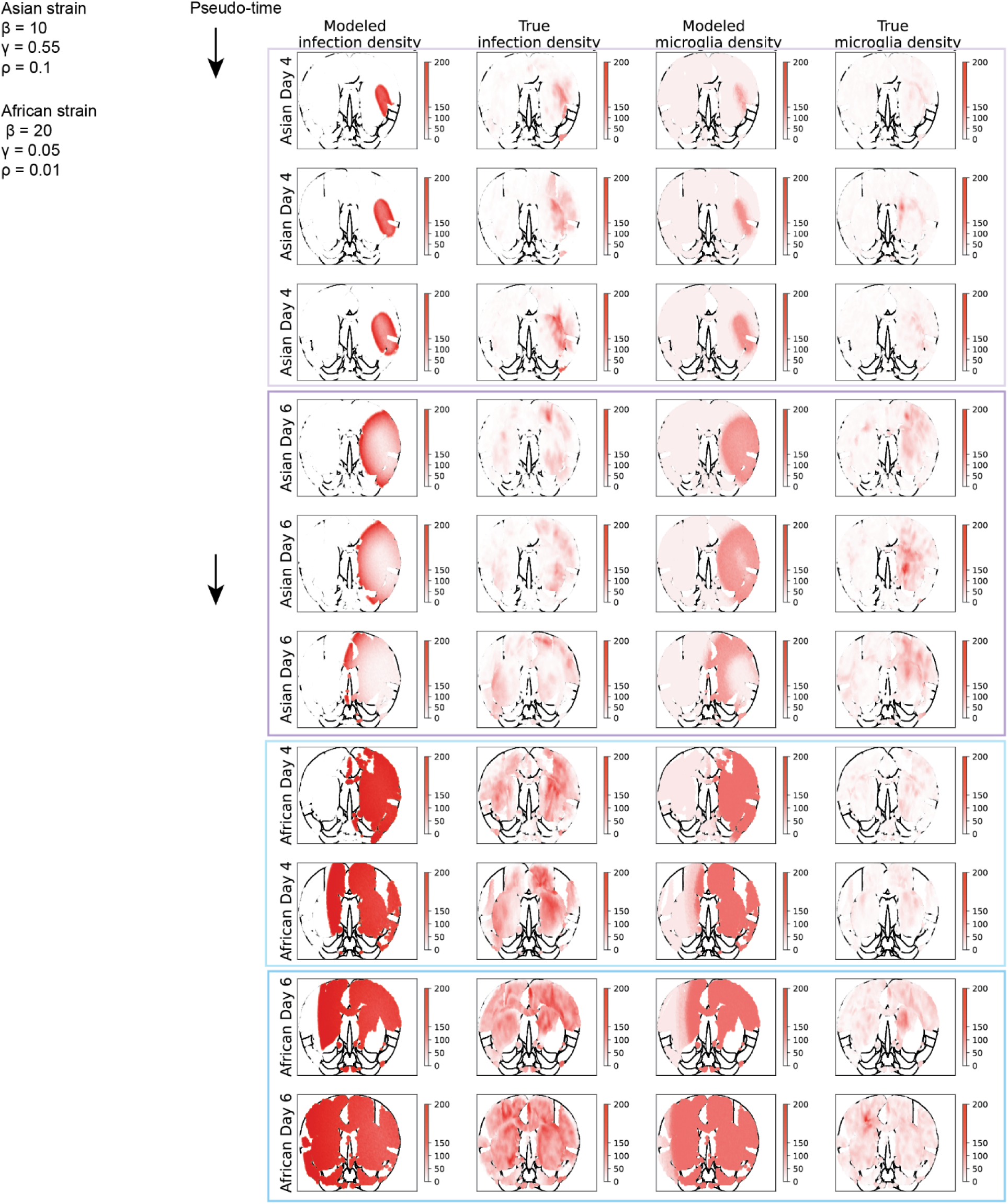
Model validation: simulated versus observed infection and microglial dynamics. Spatial maps comparing model-simulated and empirically observed infection and microglial densities for representative brain samples from Asian and African ZIKV strains at 4 and 6 dpi. Rows represent individual samples from each strain-timepoint combination; columns show: (1) modeled infection density, (2) observed infection density, (3) modeled microglial density, and (4) observed microglial density. Infection and microglial dynamics were simulated using a stochastic, density-based framework inspired by the Reed-Frost chain-binomial model (see Methods), governed by three parameters: β (infection expansion potential), γ (microglial effectiveness in dampening infection spread), and ρ (infected cell clearance probability). Strain-specific parameter regimes were fitted to the observed data: Asian strain (β = 10, γ = 0.55, ρ = 0.1) exhibits controlled infection expansion with effective microglial containment, whereas African strain (β = 20, γ = 0.05, ρ = 0.01) shows high viral expansion with reduced immune effectiveness. Close correspondence between modeled and observed spatial patterns validates the model’s ability to capture strain-specific infection-immune dynamics and supports its use for exploring parameter space beyond experimentally measured timepoints (Figure 3G).

**Figure 5S1.**
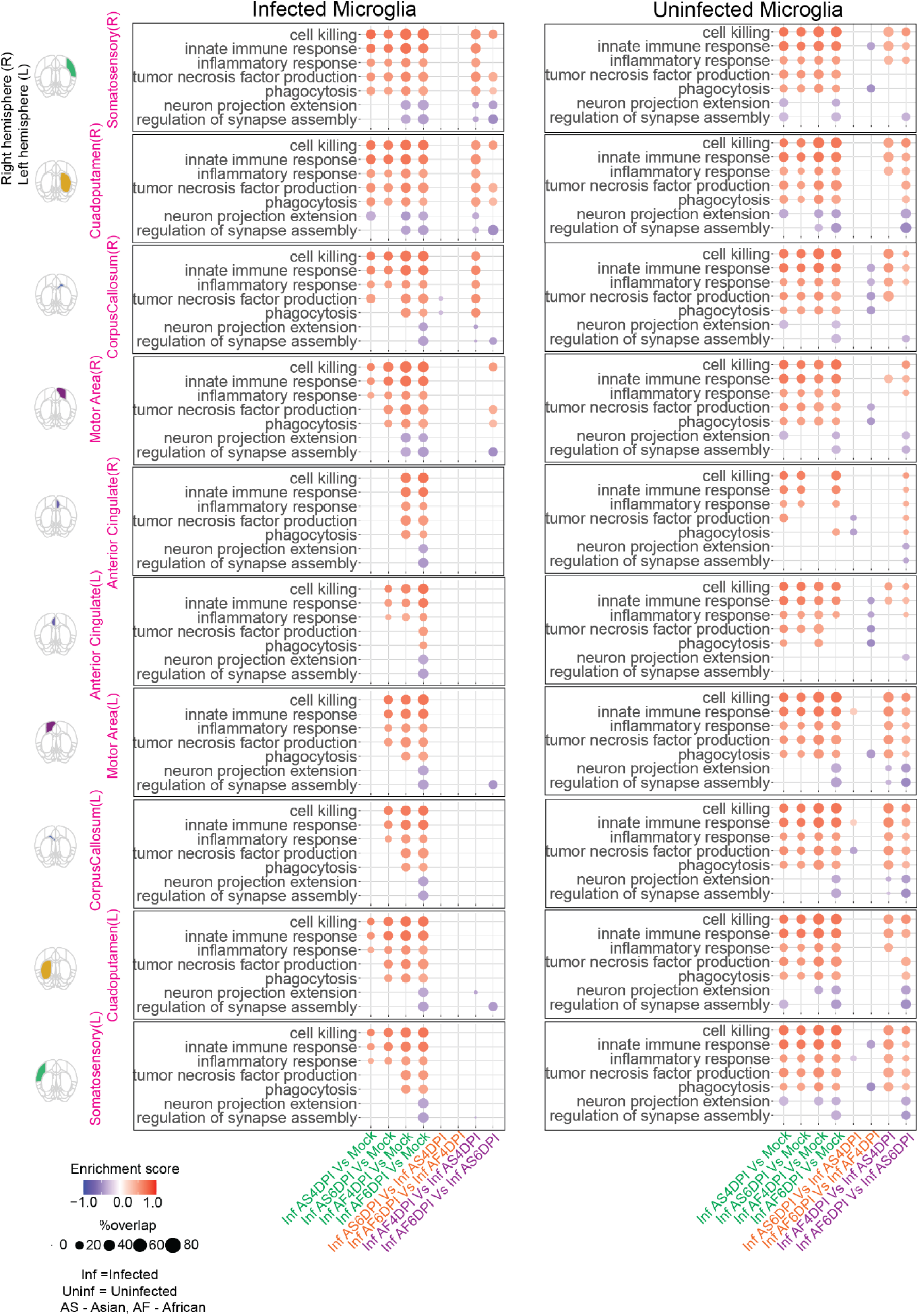
Regional functional reprogramming of infected and uninfected microglia across brain structures. Bubble plots showing enriched Gene Ontology biological process (GO:BP) terms (GSEA, adjusted P-value < 0.05) for microglia stratified by infection status and brain structure. Left panels: Enrichment in infected microglia from Asian and African ZIKV-infected samples at 4 and 6 dpi. Comparisons include infected microglia versus mock controls and direct strain-to-strain contrasts (African vs Asian) at matched time points. Right panels: Corresponding enrichment in uninfected microglia from the same infected brains, analyzed using identical comparison schemes. Bubble color indicates enrichment direction (red: upregulated; blue: downregulated); bubble size reflects the proportion of genes within each gene set that are significantly differentially expressed (|log₂ fold change| ≥ 0.30, adjusted P-value < 0.05). Results demonstrate both cell-intrinsic functional reprogramming in infected microglia and tissue-wide bystander activation in uninfected microglia within infected brains, with regional consistency in strain-dependent differences in innate immune and inflammatory pathway activation (as noted in main text Section 5). The observation that uninfected microglia exhibit similar functional changes as infected microglia suggests viral infection induces broader tissue-level immune state transitions beyond directly infected cells.

**Figure 5S2.**
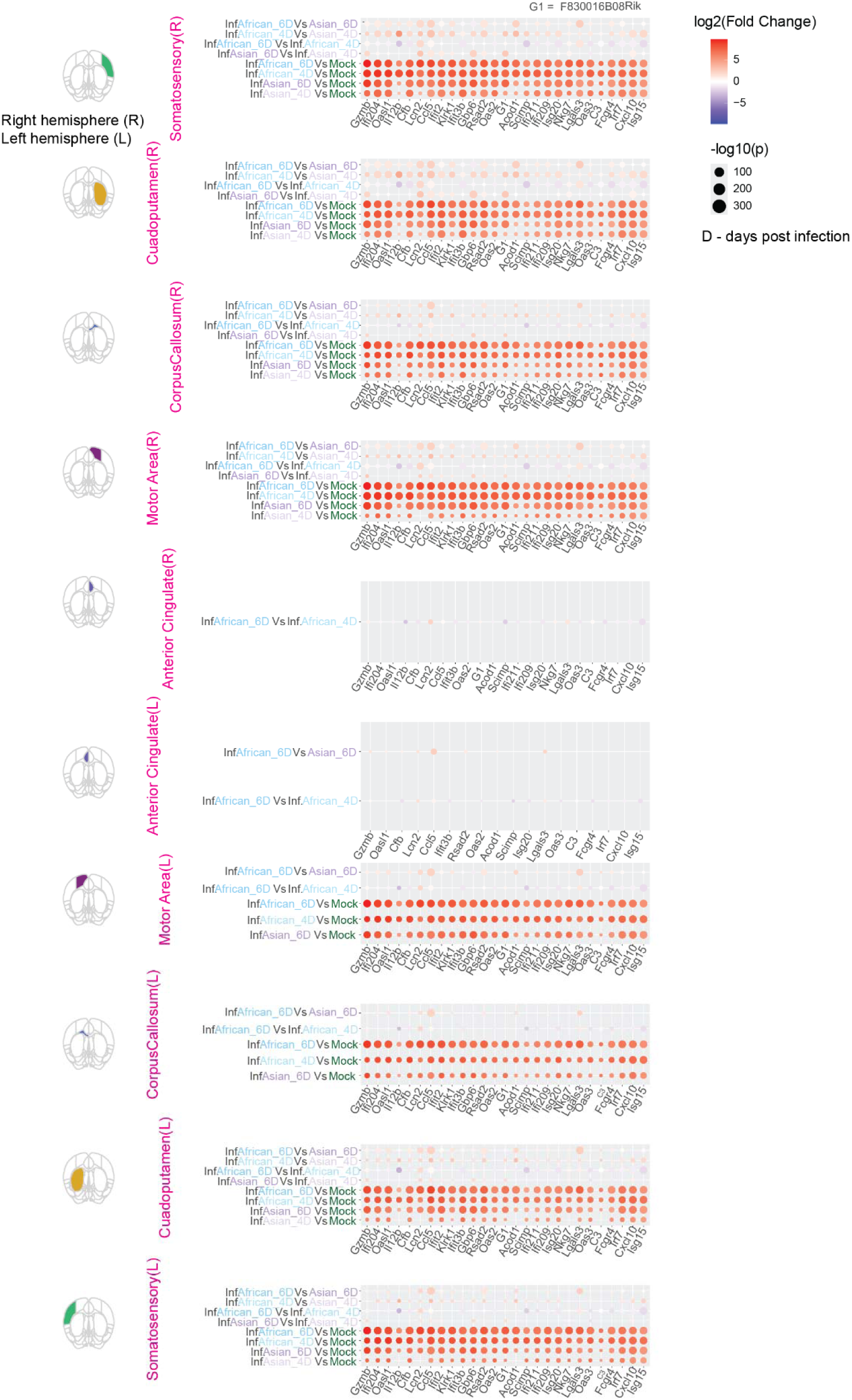
Innate immune response gene expression in infected microglia across brain structures. Bubble plots showing gene-level log₂ fold changes for innate immune response-associated genes in infected microglia across major brain structures. Rows correspond to pairwise comparisons (infected Asian 4 dpi vs mock, infected Asian 6 dpi vs mock, infected African 4 dpi vs mock, infected African 6 dpi vs mock, and strain-matched comparisons at 4 dpi and 6 dpi); columns represent individual genes within the innate immune response pathway. Bubble color denotes log₂ fold change (red: upregulated; blue: downregulated); bubble size reflects statistical significance (–log₁₀ adjusted P-value). These data reveal conserved activation of innate immune programs across strains with regional consistency.

**Figure 5S3.**
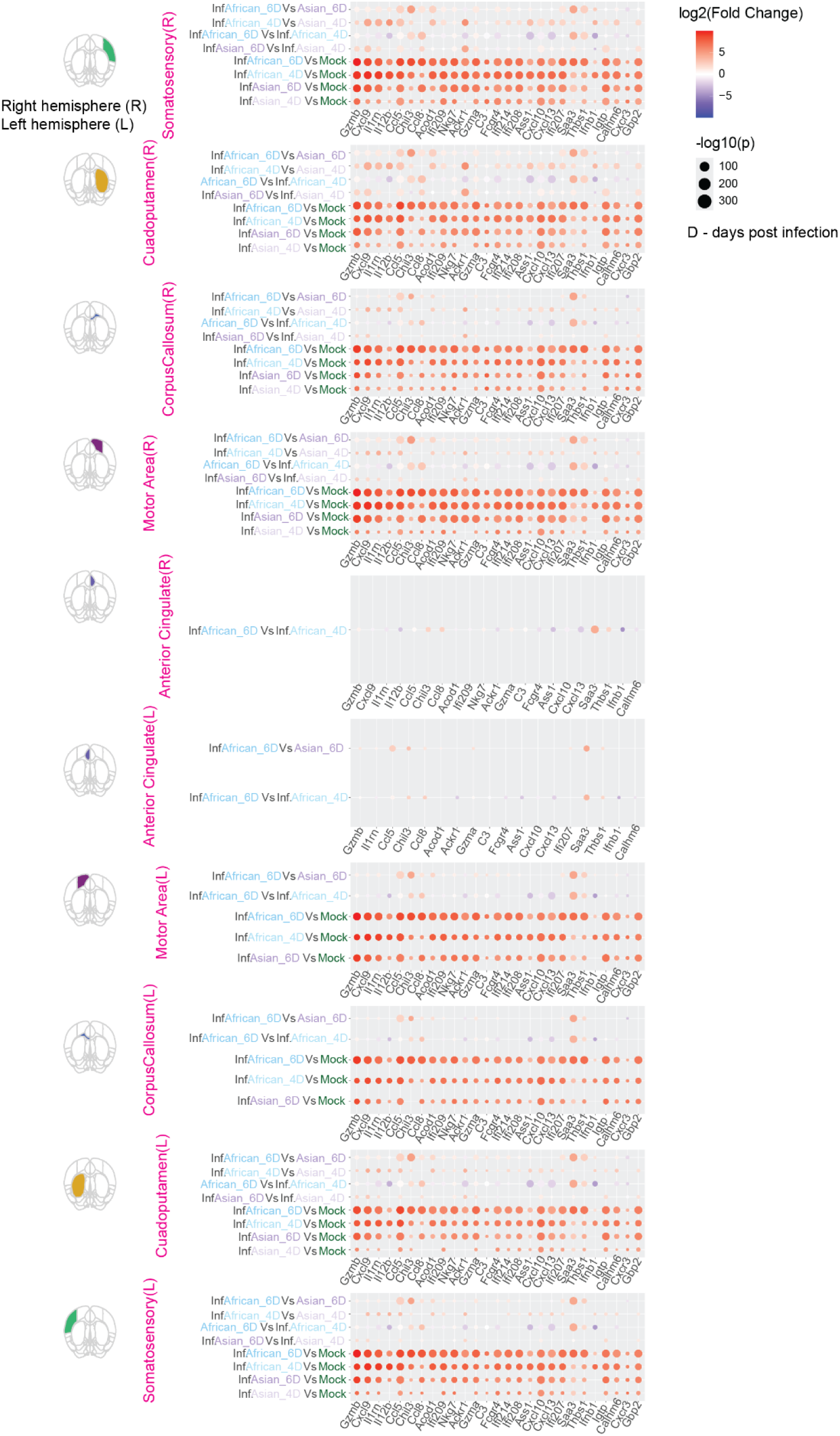
Inflammatory response gene expression in infected microglia across brain structures. Bubble plots showing gene-level log₂ fold changes for inflammatory response-associated genes in infected microglia across major brain structures. Rows correspond to pairwise comparisons (infected Asian 4 dpi vs mock, infected Asian 6 dpi vs mock, infected African 4 dpi vs mock, infected African 6 dpi vs mock, and strain-matched comparisons); columns represent individual genes within the inflammatory response pathway. Bubble color denotes log₂ fold change (red: upregulated; blue: downregulated); bubble size reflects statistical significance (–log₁₀ adjusted P-value).

**Figure 5S4.**
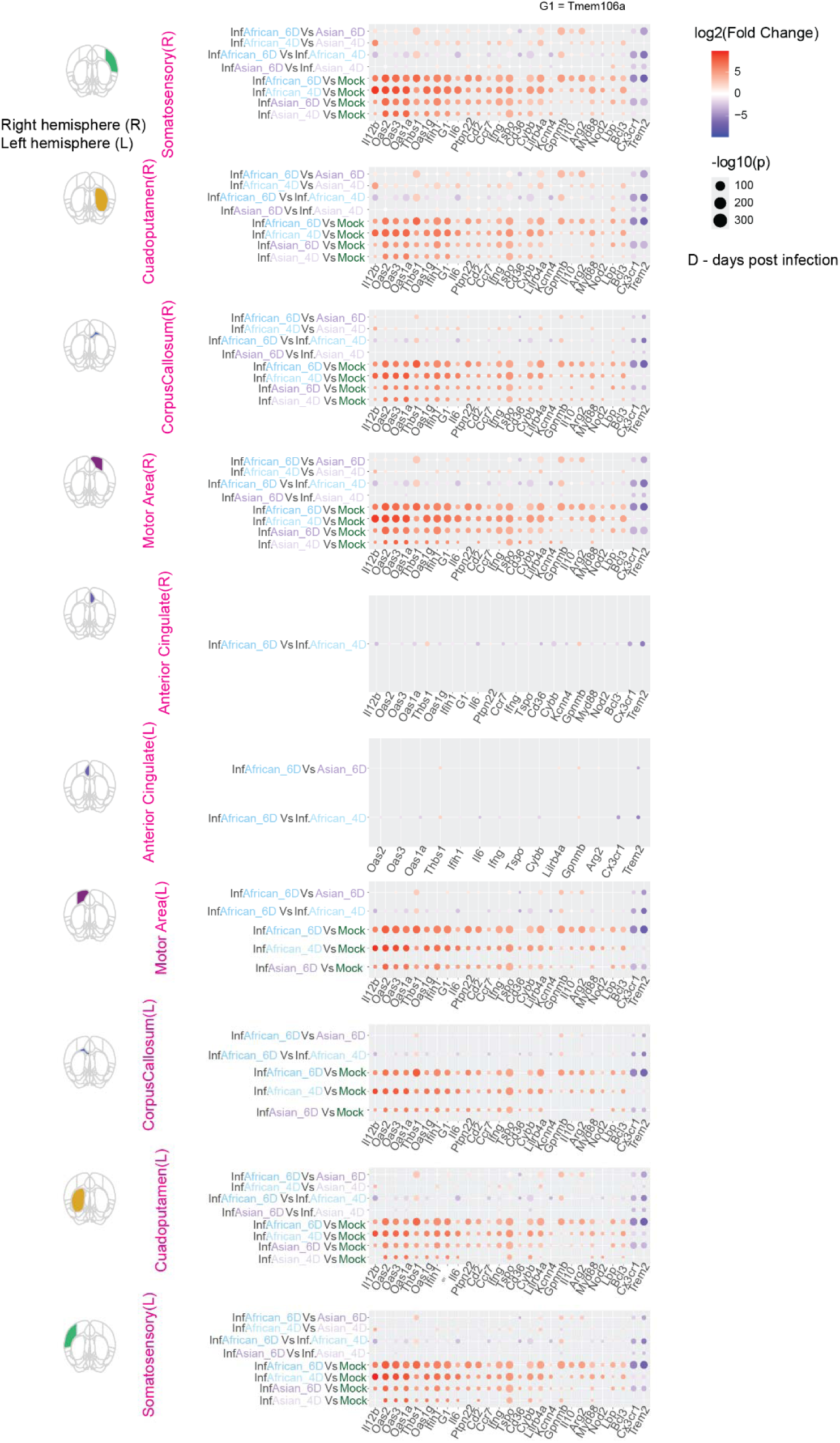
TNF production gene expression in infected microglia across brain structures. Bubble plots showing gene-level log₂ fold changes for TNF production-associated genes in infected microglia across major brain structures. Rows correspond to pairwise comparisons (infected Asian 4 dpi vs mock, infected Asian 6 dpi vs mock, infected African 4 dpi vs mock, infected African 6 dpi vs mock, and strain-matched comparisons); columns represent individual genes within the TNF production pathway. Bubble color denotes log₂ fold change (red: upregulated; blue: downregulated); bubble size reflects statistical significance (–log₁₀ adjusted P-value).

**Figure 5S5.**
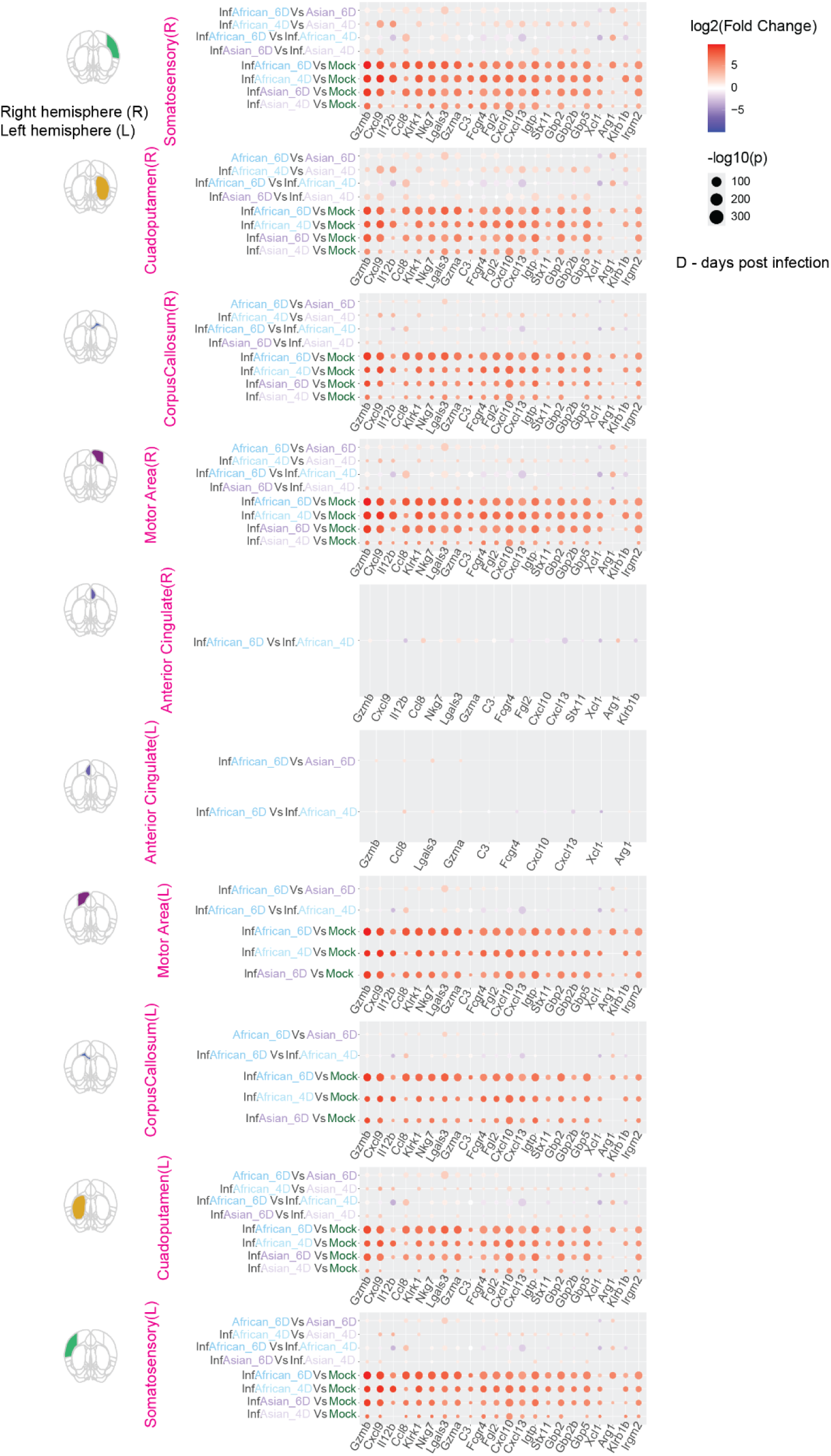
Cell killing gene expression in infected microglia across brain structures. Bubble plots showing gene-level log₂ fold changes for cell killing-associated genes in infected microglia across major brain structures. Rows correspond to pairwise comparisons (infected Asian 4 dpi vs mock, infected Asian 6 dpi vs mock, infected African 4 dpi vs mock, infected African 6 dpi vs mock, and strain-matched comparisons); columns represent individual genes within the cell killing pathway. Bubble color denotes log₂ fold change (red: upregulated; blue: downregulated); bubble size reflects statistical significance (–log₁₀ adjusted P-value).

**Figure 5S6.**
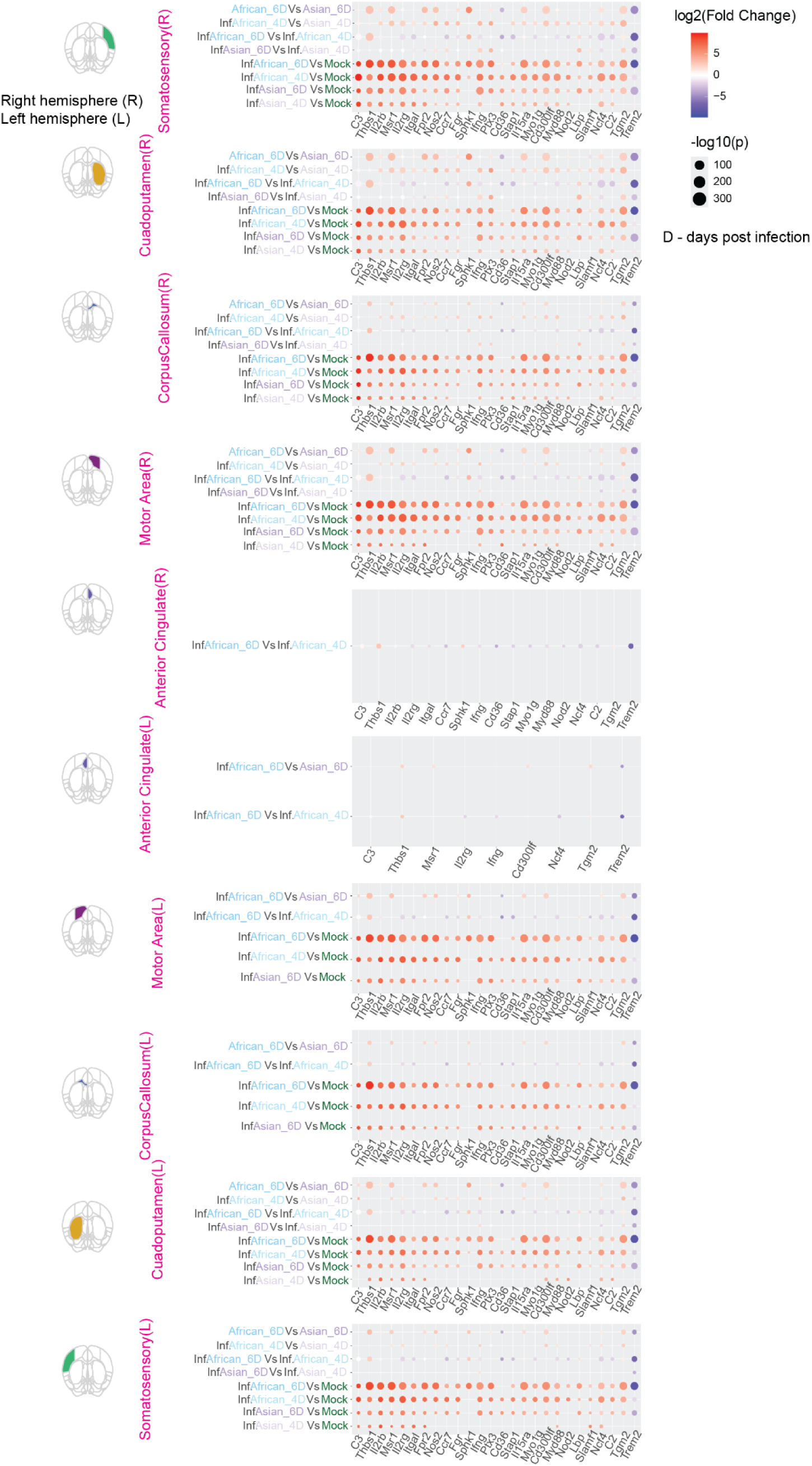
Phagocytosis gene expression in infected microglia across brain structures. Bubble plots showing gene-level log₂ fold changes for phagocytosis-associated genes in infected microglia across major brain structures. Rows correspond to pairwise comparisons (infected Asian 4 dpi vs mock, infected Asian 6 dpi vs mock, infected African 4 dpi vs mock, infected African 6 dpi vs mock, and strain-matched comparisons); columns represent individual genes within the phagocytosis pathway. Bubble color denotes log₂ fold change (red: upregulated; blue: downregulated); bubble size reflects statistical significance (–log₁₀ adjusted P-value).

**Figure 5S7.**
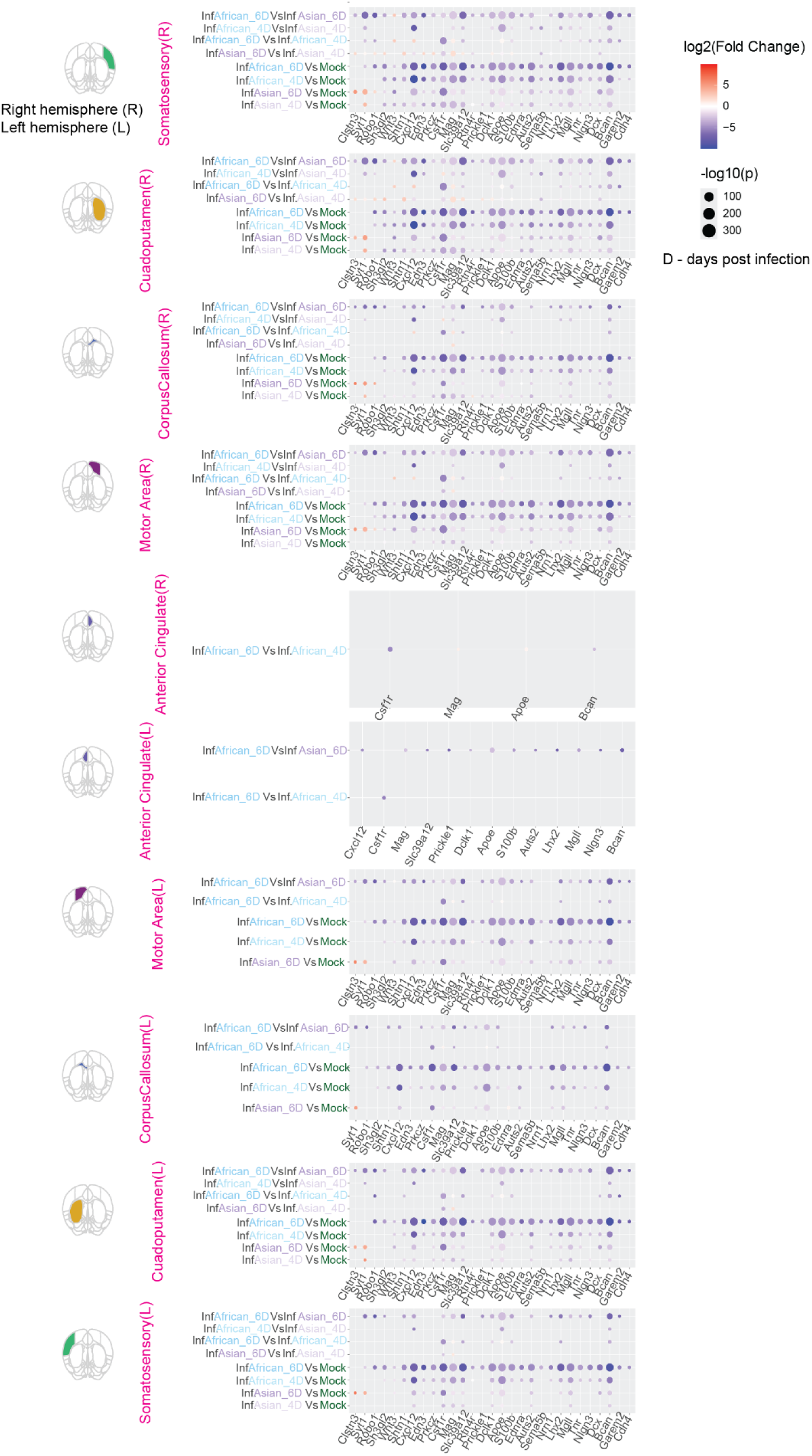
Neuron projection extension gene expression in infected microglia across brain structures. Bubble plots showing gene-level log₂ fold changes for neuron projection extension-associated genes in infected microglia across major brain structures. Rows correspond to pairwise comparisons (infected Asian 4 dpi vs mock, infected Asian 6 dpi vs mock, infected African 4 dpi vs mock, infected African 6 dpi vs mock, and strain-matched comparisons); columns represent individual genes within the neuron projection extension pathway. Bubble color denotes log₂ fold change (red: upregulated; blue: downregulated); bubble size reflects statistical significance (–log₁₀ adjusted P-value).

**Figure 5S8.**
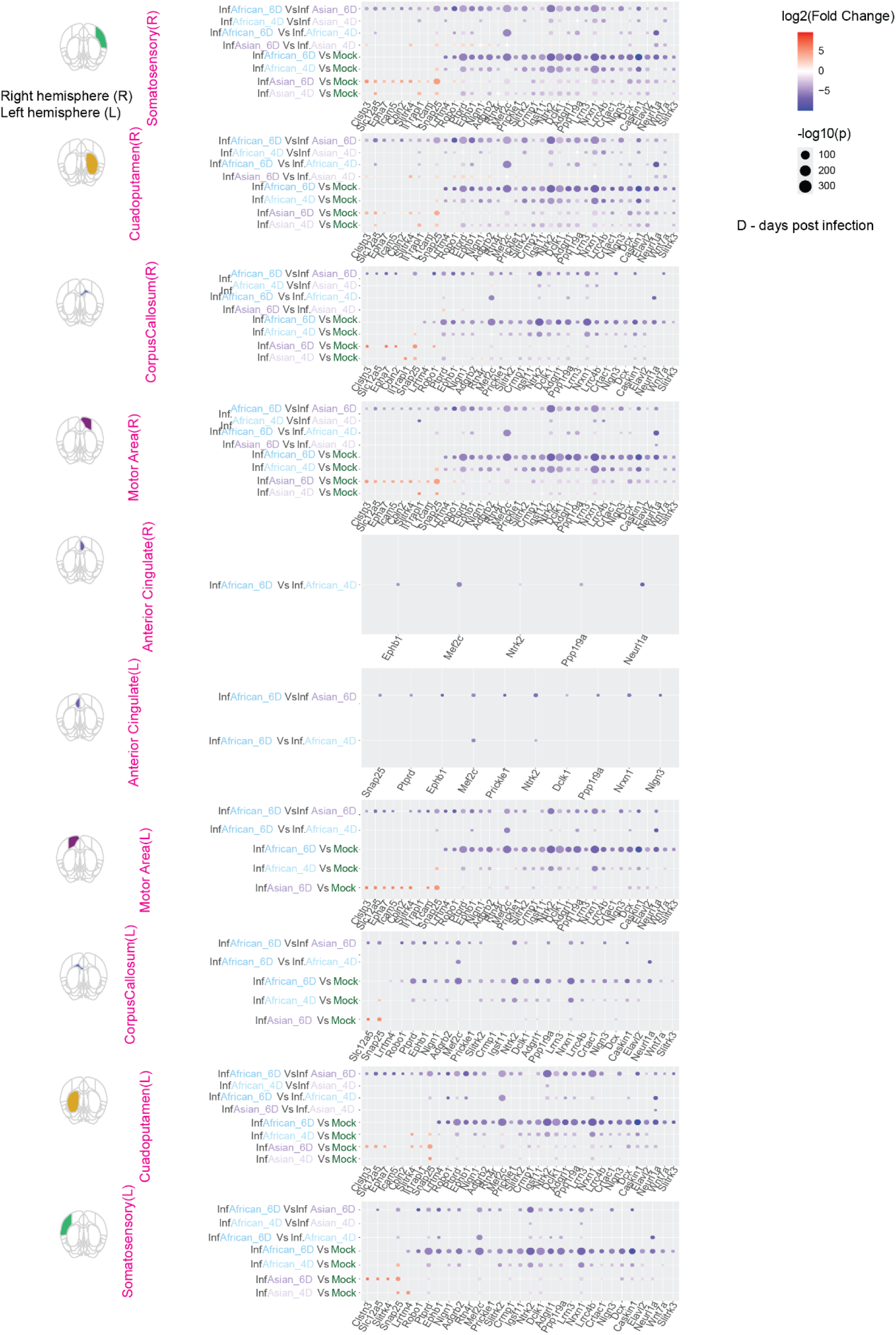
Regulation of synapse assembly gene expression in infected microglia across brain structures. Bubble plots showing gene-level log₂ fold changes for synapse assembly regulation-associated genes in infected microglia across major brain structures. Rows correspond to pairwise comparisons (infected Asian 4 dpi vs mock, infected Asian 6 dpi vs mock, infected African 4 dpi vs mock, infected African 6 dpi vs mock, and strain-matched comparisons); columns represent individual genes within the synapse assembly regulation pathway. Bubble color denotes log₂ fold change (red: upregulated; blue: downregulated); bubble size reflects statistical significance (-log₁₀ adjusted P-value).

**Figure 5S9.**
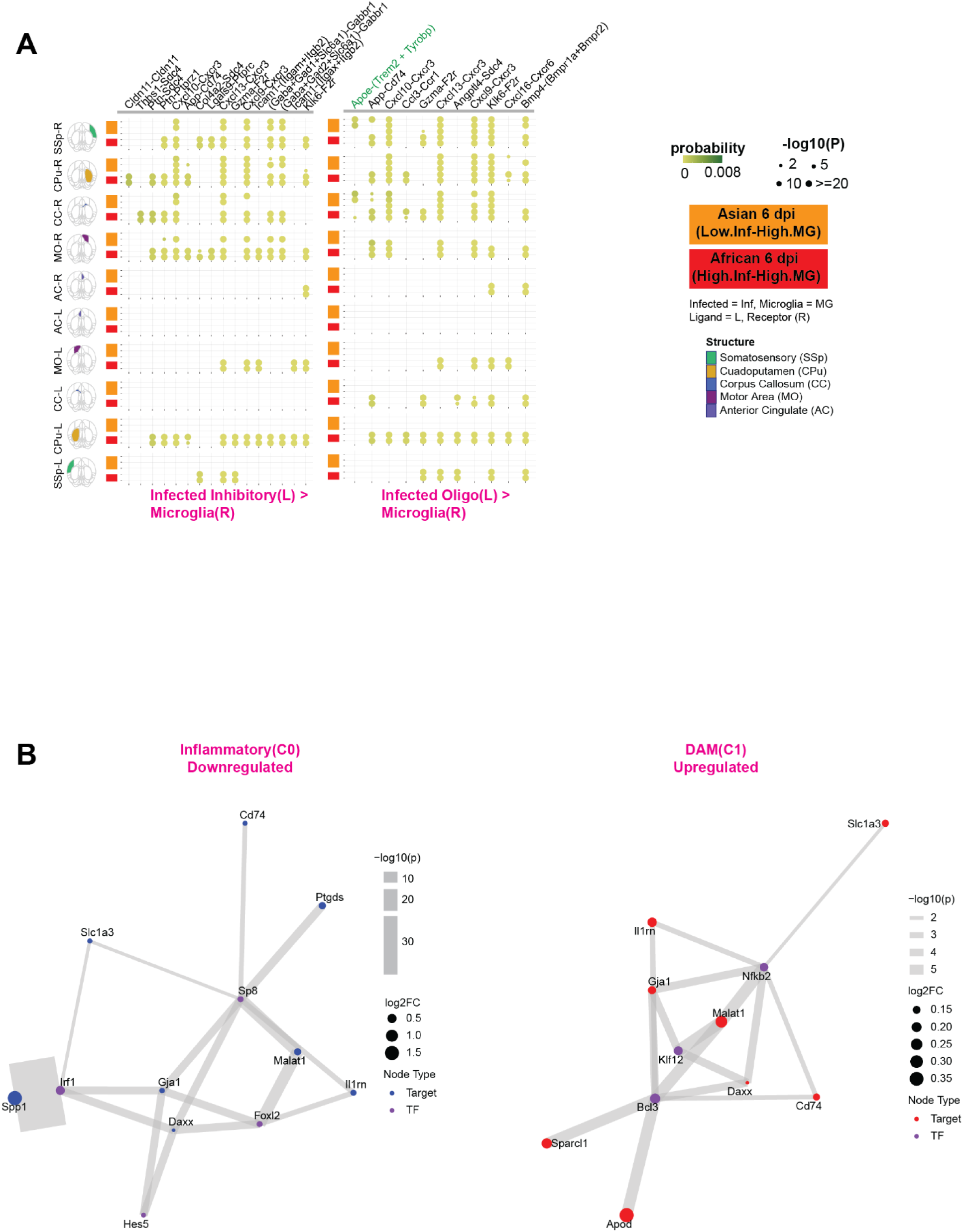
Extended cell-cell communication networks and transcription factor regulatory networks. (A) Additional cell-cell communication networks between infected cell types and microglia in spatially distinct niches. CellChat-inferred ligand-receptor interactions for infected inhibitory neurons (left panel) and infected oligodendrocytes (right panel) as ligand sources communicating with microglia as receptors. Analysis restricted to infection-contained regions (Low infection-High microglia) in Asian 6 dpi samples (yellow) versus highly infected regions (High infection-High microglia) in African 6 dpi samples (red). Bubble color represents communication probability; bubble size denotes interaction significance (–log₁₀ P-value). These panels complement main Figure 5D by revealing cell-type-specific signaling from infected inhibitory neurons and oligodendrocytes to surrounding microglia. (B) Extended gene regulatory networks for microglial subtypes. Network plots showing predicted regulatory relationships between top transcription factors (TFs; purple nodes) and their significant target genes (adjusted P-value < 0.05) that are differentially expressed compared to Homeostatic microglia baseline. Left panel: Inflammatory microglia showing TF-driven downregulation of targets. Right panel: DAM microglia showing TF-driven upregulation of targets. Target gene nodes are colored red (upregulated) or blue (downregulated) relative to Homeostatic state; node size represents absolute log₂ fold change. Edge width reflects strength of TF-gene regulatory prediction from CellOracle (-log₁₀ P-value). These panels complement main Figure 5F by showing the reciprocal regulatory patterns (Inflammatory downregulated genes and DAM upregulated genes).

**Figure 5S10.**
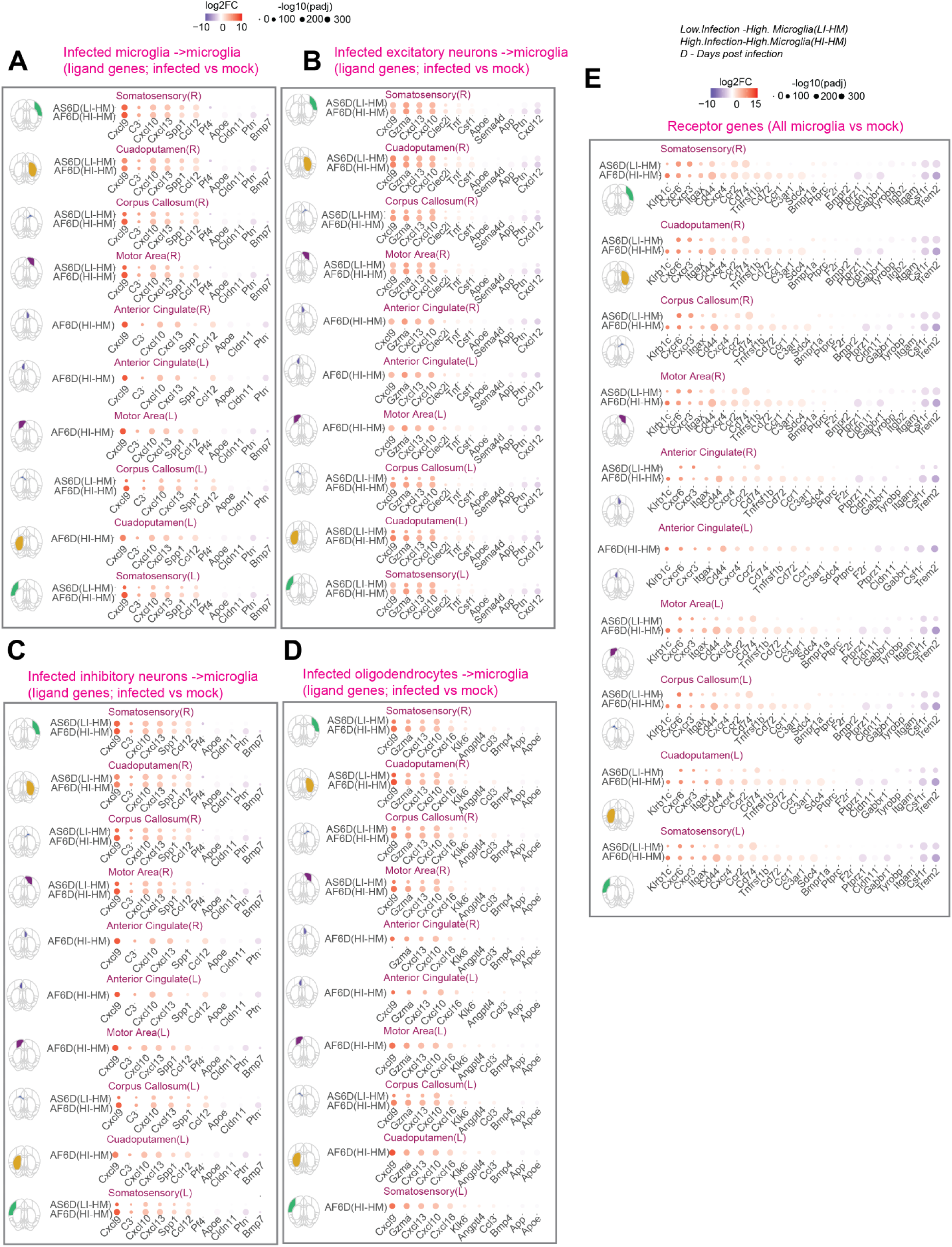
Transcriptional regulation of cell-cell communication pathway components in infection-contained versus highly infected regions. Dot plots showing differential expression of ligand and receptor genes involved in cell-cell communication pathways between infected cells and microglia. Genes shown are those identified in the CellChat analysis (Figure 5D, 5S9) and are significantly differentially expressed in infection-contained regions (Low infection-High microglia; AS6DPI) and highly infected regions (High infection-High microglia; AF6DPI) compared to mock controls within matched brain structures. Dot color indicates average log₂ fold change relative to mock; dot size indicates statistical significance (–log₁₀ adjusted P-value). (A) Ligand genes expressed by infected microglia in the infected microglia → microglia communication pathway. Differential expression assessed by comparing infected microglia versus mock microglia within each structure. (B) Ligand genes expressed by infected excitatory neurons in the infected excitatory neuron → microglia communication pathway. Differential expression assessed by comparing infected excitatory neurons versus mock excitatory neurons. (C) Ligand genes expressed by infected inhibitory neurons in the infected inhibitory neuron → microglia communication pathway. Differential expression assessed by comparing infected inhibitory neurons versus mock inhibitory neurons. (D) Ligand genes expressed by infected oligodendrocytes in the infected oligodendrocyte → microglia communication pathway. Differential expression assessed by comparing infected oligodendrocytes versus mock oligodendrocytes. (E) Receptor genes expressed by microglia (all communication pathways). Differential expression assessed by comparing all microglia in infected samples versus mock microglia. Together, these panels reveal transcriptional regulation of key signaling pathway components, demonstrating that the disruption of Apoe-Trem2 signaling in African strain-infected brains (Figure 5D, main text) is accompanied by coordinated downregulation of both ligand and receptor components, particularly in highly infected regions.

**Figure 6S1.**
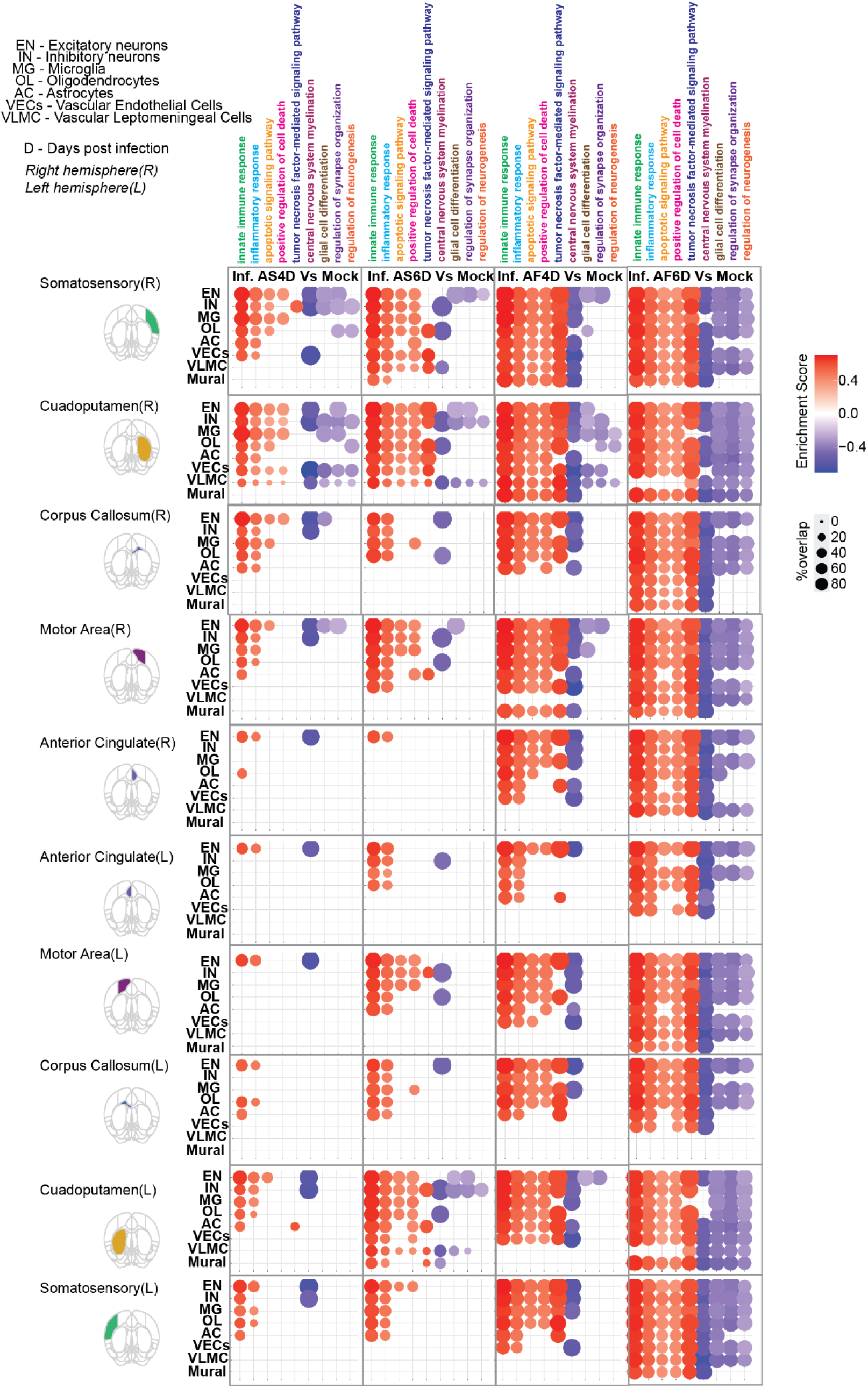
Functional reprogramming of infected structural cell types across brain regions. Bubble plots showing significantly enriched Gene Ontology Biological Process (GO:BP) terms (GSEA, adjusted P-value < 0.05) for infected structural cell types (excitatory neurons, inhibitory neurons, astrocytes, oligodendrocytes) compared to mock controls across major brain structures. Separate comparisons are shown for each condition (Asian 4 dpi, Asian 6 dpi, African 4 dpi, African 6 dpi). Only cell types with ≥30 infected cells per structure are included to ensure statistical robustness. Bubble color indicates enrichment direction (red: upregulated pathways in infected cells; blue: downregulated pathways in infected cells relative to mock); bubble size reflects the proportion of genes within each GO term that are significantly differentially expressed (adjusted P-value < 0.05, |log₂ fold change| ≥ 0.30).

**Figure 6S2.**
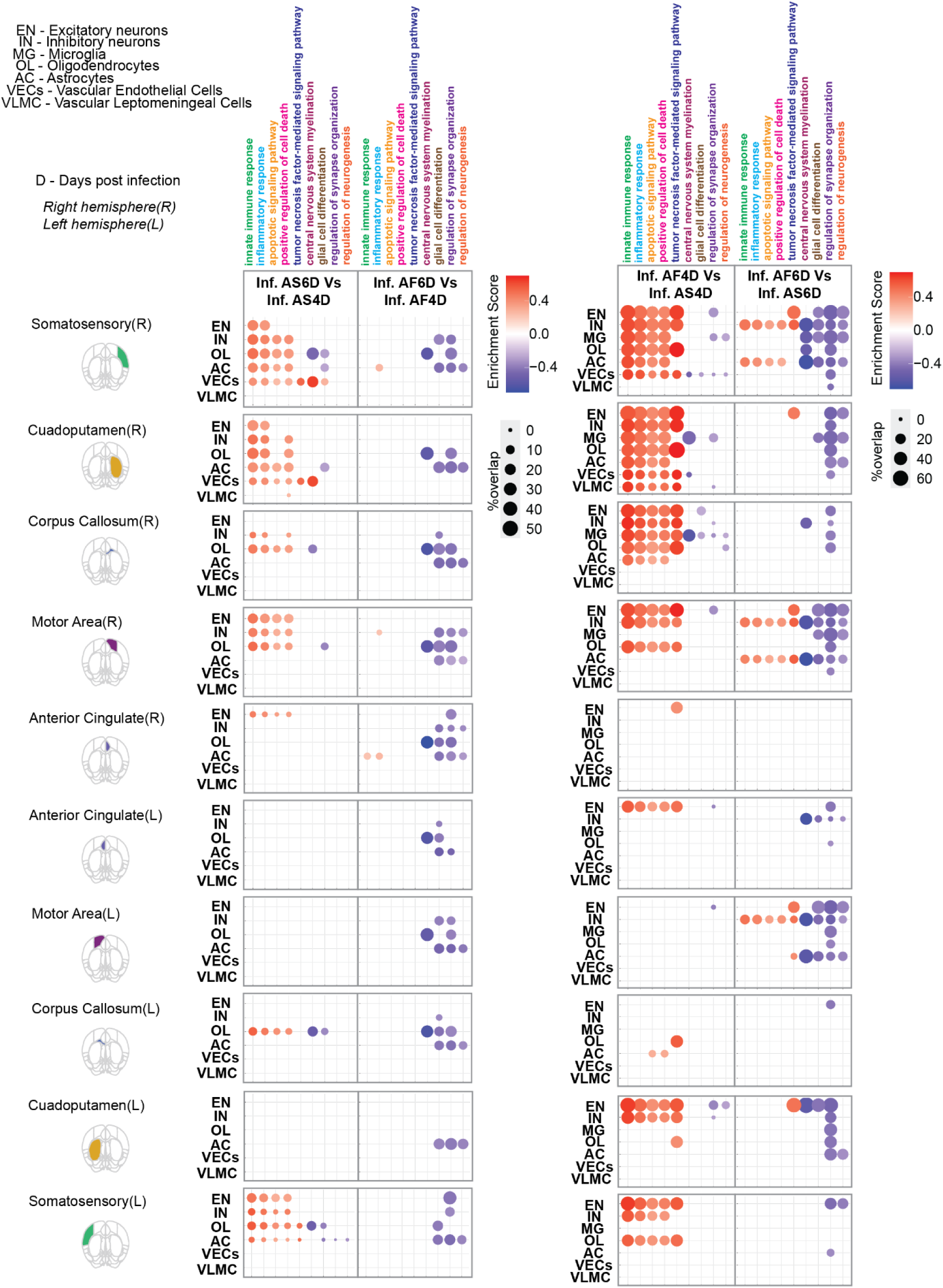
Temporal and strain-specific functional reprogramming of infected structural cell types. Bubble plots showing GO:BP enrichment for infected structural cell types (excitatory neurons, inhibitory neurons, astrocytes, oligodendrocytes) across temporal and strain-specific comparisons. Temporal comparisons: Asian 6 dpi versus Asian 4 dpi; African 6 dpi versus African 4 dpi. Strain-specific comparisons: African versus Asian at matched time points (4 dpi and 6 dpi). Only cell types with ≥30 infected cells per structure are included. Bubble color indicates enrichment direction (red: upregulated in the numerator condition; blue: downregulated in the numerator condition); bubble size reflects the proportion of genes within each GO term that are significantly differentially expressed (adjusted P-value < 0.05, |log₂ fold change| ≥ 0.30).

Video S1. Temporal evolution of simulated infection and microglial dynamics for Asian and African ZIKV strains.

## CORRESPONDANCE AND REQUESTS FOR MATERIALS

Should be addressed to Shou-Jiang Gao or Yufei Huang.

